# A computational framework for designing micron-scale crisscross DNA megastructures

**DOI:** 10.64898/2026.01.23.701435

**Authors:** Matthew Aquilina, Florian Katzmeier, Minke A.D. Nijenhuis, Siyuan Stella Wang, Corey Becker, Yichen Zhao, Su Hyun Seok, Julie Finkel, Huangchen Cui, Jaewon Lee, Seungwoo Lee, William M. Shih

## Abstract

Crisscross polymerization enables the assembly of hundreds of unique DNA origami ‘slats’ into micron-sized structures with nanoscale precision. To design these megastructures, thousands of handle sequences from a fixed library must be assigned to individual slats to encode the desired binding architecture. This complexity presents two major challenges: handles must be selected to minimize parasitic interactions that compete with desired assembly, and the fabrication of hundreds of unique slats creates a substantial logistical burden. Here, we develop a unified framework that standardizes the design and fabrication of crisscross megastructures. We use an evolutionary algorithm to optimize handle assignment and minimize parasitic binding between slats, paired with a graph-based algorithm that expands the handle library. Together, these algorithms enable the assembly of large, multi-layered megastructures that otherwise would be produced at negligible yields. We have released this framework as #-CAD, an open-source graphical application that integrates these algorithms, streamlines laboratory workflows, and makes crisscross DNA origami more broadly accessible.

---

Following its introduction in 2006 [1], DNA origami has matured into a versatile platform for positioning [2], manipulating [3], protecting [4], or encapsulating [5, 6] particles on the nanoscale. This rapid evolution was enabled in large part by dedicated software tools: caDNAno [7] provided the first graphical interface for arranging DNA staple strands on a scaffold, DAEDALUS [8] automated staple routing to generate a variety of three-dimensional shapes, oxDNA [9–12] enabled the simulation of structural stability, and later packages introduced more specialized features such as curved geometries [13], multimeric assemblies [14, 15], and crystal lattice architectures [16]. Today, the vast majority of DNA origami designs are constructed, tested and visualized using one or more of these software tools [17], broadly expanding accessibility.

Beyond nanoscale design, the crisscross paradigm [18] enables the assembly of more than one thousand unique DNA origami monomers into supramolecular structures, effectively extending nanoscale precision to the microscale. These ‘megastructures’, which comprise more than 10^6^ nucleotides [19], reintroduce many of the practical challenges encountered in conventional DNA origami, now magnified by the vastly larger number of components and interactions. Instead of arranging staple strands to fold a scaffold into a three-dimensional shape, a crisscross designer must position hundreds of six-helix bundle building blocks (‘slats’) to construct a three-dimensional megastructure. To weld these slats together, thousands of DNA binding sites (‘assembly handles’) with specific sequences and positions must be assigned. At the same time, unwanted off-target binding events between slats, hereafter referred to as ‘parasitic interactions’, must be avoided. Once a design is completed, the component slats must first be individually folded, then pooled, purified, and assembled. Applying this workflow in practice is a significant logistical undertaking even when assisted by a robotic liquid handler.

In this work, we introduce #-CAD, a computational framework that makes megastructure design more accessible and coordinates the downstream fabrication workflow. Through its graphical interface, users define megastructure geometry and place cargo molecules, while #-CAD generates fabrication-ready outputs, including robotic liquid handler instructions and standardized laboratory protocols. While #-CAD greatly simplifies the design of progressively larger megastructures, increased size leads to a growing number of parasitic interactions among slats that limit assembly yield [19] or prevent assembly altogether. We thus defined a thermodynamics-inspired Loss function that classifies handle assignments by their parasitic interactions, and predicts megastructure completion rates. Using an evolutionary algorithm that optimizes this Loss function, we are able to maximize the yield of fully formed megastructures. In particular, complex multilayer designs previously obstructed by parasitic interactions now achieve practical assembly yields that far exceed those obtained with random handle assignments. The evolutionary algorithm has been fully integrated within #-CAD, which can be applied in just two clicks. In parallel, we doubled the size of our DNA handle library [19] from 32 to 64 sequence pairs using a graph-based selection algorithm, further broadening the design space.

To demonstrate the capability of our new workflow and algorithm, we present a gallery of experimentally realized megastructures that illustrates the range of designs accessible with this framework. The entire software stack has been released as a unified open-source package that runs on a personal computer, simplifying access to customized megastructure design for a wide range of applications. In particular, megastructures are well suited for applications requiring structural customization on length scales comparable to the wavelength of visible light, such as optical metamaterials [20] or artificial antigen-presenting cells [21].

## Principles of Crisscross Assembly and Associated Parasitic Interactions

Megastructures consist of 450 nm-long six-helix bundle building blocks, which we refer to as slats, arranged in a crisscross configuration. At every crossing point, complementary handle sequences hybridize to form 7 bp duplexes that collectively hold the megastructure together. Each slat has two binding faces along its long axis, both presenting 32 regularly-spaced positions for placing programmable assembly handle sequences. These sequences dictate which slats bind together in the megastructure, effectively encoding the final shape and assembly pathway (Fig. 1a).

**Fig. 1:**
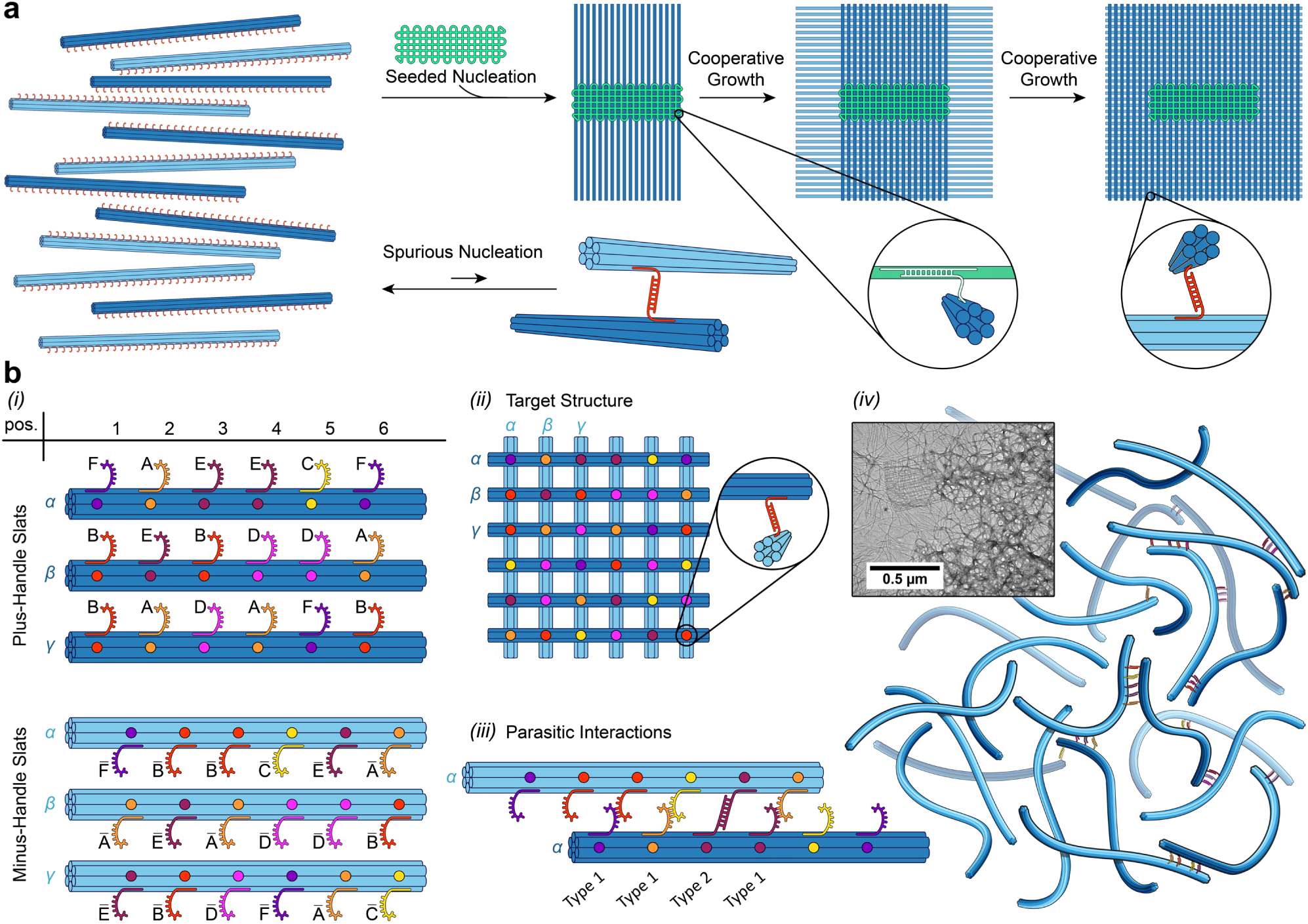
Conceptual overview of megastructure assembly using the crisscross paradigm. **(a)** Schematic illustration of the characteristic all-or-nothing behavior of crisscross assembly. Two pathways are shown. *Lower pathway:* Slats can bind only spuriously at a 90°angle through a single handle, forming weak and unstable complexes. The unbound state is therefore energetically favored, suppressing spontaneous nucleation. *Upper pathway:* A seed recruits several slats and arranges them into a geometry that enables joint-neighbor capture of subsequent slats that attach through multiple handles, leading to rapid completion of the megastructure. The handle binding schemes between the seed and the slats, as well as between slats themselves, are illustrated in the insets. **(b)** Conceptual illustration of the handle library using a reduced set of plus:minus handle pairs and positions. **(i)** Example set of plus-handle bearing and minus-handle bearing slats. Plus-handles and minus-handles are illustrated as colored single-stranded DNA segments labeled with capital letters. Complementary sequences share the same color, and the complement is indicated with a bar (e.g. A and Ā). The arrangement of handles on each slat encodes its position in the target megastructure. **(ii)** Target megastructure where plus:minus handle bonds are indicated by colored circles. The slats shown in (i) can be matched to the target megastructure via color-coded Greek letters. **(iii)** Possible parasitic interactions, in which two slats align in parallel and form unintended bonds. These include bonds between cognate plus:minus handle pairs (Type-2 binding) as well as bonds between non-cognate handles arising from partial complementarity (Type-1 binding). **(iv)** Transmission electron microscopy (TEM) image showing a megastructure adjacent to a large aggregate of slats formed through parasitic interactions, together with a schematic illustrating the parasitic interactions within the aggregate.

Generally, hierarchical assembly approaches produce substantial fractions of partially-completed structures, often requiring multi-step protocols, reducing the yield of fully assembled products with each step [22]. The crisscross strategy mitigates this problem by combining a strong nucleation barrier with seed-mediated cooperative growth, resulting in an effectively all-or-nothing assembly process [18]. As shown in Fig. 1a, slats can encounter each other in the correct crossing geometry but will engage only with a single handle bond. These single-handle contacts have been intentionally designed to be too weak to persist as part of a strategy to minimize spurious nucleation [19]. Productive assembly occurs only when a seed captures several slats simultaneously and positions them so that incoming slats can bind with multiple handles at once. Once this joint-neighbor capture front has been established, coordinated recruitment of successive slats rapidly completes the megastructure (Fig. 1a, upper pathway).

In principle, every intended bond in a megastructure could be encoded by a uniquely designed complementary sequence pair. However, as this would require synthesizing thousands of bespoke staple strands for each new design, such a strategy quickly becomes impractical as structure size increases. Instead, we employ a fixed library of handle sequence pairs from which every bond in a megastructure is selected. In our notation, one handle in each complementary pair is designated as the plus-handle (+), and the other as the minus-handle (-). Fig. 1b (i)-(ii) gives a conceptual overview of the handle library using a set of truncated slats and a reduced set of plus:minus handle sequences. In this work, we maintain 64 handle sequences that can occupy 32 positions on each slat, requiring 64 × 32 unique plus-handle strands that combine each assembly handle with its corresponding scaffold-binding staple sequence. Together with the plus-handle staple strands, a complementary set of 64 × 32 minus-handle staple strands as well as 2 × 32 additional filler staple strands to cover unused binding sites are also required. Apart from the handle library, we also maintain a set of 4 × 32 cargo tag staple strands to attach guest molecules and 10 × 16 seed-binding staple strands to nucleate assembly from a seed origami starter. In total, this yields a library of 4,448 distinct staple strand sequences, which can be selected to customize the two binding faces of every slat in a megastructure.

Because all slats draw from the same finite assembly handle pool, they are prone to unwanted parasitic interactions. Two slats can align in parallel in an arrangement that allows their handle sequences to form unintended bonds (Fig. 1b (iii)). These parasitic interactions are composed of weak partially complementary bonds between noncognate sequences (Type-1 binding), and fully cognate plus:minus handle bonds (Type-2 binding). Such interactions can drive the formation of aggregates (Fig. 1 (iv)) in competition with the desired target megastructure, thereby limiting yield and slowing down assembly of the latter. To mitigate these parasitic interactions, we employ two aforementioned sequence-level strategies. Our evolutionary algorithm strategically assigns handle sequences to slats such that the number of unintended Type-2 bonds between two slats are reduced. Simultaneously, the expansion of the handle library from 32 to 64 sequence pairs further reduces the likelihood of unintended Type-2 bonds. The 64 sequences were selected using a graph-based algorithm in order to weaken Type-1 partial-complementarity interactions between noncognate sequences. Both approaches are described in detail in later sections.

## Megastructure Fabrication Pipeline

Our approach to megastructure production is centered around a fabrication pipeline that borrows elements from the 3D-printing workflow (Fig. 2). A megastructure is designed in CAD software, the handle arrangement is algorithmically optimized, the finalized design is exported as machine-readable instructions, and a liquid handler dispenses the required staple strands from our fixed handle library (‘ink’) into reaction vessels for subsequent manual processing. Importantly, because all handles are selected exclusively from this fixed library, new megastructure designs do not require ordering custom DNA strands.

**Fig. 2:**
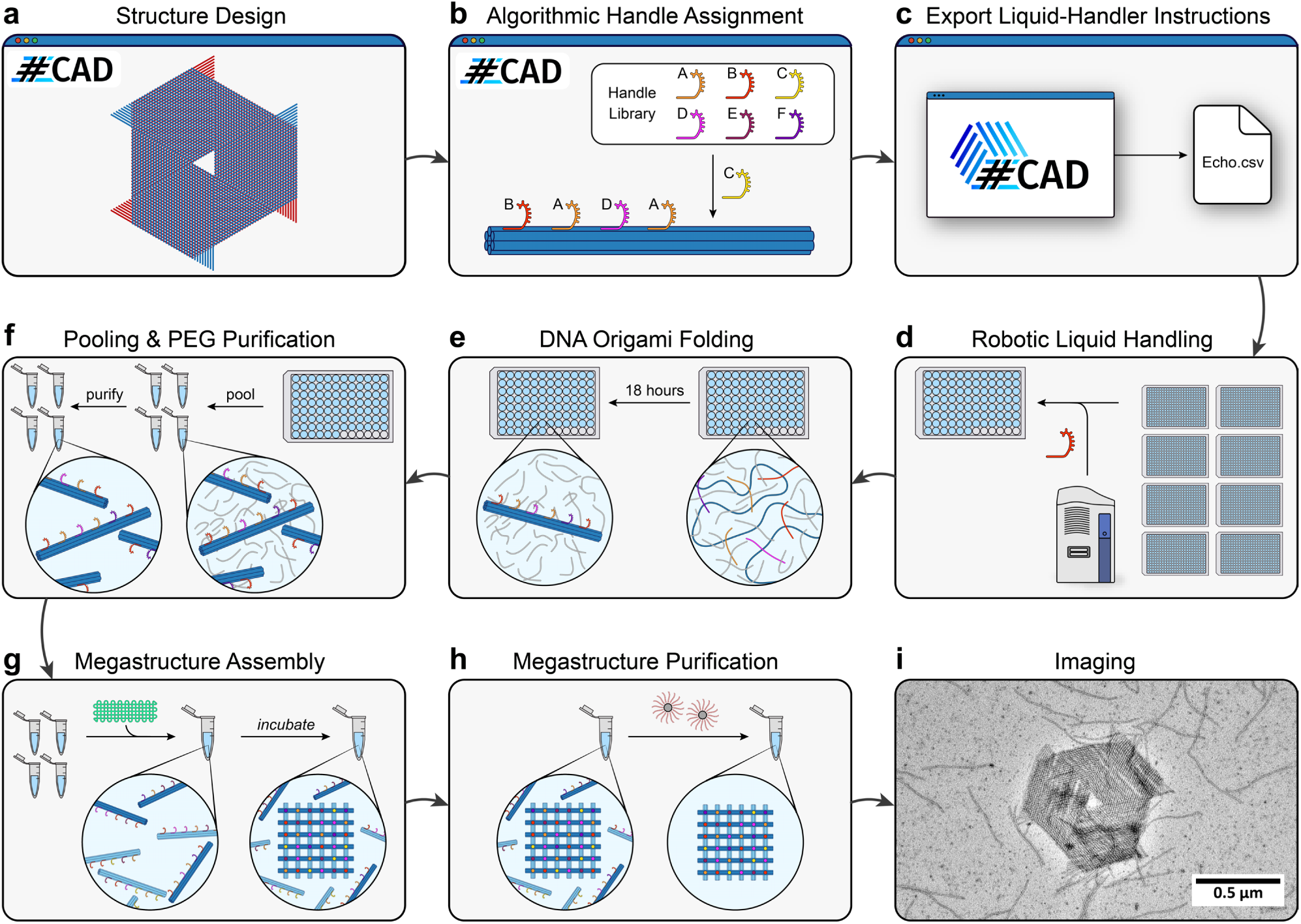
Step-by-step workflow for assembling crisscross DNA origami structures. **(a)** An example megastructure comprised of 192 slats is designed using our open-source software #-CAD. **(b)** Plus-handles and minus-handles are algorithmically assigned to encode the megastructure geometry while minimizing parasitic interactions. **(c)** The optimized design is exported as liquid-handling instructions, requiring over 12,000 individual transfers for this design. **(d)** A robotic liquid handler transfers plus and minus handles from 384-well source plates into 96-well target plates, where each well corresponds to a single slat. **(e)** Scaffold DNA, structural staple strands, and folding buffer are added, and the slats are folded by thermal cycling. **(f)** Folded slats are pooled into groups for easier handling and purified by PEG precipitation to remove excess staple strands; for a 192-slat design, this corresponds to twelve groups of sixteen. **(g)** Purified slat groups are combined with seed DNA origami and assembly buffer, and incubated at 37 °C for several days to form the megastructure. **(h)** Assembled megastructures are purified using magnetic beads to remove excess slats. **(i)** Structure formation is verified by TEM (or other microscopy). The displayed image corresponds to the design shown in panel a, which we refer to as the ‘Shuriken’.

We developed #-CAD to provide control over this workflow (Fig. 3). At the design stage, the graphical interface provides tools to construct megastructures with diverse architectures. Slats can be stacked into two or more layers, enabling 2D or 3D designs. Furthermore, slats can be arranged on square or triangular grids with 90°or 60°connection angles, giving rise to distinct structural motifs (Fig. 4). In addition, automatic 3D rendering of the megastructure provides real-time visualization of design progress. Once a design is completed, the evolutionary algorithm can be run directly within the software through a simple graphical interface with built-in real-time monitoring.

**Fig. 3:**
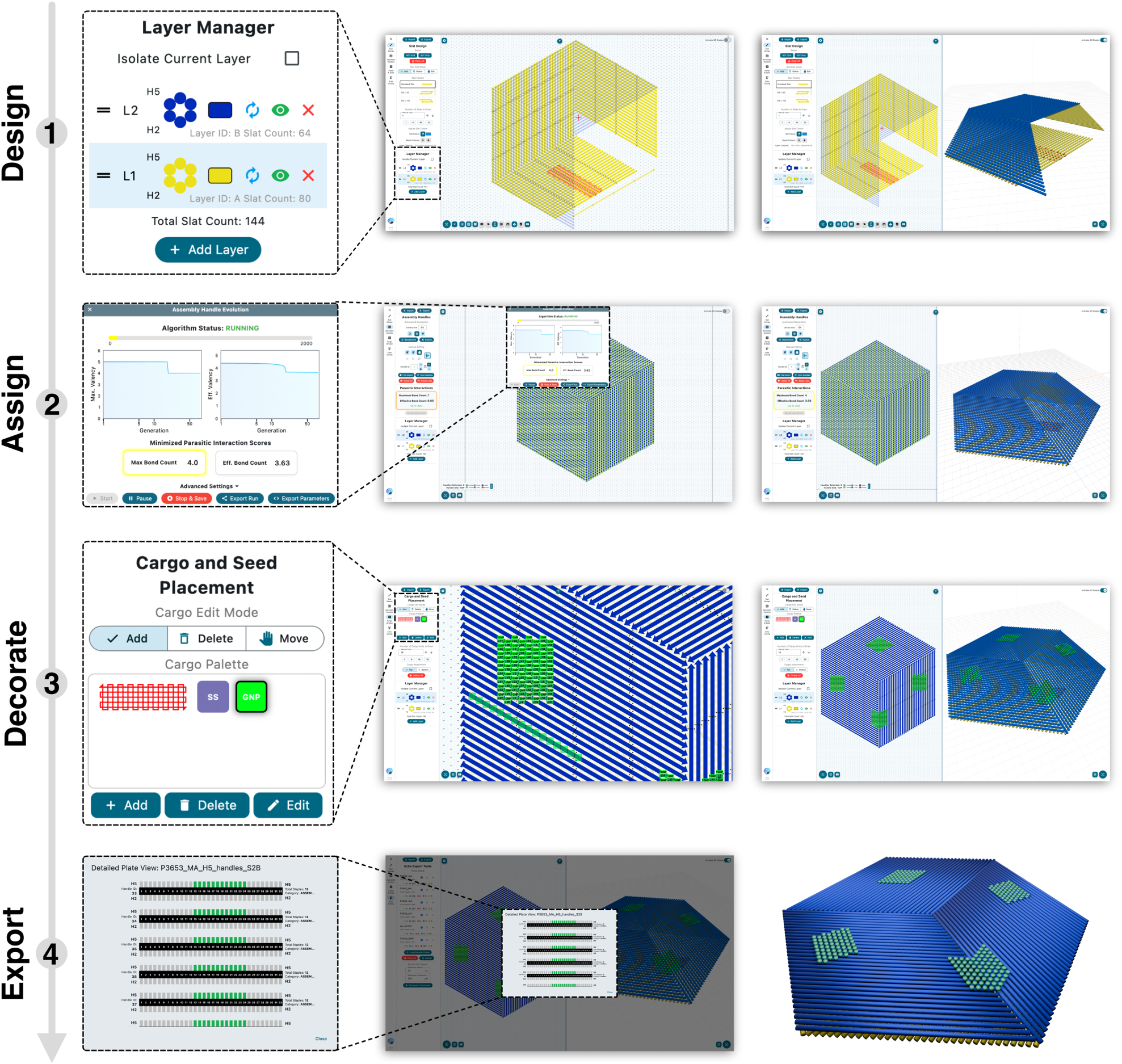
#-CAD design software workflow. Step 1: **Design**. Slats are placed by the user onto a design canvas and assigned to different layers. The central panel displays a partially completed design, with a zoomed-out view of the layer manager highlighted, allowing selection of the layer into which slats are placed. Slats can be organized with 90°or 60° binding angles; the example shown uses a 60°geometry. The right panel shows the full interface that includes an interactive 3D rendering of the current structure. Step 2: **Assign**. Handles are assigned to the layer interfaces using the evolutionary algorithm integrated into #-CAD and accessed through a graphical interface requiring minimal user interaction. The interface provides a live view of the optimization progress, including a real-time plot. Step 3: **Decorate**. Final design adjustments are made, including selection of cargo positions and specification of the seed location that defines the initiation site of megastructure assembly. In the example shown, four cargo patches have been placed, with green selected as their display color. Step 4: **Export**. A .csv file is automatically generated for pooling slat handles using a robotic liquid handler. A final confirmation view (left) displays the availability of handle sequences in the storage plates to prevent errors. Additional export options include a Blender-compatible 3D representation (right), plate layouts for laboratory workflows and pipetting instructions. Complete #-CAD designs can be saved into Excel format, with separate sheets for slat positions, handle identities and cargo tags, allowing for convenient manual editing.

**Fig. 4:**
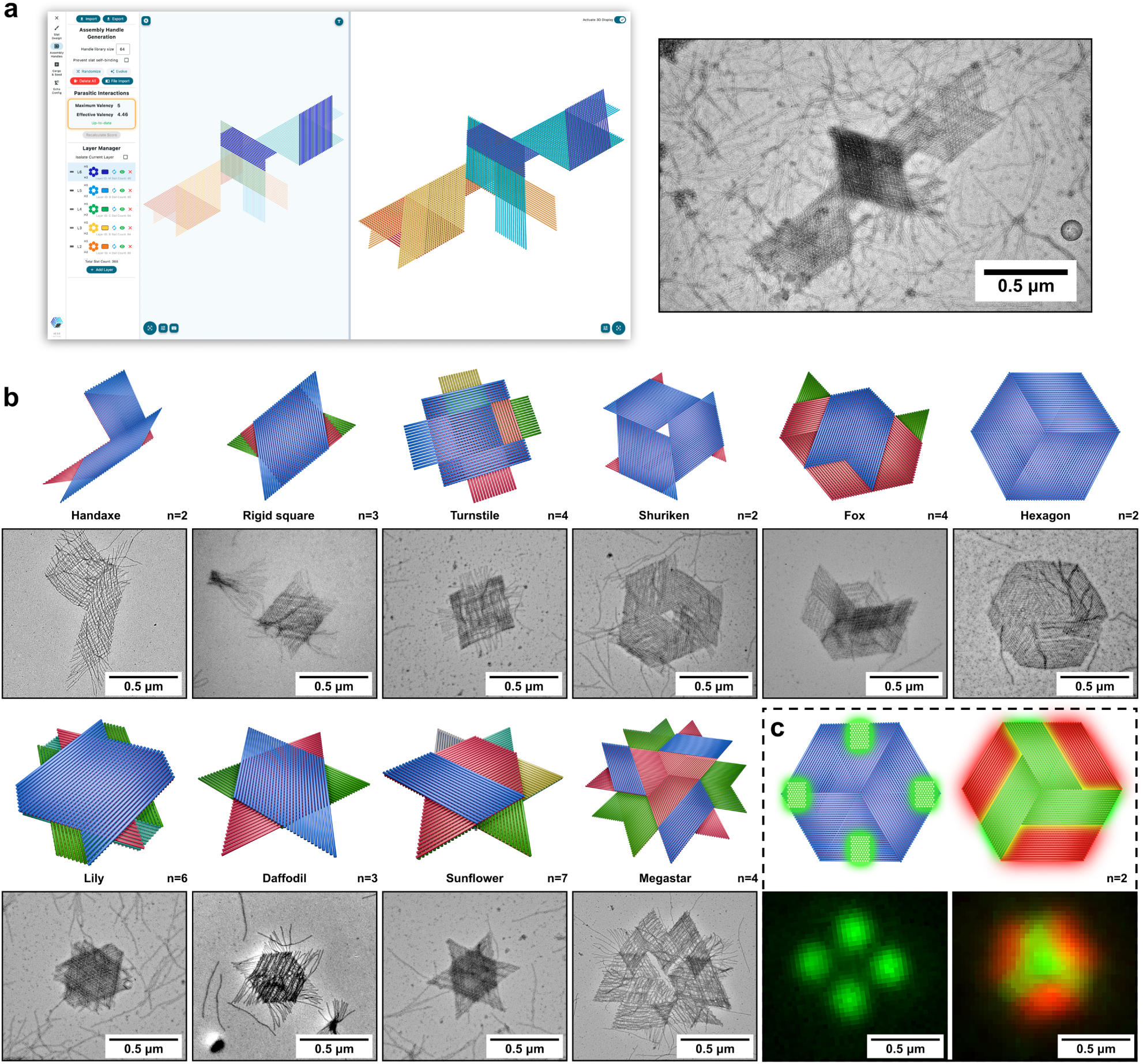
Megastructure design gallery. **a** Showcase of our largest multi-layered design to-date: a 7-layered ‘Bird’ megastructure with 368 unique slats and a wingspan of approximately 2 µm. The left panel shows the design in #-CAD while the right panel shows a negatively-stained TEM image of a single particle. The core has a darker tone due to the larger number of layers stacked on top of each other. **b** A large variety of designs illustrating different strategies for constructing unique structural motifs. For each design, the top panel shows a 3D schematic of the structure while the bottom panel contains a TEM image of a single particle. From left to right, top to bottom: the ‘Handaxe’, ‘Rigid square’, ‘Turnstile’, ‘Shuriken’, ‘Fox’, ‘Hexagon’, ‘Lily’, ‘Daffodil’, ‘Sunflower’ and ‘Megastar’. **n** indicates the number of slat layers in each design. **c** Two Hexagon megastructures modified to include fluorescent elements. The left design bears DNA cargo handles on its top layer that can be targeted by fluorescent DNA nanocubes [23]. The right design contains slats that have been modified to include fluorophores in their core staple strands. The bottom panels show fluorescent microscope images of a single particle, illustrating that super-resolution is no longer necessary to identify megastructures or their cargo at this scale. Larger field-of-view images for all structures are provided in the Supplementary Information.

To complete the megastructure design, users can specify the location of the seed origami, which determines the slats that initiate megastructure assembly. Optional cargo dock handles for attaching guest molecules, such as other DNA origami nanostructures or fluorophores, can be added directly to the design. Multiple visual customization options and predefined template patterns are available to allow efficient cargo placement and distribution.

Finally, #-CAD allows users to load custom plate layouts defining the storage locations of handle sequences. This enables the software to generate liquid handling commands for an Echo 525 liquid handler, prepare manual pipetting instructions for slat folding and PEG precipitation, and export the design to common 3D graphical formats for visualization and collaborative use.

To demonstrate the versatility of our approach and the design freedom afforded by #-CAD, we designed and fabricated a wide range of megastructures using our pipeline (Fig. 4). The megastructures selected range from simpler designs intended to explore the impact of specific structural motifs (Fig. 4b, ‘Handaxe’, ‘Shuriken’, etc.) to 2 µm diameter multi-layer showcases (Fig. 4a).

In standard 90° designs, assembly handle bonds between layers allow relative shear between slats in neighboring layers, resulting in flexible megastructures in which the intended 90°slat angles are not always conserved. This shear can be partially reduced by increasing the number of stacked layers (‘Turnstile’), but is more robustly suppressed in 60° designs. In 60° architectures, slats are stacked in three orientations separated by 60° angles, producing triangulated structures that suppress shear and yield rigid geometries conserved across most particles (e.g. Daffodil and Sunflower). These designs enable structural motifs that are inaccessible in 90°geometries, including finite rigid two-layer sheets (Hexagon), motifs featuring defined pores (Shuriken), and more complex shapes such as the Bird shown in Fig. 4a.

However, 60°designs can be more challenging to fabricate. Ambiguous slat growth pathways can reduce yield (e.g. the Handaxe, Supp. Fig. 2), and achieving full triangulation can act as a kinetic bottleneck during assembly, as explored in more detail for the Sunflower design below. Conversely, large multi-layer structures designed without sufficient support for seams, such as the Megastar, frequently exhibit defects and tearing (Fig. 4b).

Furthermore, given their micrometer scale, megastructures are large enough to be visualized directly using fluorescence microscopy without any super-resolution techniques. In Fig. 4c, we configured the Hexagon to contain cargo handles that can bind fluorescent nanocubes (Method 16) in four distinct patches, or to incorporate slats containing oligonucleotide-conjugated fluorophores (Method 15). We were able to identify the designed structures directly from their fluorescent cargo, demonstrating that megastructures can be characterized without relying on the more niche transmission electron microscopy (TEM) and atomic force microscopy (AFM) common to DNA origami literature. Taken together, these examples illustrate the design freedom afforded by our #-CAD controlled fabrication pipeline, enabling the prototyping, testing, and iteration of new and more complex megastructures with minimal friction.

## Evolutionary Algorithm: Classification of Handle Assignments

The effectiveness of our pipeline demonstrated in Fig. 4 was largely enabled by the reduction of Type-2 parasitic interactions (Fig. 1b) using our evolutionary algorithm. As the algorithm’s objective function, we defined a ‘Loss’ value that classifies handle assignments based on Type-2 parasitic interactions. Lower Loss values correspond to better handle assignments with fewer and consequently weaker parasitic interactions. The Loss is defined, together with an explanatory illustration for a square-shaped megastructure, in Fig. 5a. To compute the Loss for a given handle assignment, all possible pairings of plus-handle bearing and minushandle bearing slats are computationally screened for potential parasitic interactions. In the example shown, this corresponds to all pairings of light-blue minus-handle bearing and dark-blue plus-handle bearing slats. Each of these combinations can adopt multiple relative binding configurations, of which a subset is illustrated for one example combination in Fig. 5a.

**Fig. 5:**
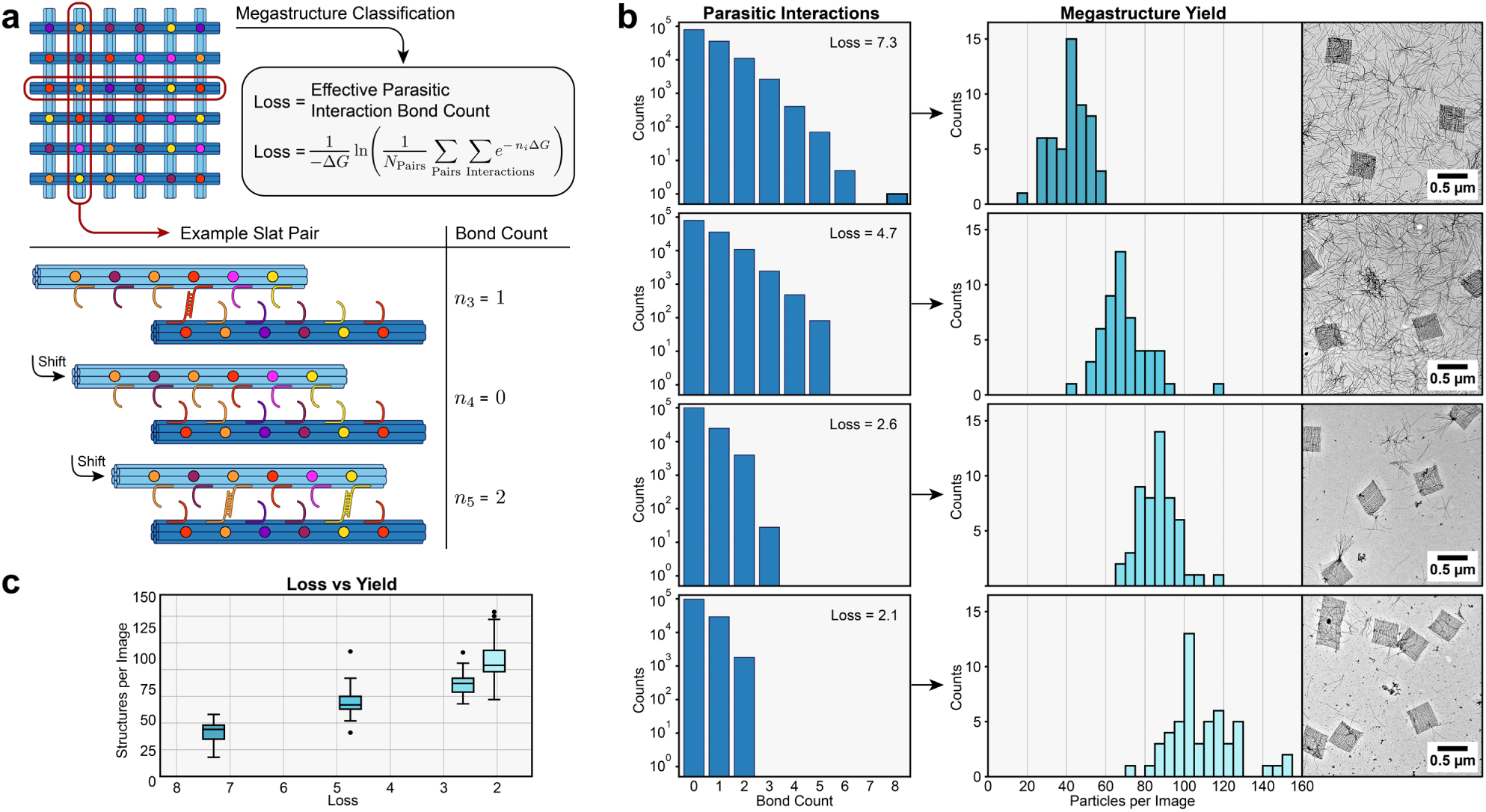
Computation of the Loss and its effect on assembly yield. **a** Explanatory illustration of the Loss computation for a square-shaped megastructure, with a reduced set of positions and color-coded plus:minus handle pairs. One example slat combination is outlined in red, and several of its binding configurations are shown in a table. The bond count for each configuration is listed in the right column. The computational strategy to generate these configurations is indicated by arrows labeled ‘Shift’. **b** Experimental comparison of the assembly yield of four square-shaped megastructures designed with distinct handle assignments and Loss values. The left column shows histograms of the corresponding parasitic interaction bond counts between the slats in each design. The right column shows representative TEM images and yield histograms obtained by counting complete megastructure particles from 50 TEM images per sample. **c** Box plot of the histogram data in panel b, summarizing the relationship between assembly yield and Loss. Boxes indicate the interquartile range (IQR). Whiskers extend to 1.5 × IQR. Black lines mark medians, and black points represent outliers.

Computationally, these configurations are systematically generated by sliding one slat relative to the other. Then, for each configuration, we determine the bond count as the number of plus-handles aligned with their complementary minus-handle counterparts. Notably, shifting one slat, as illustrated in Fig. 5a, is not sufficient to generate all possible binding configurations between two slats. Because slats are three-dimensional objects, binding can also occur when one slat is rotated end-to-end, reversing the handle order. The shift operation must therefore be applied again to this reversed configuration, yielding an additional set of possible binding configurations. From all the recorded bond counts, we compute the Loss as a thermodynamically weighted average, which we interpret as an effective parasitic interaction bond count. With this interpretation, all plus-handle bearing slats bind to minus-handle bearing slats with a number of bonds equal to the Loss. The derivation of this property and further details are provided in Supplementary Note 1.

In Fig. 5b, we present experimental results showing how parasitic interactions influence the yield of fully formed megastructures in an assembly reaction. We assembled four square-shaped (‘Square’) megastructures with different handle assignments covering a range of Loss values (Method 18). A histogram of the number of parasitic interactions per bond count for each handle assignment is shown in the left column of Fig. 5b. To quantify the assembly yield, we recorded 50 TEM images for each handle assignment and counted the number of fully assembled megastructure particles. The number of completed megastructures per image, together with representative TEM images, are provided in the right column of Fig. 5b. Samples with lower Loss values yielded more fully assembled megastructures, as reflected by the rightward shift of the corresponding histograms. The overall increase in assembly yield with decreasing Loss is summarized in the box plot in Fig. 5c. Additionally, the TEM images of the two squares with lower Loss value showed fewer unbound slats after purification, suggesting that lower Loss also enable more efficient purification. We hypothesize that during magnetic bead purification, slat aggregates driven by parasitic interactions trap megastructures, and are consequently pulled down together with the magnetic beads, resulting in free slats carrying over into the final purified solution.

Finally, an experiment comparing the yield of the two megastructures with Losses 7.3 and 2.6 over time confirmed that the results obtained are representative of the final yield after assembly is complete (Extended Data Fig. 1). Overall, these analyses show that parasitic interactions inhibit megastructure formation and complicate purification, with the Loss function serving as a reliable predictor of assembly efficiency.

## Evolutionary Algorithm: Optimization of Handle Assignments

The experiments of Fig. 5 confirm the Loss as an effective measure of the quality of a megastructure’s handle assignment. To minimize this Loss, we adopted a classic evolutionary algorithm [24] that iteratively refines handle assignments to minimize parasitic interactions. The workflow of this algorithm is shown schematically in Fig. 6a. Starting from a population of random assignments, the algorithm evaluates designs by their Loss values, retains the best-performing ones, and generates new candidates from these survivors through random handle swaps. Repeating this process over successive generations progressively reduces parasitic interactions and yields optimized handle assignments with minimal Loss values.

**Fig. 6:**
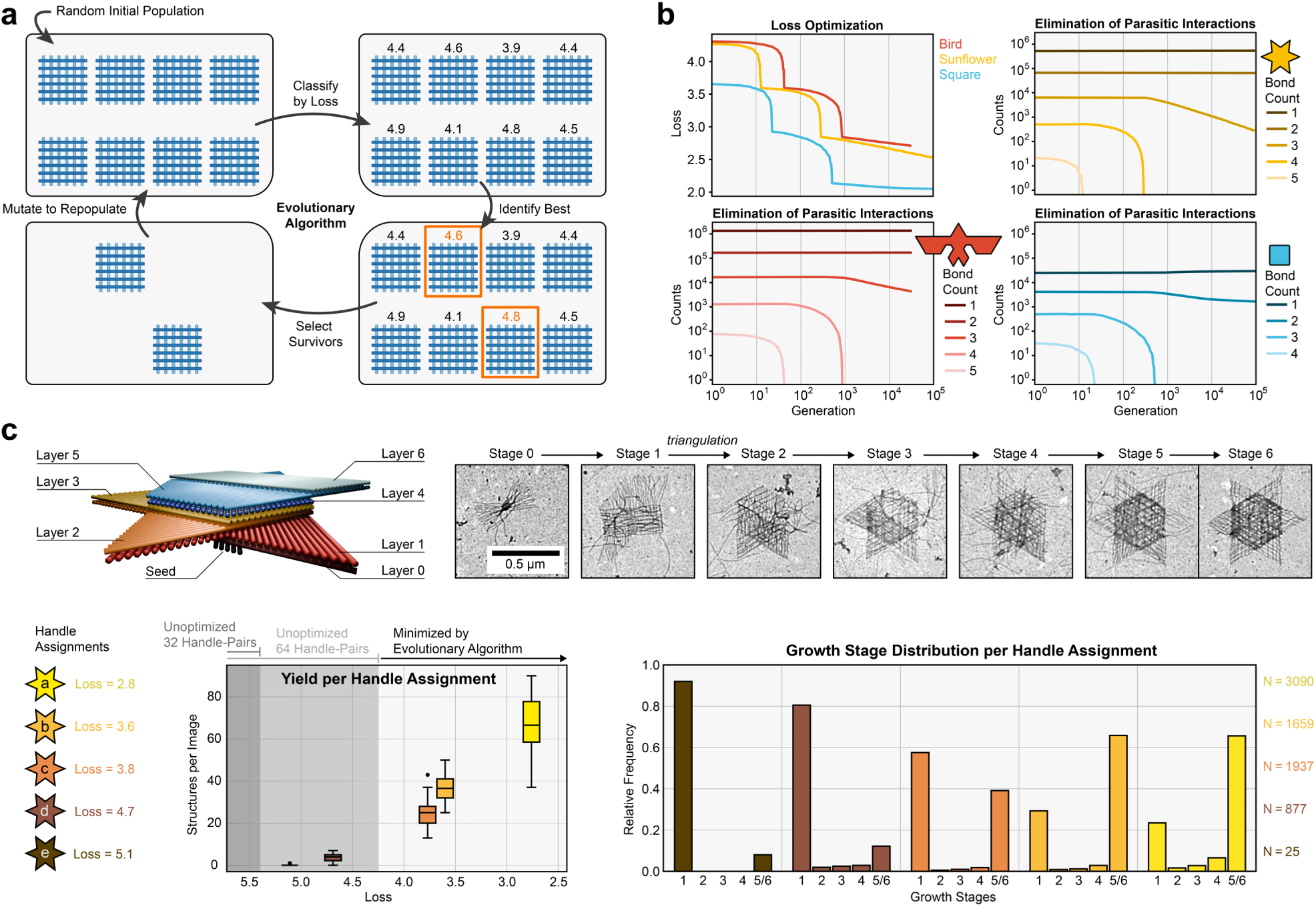
Evolutionary optimization of handle assignments. **a** Schematic illustration of the evolutionary algorithm. In brief, we initialize a population of megastructures with random handle assignments. Next, we compute the Loss for each assignment (example Loss values are shown above the squares). Finally, we select the designs with the lowest Losses (orange outlines) and discard all others. The selected designs are used as templates for the next generation, which are mutated through random handle swaps. **b** Example runs of the evolutionary algorithm for the Square (64 slats), Sunflower (192 slats), and Bird (368 slats) designs. The upper-left plot shows the evolution of the Loss of the best candidate for all three structures. The remaining plots track, for each structure, how the number of parasitic interactions changes over generations, with separate lines indicating different interaction bond counts. The evolutionary algorithm generation is labeled in powers of 10 (from 1 to 100,000) on the x-axis on the lower plots. **c** Dependence of megastructure yield and structural completeness on the Loss for the multilayer Sunflower design. The 3D rendering (top left) provides a side view of the Sunflower with labeled layers. The TEM image sequence (top right) shows the identifiable growth stages, where each stage corresponds to a layer in the 3D rendering. Stage 1 is a square lattice, while stage 2 and onward contain triangulated layers. Stages 5 and 6 are visually almost indistinguishable and were grouped together for analyses. The box plot (bottom left) reports the number of fully assembled megastructures (stages 5 and 6) detected by TEM for each handle assignment at different Loss values. Each box summarizes 30 TEM images. Boxes indicate the IQR, whiskers span 1.5× IQR, black lines mark medians, and black points denote outliers. Grey regions mark the Loss ranges achievable by randomly generating handle assignments without using our algorithm. Dark grey and light grey correspond to what is achievable with 32 and 64 handle pairs, respectively. Theses boundaries were computed by generating 10,000 random assignments for each handle library size. To obtain a practical yield, handle assignments need to have a Loss below a certain threshold, which cannot be achieved for the Sunflower design without our evolutionary algorithm. The bar graph (bottom right) shows the normalized distribution of growth stages achieved for each Loss, with total structure counts indicated next to the plot. Completed structures are enriched at lower Loss values.

In Fig. 6b we present representative runs of the evolutionary algorithm for the Square (64 slats), Sunflower (192 slats), and Bird (368 slats) designs, showing the evolution of the Loss function and the elimination of parasitic interactions separated by bond count for each design (Method 5). Across all cases, the Loss decreases over successive generations, with smaller structures reaching lower final Losses due to the reduced number of possible slat combinations. The evolution of parasitic interactions reveals that high bond count interactions are eliminated first, followed by progressive elimination of lower bond count ones. The sudden drops in the Loss function correspond to generations in which an entire bond count class of parasitic interactions is completely eliminated. Further details on adjustable parameters such as mutation rate, population size, and their influence on algorithmic performance are provided in Method 4 and Extended Data Fig. 2/ Extended Data Fig. 3.

To demonstrate how the evolutionary algorithm enables the design of complex megastructures, we investigated the relationship between Loss, assembly yield, and structural completeness for the multilayer Sunflower design (Method 20). The Sunflower was designed to have assembly stages that are easily distinguishable, as shown in Fig. 6c. We generated five distinct Sunflower handle assignments with different Losses, assembled the corresponding megastructures, purified the samples, and recorded 30 TEM images per sample. Each observed particle was counted and classified according to the growth stage it had reached. The total number of completed structures per image was then plotted against the corresponding Loss (Fig. 6c, lower left box plot). The results reveal a pronounced dependence of yield on Loss: designs with the highest Loss produced almost no complete structures (most TEM images were empty), whereas the lowest Loss designs yielded a median of approximately 65 completed structures per image. To place these results in context, we compared the Losses of our designs with those obtained from randomly generated handle assignments (grey regions in Fig. 6c). The two worst Losses among our tested designs fell within the range possible with a random assignment of 64 handle-pairs (light grey region), while a random assignment of just 32 handle-pairs would have exhibited even higher Losses (dark grey region). This comparison highlights that the evolutionary algorithm is essential for the successful assembly of complex multilayered megastructures.

Apart from the global yield analysis, we analyzed structural completeness for each handle assignment (Fig. 6c, lower right plot). The fraction of completed structures increased with better Losses, following the same trend observed for overall yield. Interestingly, for each design, most structures were found either in growth stage 1 or fully completed, resulting in a bimodal distribution that suggests a kinetic barrier after stage 1. We hypothesize that this barrier coincides with the onset of triangulation during growth i.e. the point at which the structure must rigidify into a triangular grid to allow additional slats to bind and continue assembly.

## Expanded Handle Library

Another solution to minimizing the Loss is to use a larger handle library. With more available sequences, the probability that a parasitic interaction between two slats involves a fully complementary plus:minus handle bond decreases (Type-2 binding; Fig. 1b). However, larger handle libraries also increase the likelihood of partial complementarity between non-cognate plus-handles and minus-handles (Type-1 binding; Fig. 1b). The ideal would therefore be a large pool of orthogonal plus:minus handle pairs, in which each sequence strongly binds only to its cognate complement, while all other interactions remain weak.

We developed a graph theory based algorithm to identify an expanded set of 64 orthogonal sequence pairs. We determine sequence orthogonality by computing binding energies using NUPACK thermodynamic models [25–27] (Method 21). The initial steps of the sequence search algorithm are illustrated in Fig. 7a. In brief, 7-mer sequences are first filtered as potential candidates and their on- and off-target binding energies are pre-computed. As part of this process, a combination of two candidate pairs needs to be validated as being compatible. This is done by ensuring that all off-target interactions between the four sequences involved have binding energies above a predefined cutoff.

**Fig. 7:**
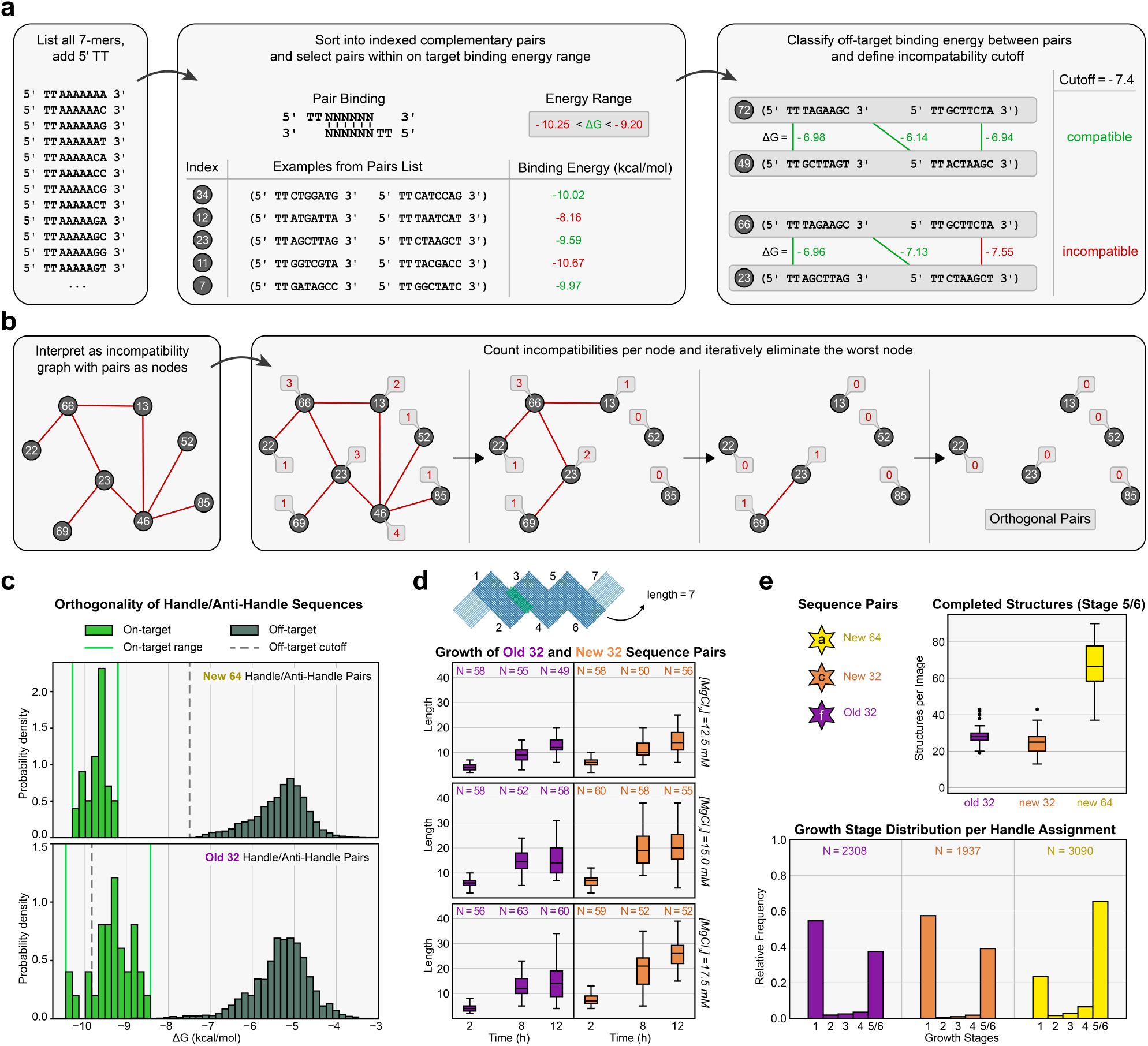
Graph based algorithm to find orthogonal sequences. **a** Preparation steps for the sequence search algorithm. Left: All 7-mer DNA sequences with an added 5*^′^* ‘TT’ are listed. Middle: Complementary sequences are paired, assigned numeric indices, and annotated with their binding energy (Δ*G*). Pairs outside the on-target energy range (red values) are removed before the next stage. Right: Pair combinations are evaluated for off-target interactions. Combinations for which all off-target interactions are above the cutoff (green values, upper example) are classified as compatible, whereas those containing at least one interaction below the cutoff (red values, lower example) are incompatible. **b** Graph-based sequence search. Left: Example illustrating how a list of incompatible sequence-pair combinations is represented as a graph. Sequence pairs appear as nodes labeled with their numeric indices, and edges indicate incompatibility between the corresponding pairs. Right: Example of the iterative elimination procedure used to obtain an orthogonal set. At each iteration, nodes are labeled with the number of incompatibilities they have, and the node with the highest count is removed. **c** Energy distributions for the new 64-pair library and the previously used 32-pair library [19]. **d** Comparison of crisscross polymer growth for a periodic, zig-zag-shaped design using the old and new handle libraries. The top illustration shows the zig-zag polymer with labeled corners, indicating how polymer length was measured. The box plots display the length distributions of polymers assembled with the old 32-pair library or the new 32-pair subset at different magnesium concentrations and sampling times. The number of counted structures is provided above each box. Boxes indicate the IQR; whiskers extend to 1.5×IQR. Black lines mark medians, and black points represent outliers. **e** Dependence of yield and structural completeness on the handle library for the multilayer Sunflower design of Fig. 6. Assembly was performed using the same experimental conditions as in Fig. 6c. The old 32-pair handle library is compared to a restricted new 32-pair subset and the full new 64-pair library. The upper box plots show the number of completed multilayer megastructures (growth stages 5–6) identified in TEM images. The lower bar plots show the corresponding distributions of growth stages for each library. Data for the new 32-pair and new 64-pair libraries correspond to handle assignments with Losses of 3.8 and 2.8, respectively (Fig. 6c). The old 32-pair library uses the same handle assignment as the new 32-pair subset and therefore has the same Loss. The old 32-pair and new 32-pair libraries show comparable performance, whereas the 64-pair library yields a higher fraction of completed structures and increased overall yield.

To identify an orthogonal subset from the screened sequence pairs, we interpret the list of incompatible pair combinations as a graph in which each node represents a sequence pair and each edge denotes an incompatibility [28]. A visual example of this graph representation is shown in Fig. 7b. In graph terminology, the search for orthogonal sequence pairs corresponds to finding a maximum independent set, which is an NP-hard problem. We therefore use a heuristic, iterative pruning procedure [29, 30]. In each iteration, we remove the node with the highest number of incompatibilities and then recalculate the incompatibility counts of the remaining nodes. Repeating this procedure progressively eliminates sequence pairs involved in many incompatibilities. Once all edges are removed, the remaining nodes form an edgeless subgraph, corresponding to an orthogonal set of sequence pairs that can coexist in the same handle library. The full algorithmic workflow, including tie-breaking strategies, is detailed in Method 21.

The distributions of on-target and off-target binding energies for the new library identified using this method are shown in Fig. 7c, together with the corresponding distributions for the 32-pair library used in the first crisscross megastructure prototypes [19]. Each plot displays the on-target binding energies of the cognate pairs as well as the off-target energies arising from non-cognate interactions within the respective pools. The allowed on-target energy window and the off-target cutoff are indicated for both libraries. Notably, in the previously used sequence set, the off-target energy distribution extends into the on-target energy range due to several strong off-target interactions. In contrast, the newly designed 64-pair library exhibits a clear separation between on-target and off-target energies, with a pronounced gap between the two distributions. This gap reflects the improved orthogonality achieved by our thermodynamic and graph-based selection strategy.

We experimentally evaluated 32 handle pairs from the new sequence library using a zig-zag shaped crisscross polymer. This structure consists of repeating units formed by two groups of 16 slats that grow in discrete steps, enabling its length to be measured by counting the number of corners in each polymer (Fig. 7d, top). Assembly reactions were set up with either the old 32-pair library or the new 32-pair subset at three magnesium concentrations (12.5 mM, 15 mM, and 17.5 mM; Method 22). Samples were collected after 2 h, 8 h, and 12 h of incubation. Crisscross polymers were imaged by scanning a TEM grid and counting the number of corners in each structure encountered. The resulting length distributions are shown in Fig. 7c. Both libraries produced similar polymer lengths across most conditions. Overall, the two libraries performed comparably when considering the same number of sequence pairs.

To compare the two libraries on a megastructure with optimized Loss, we assembled and characterized an additional Sunflower (Fig. 6c) with Loss 3.8 using the old 32-pair library. We compared both the quantity of completed megastructures and the growth stage reached for Sunflowers assembled using both new and old handle libraries (Fig. 7e). The old 32-pair library and the new 32-pair subset again performed similarly, showing nearly identical growth-stage distributions and a comparable number of completed megastructures. As expected, the larger 64-pair library produced roughly twice as many completed structures and exhibited a substantially larger fraction of fully assembled particles.

The results indicate that when comparing the same number of sequence pairs, the two libraries appear to perform similarly in terms of megastructure growth speed and yield. Our strategy for selecting handle sequences was clearly effective, but the impact of highly separated on-target and off-target binding energies appears to be less important for crisscross assembly. It is likely that a certain threshold of on-target/off-target separation is still required, which our library selection strategy can guarantee. Nevertheless, expanding to 64 sequence pairs allows for improved performance by enabling the evolutionary algorithm to minimize the Loss even further. This ‘new’ library was used to create all the megastructures described in this work.

## Conclusions

Throughout this work, we developed and validated a complete pipeline for designing and fabricating crisscross megastructures. We based our framework on #-CAD, a graphical software package that allows a user to design and visualize a megastructure *ab initio*. Its interactive features together with its real-time 3D view enable rapid prototyping of new designs. Alongside #-CAD, we advanced the capabilities of crisscross through a set of computational approaches to improve the yield of fully-formed megastructures. Our evolutionary algorithm, which is directly integrated into #-CAD, can be used to minimize parasitic interactions that otherwise significantly hinder assembly. The algorithm has unlocked the fabrication of complex, multi-layer designs that were unachievable using the prior version of our design framework. This enabled us to create a gallery of new structural motifs using both 90°and 60°slat connections, with the 60°system yielding more rigid and intricate megastructures. Finally, we implemented a graph-based algorithm to expand our crisscross handle library from 32 to 64 sequence pairs. The expanded library further reduces parasitic interactions, and allowed us to investigate the impact of handle library orthogonality (Type-1 interactions) on crisscross assembly.

With this foundation, investigating new applications for crisscross origami becomes substantially more accessible. A key strength of crisscross DNA origami is its nanoscale positioning accuracy across micronscale spans. This capability enables applications such as the 3D positioning of optical elements [20] or artificial antigen-presenting cells that require precise nanoscale organization of antigens across a micronscale surface [21]. The ability to produce large numbers of identical micron-scale particles with well defined geometry enables their use as microrobotic devices, including microswimmers [31] with cargo delivery capabilities [32].

Our findings also lay the groundwork for further advances of the core crisscross platform. While parasitic interactions are minimized at the design stage, complementary strategies could further reduce their impact during assembly. For example, solid-phase approaches that partition the assembly into smaller steps limit the number of slats present in solution simultaneously and thus reduce the likelihood of parasitic interactions. In addition, transient heat spiking during assembly could disrupt aggregates and limit the accumulation of parasitic interactions. Furthermore, new design strategies could be explored. Introducing new slat types with different geometries could enable additional structural motifs or open alternative routes to three-dimensional architectures that do not rely on the layer-stacking approach as detailed in this paper.

Finally, while the handle library used in this work comprises 64 sequence pairs, smaller libraries can still yield practical assemblies when optimized using the evolutionary algorithm. As shown in Fig. 5, acceptable megastructure yields can be obtained at higher Loss values, particularly for smaller designs like the Square. In this regime, reducing the handle library size represents a viable trade-off between yield and cost. For example, using a library of 24 sequence pairs, our evolutionary algorithm achieved a Loss of 3.7 for the Square design. Reducing the library from 64 to 24 sequence pairs decreases the total number of staple strands from 4096 to 1536, significantly lowering the cost of entry. To reduce costs even further for targeted applications, purchasing only the subset of staple strands required for a single design is also sufficient.

## Extended Data Figures

**Extended Data Fig. 1.**
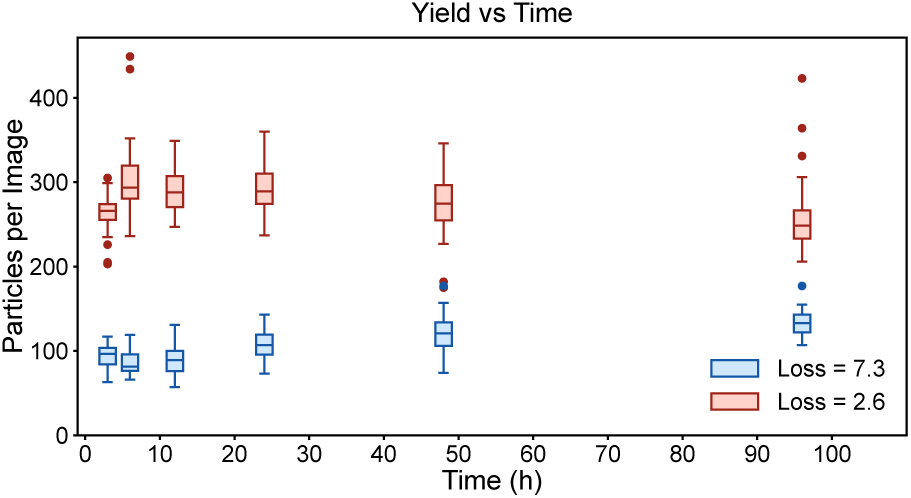
Assembly kinetics of megastructures with different Losses. Two Square megastructures with Losses of 7.3 and 2.6 were selected from the experiment shown in Fig. 5 and monitored over time. Assembly reactions were sampled at time points between 3 h and 96 h, while the data in Fig. 5 correspond to an incubation time of 41 h. For each time point and condition, we recorded 30 TEM images, counted the number of completed megastructures, and summarized the statistics in box plots. Both time traces remained constant throughout the monitored interval, with the design with higher Loss consistently yielding fewer assembled structures. This indicates that assembly in the experiment shown in Fig. 5 had already reached completion and that the data represent the final assembly yield. For these experiments, purification was initiated at each sampling time point by adding magnetic capture beads to the assembly mixture, allowing assembly to continue during the 15.5 h capture process. Consequently, while the data demonstrate that Loss affects the final yield, a potential influence on the assembly rate cannot be excluded (see Method 19).

**Extended Data Fig. 2.**
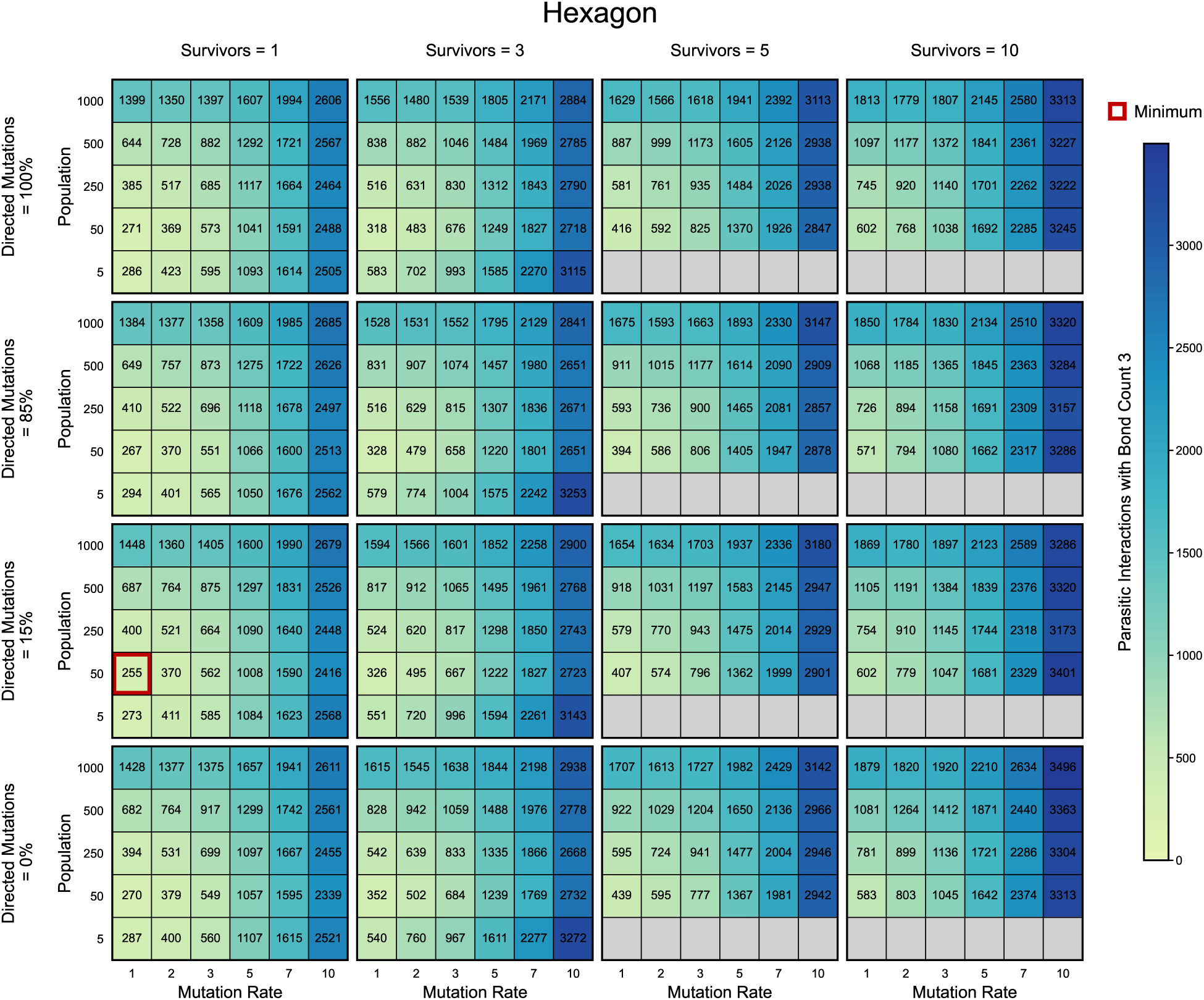
Optimal parameter search of the evolutionary algorithm on a Hexagon. This figure shows heat maps of the final number of parasitic interactions with bond count 3 obtained when evolving handle assignments for the two-layer Hexagon megastructure, plotted as a function of the user-adjustable parameters of the evolutionary algorithm (Method 4). These parameters are: *population size*, which specifies how many new candidate assignments we create per generation; *survivors*, defined as the number of top-performing candidates that seed the next generation; *mutation rate*, given as the expected number of handle swaps applied to each survivor to generate an offspring; and *directed mutation probability*, which defines how often mutations are restricted to slats involved in parasitic interactions with the highest bond count. To compare parameter combinations on an equal computational footing, we fixed the total number of megastructure evaluations by adjusting the number of generations such that the product of population size and generations remained constant at 500,000, keeping the total number of generated assignments the same in all runs. The optimal parameter combination is highlighted in red (*mutation rate* = 1, *survivors* = 1, *population size* = 50, *directed mutation probability* = 15%). Overall, small survivor numbers, small mutation rates, and population sizes around 50 yield the lowest number of parasitic interactions with bond count 3 under the fixed-budget constraint, whereas directed mutations provide only a modest improvement with a shallow optimum near a directed mutation probability of approximately 15%. We repeated the parameter search for the multilayer Sunflower design, and the results are shown in Extended Data Fig. 3. For the Sunflower, the same trends were observed: low survivor numbers, a mutation rate of one, and a directed mutation probability of 15% performed best while the optimal population size shifted to five.

**Extended Data Fig. 3.**
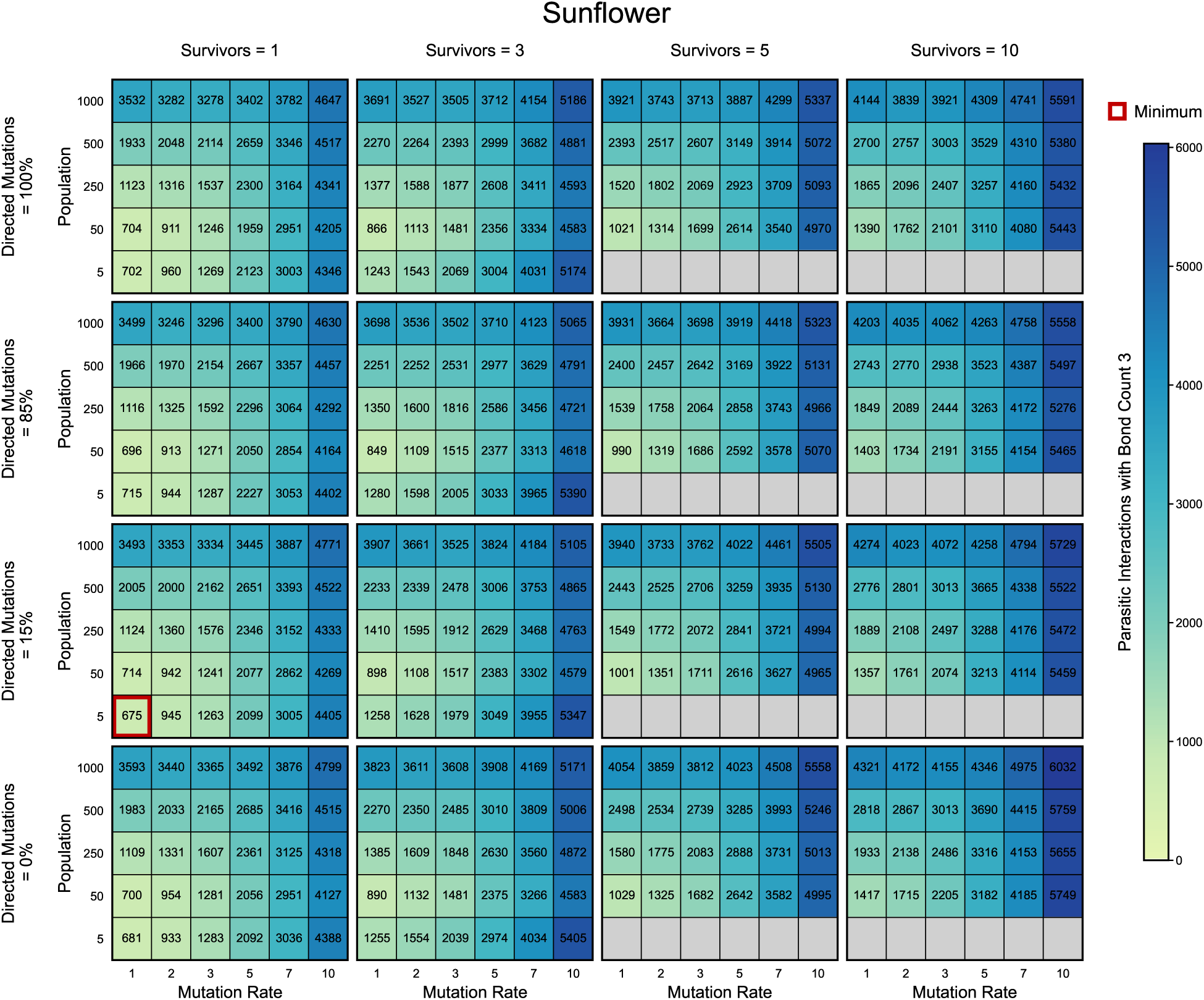
Optimal parameter search of the evolutionary algorithm on a Sunflower. This figure shows heat maps of the final number of parasitic interactions with bond count 3 obtained when evolving handle assignments for the multilayer Sunflower megastructure. The analysis follows the same procedure as in Extended Data Fig. 2. Similar to the Hexagon, low survivor numbers, a mutation rate of one, and a directed mutation probability of 15% perform best. However, the optimal parameter combination differs in the population size, which shifts to five for the Sunflower geometry.

## Methods

### Method 1: #-CAD, Scripting Library Development & General Data Analysis

#-CAD was developed in Dart using the Flutter framework. A single codebase was used for all versions of the application (web, macOS, Windows & Linux). We made use of other open-source libraries such as the three js package for 3D structure viewing. The accompanying scripting library was developed using Python 3.11, and is available for download on PyPi or GitHub. The library also makes use of various open-source libraries such as Blender’s *bpy*, which allows it to directly export Blender files from megastructure designs. Both the scripting library and #-CAD use a single unified file format stored within Microsoft Excel worksheets (.xlsx). Files can also be edited by hand for convenience.

All data analysis and figure generation was conducted in Python using standard graphical libraries such as *matplotlib* or *pandas*. Scripts for regenerating figures are provided in #-CAD’s GitHub homepage.

### Method 2: Megastructure Design using #-CAD

All megastructures in this work were designed with #-CAD. A detailed manual is provided on our GitHub homepage, but a summary of the workflow is given below:

- First, place slats on the 2D design canvas. Megastructures can have any number of layers stacked vertically, and slats can be placed in all allowed orientations on any layer. For 90° designs, slats can be placed horizontally or vertically. For 60°designs, slats can adopt three distinct orientations, each rotated by 60° relative to the others. Convenience features such as the ability to place multiple slats in one go are also available.
- When the design is complete, run the evolutionary algorithm (see Method 5) to optimize handle library assignment and thereby reduce parasitic interactions.
- Place the seed DNA origami structure above or below a set of 16 slats as the starting point for assembly. The seed must cover 80 handle attachment points, and slats cannot be placed parallel to the seed’s long axis.
- Cargo can be placed on the 2D canvas as single attachment points through #-CAD’s cargo interface at positions where assembly handles are typically located.
- Save the design to an Excel workbook or export liquid handler instructions directly. Graphics can be exported from #-CAD or generated using the Python scripting library.

Details on exporting a design to a liquid handler are provided in Method 7.

### Method 3: Computing the Parasitic Interaction Bond Count of a Megastructure

To compute the effective parasitic interaction bond count (Loss) of a megastructure handle assignment, we used the following procedure:

- Extract all slats from a given handle assignment (Method 2) and sort them into two groups: plus-handle bearing slats and minus-handle bearing slats. Slats containing both types appear in both groups.
- For each slat with assembly handles, encode the handles as a one-dimensional ordered array of integer IDs, where each ID corresponds to a unique handle sequence in the library. Unoccupied positions are encoded as zeros.
- Repeat this process for minus-handles. Complementary plus-handle and minus-handle pairs share the same integer ID.
- For any pair of plus-handle and minus-handle arrays, compare the two arrays position by position and count the number of positions where the IDs match. Zeros are ignored.
- Repeat this comparison for all possible relative alignments of the two arrays. In more detail, we begin with the leftmost entry of one array aligned with the rightmost entry of the other and then iteratively shift one array stepwise to the right. This shifting procedure is analogous to a discrete convolution, except that instead of multiplying entries, we count positions where identical nonzero integers align.
- Flip one array and repeat the above operation. With 32 handle positions on each slat, there are 126 unique pairwise comparisons between a plus-handle slat and a minus-handle slat.
- To calculate the Loss, apply the equation in Fig. 5a using a dimensionless Δ*G* of 10.
- Repeat the above calculations for every plus-handle and minus-handle array combination available.
- The Loss of a handle assignment can then be computed from the resulting list of parasitic interaction bound counts.

The above procedure was implemented in both Python and C. The Python version uses *numpy*’s array format for vectorization and can be used directly within the scripting library. The C version is more memory-efficient and is therefore suitable for larger designs and is our primary choice for the evolutionary algorithm (see Method 4). The C version is importable within our Python scripting library as a sub-package called `eqcorr2d`.

### Method 4: Evolutionary Algorithm Development

The evolutionary algorithm optimizes the handle assignments of a megastructure by minimizing the effective parasitic interaction bond count between slats (Loss).

User-defined parameters:

- *P* – population size per generation.
- *s* – number of survivors retained each generation.
- *H* – number of unique sequences in the handle library.
- *G* – total number of generations to run.
- *m* – expected number of mutations per offspring (mutation rate).
- *p_d_* – probability of using directed rather than random mutation targeting.

Workflow:

- Initialize a population of *P* candidates either randomly or by including a user-defined seed array. Each candidate consists of a complete plus:minus handle assignment for the megastructure, drawn from a library of *H* unique handle types.
- Compute the Loss for each candidate. Retain the best *s* candidates (survivors); discard all remaining candidates.
- The next generation is created from mutated copies of these survivors. Generate a new offspring by selecting a survivor at random and mutating each position of its plus:minus handle array with probability *p_m_*. This probability is normalized such that the expected total number of mutations per offspring is *m*. For a handle array that contains *N* mutable positions we use *p_m_* = *m/N*, which yields an approximately Poisson-distributed number of mutations with mean *m*.
- To concentrate search pressure on the most problematic regions of a survivor, we implemented directed mutations: with probability *p_d_*, mutation targets are restricted to slats that were involved in parasitic interactions with the highest bond count; with probability 1 *− p_d_*, mutations are applied uniformly at random. Importantly, the expected number of mutations *m* is held constant regardless of whether directed or random targeting is used. Because the number of mutations is Poisson distributed we may also produce zero mutations in a single pass. We therefore enforce at least one mutation in such cases to ensure continued progress. Consequently, the effective mutation rate becomes *m*_eff_ = *m* + *e^−m^* since Pr(*K* = 0) = *e^−m^* for a Poisson-distributed number of mutation events with mean *m*.
- Additionally, we retain an untouched copy of the best survivor in each generation to guarantee retention of the lowest-Loss design observed so far.
- Repeat the above for *G* generations.

### Method 5: Applying the Evolutionary Algorithm

For each design presented in Fig. 4, we used the evolutionary algorithm to optimize the handle assignments. We ran the algorithm until it appeared to plateau, typically for more than 1000 generations, which usually resulted in a 2–3 unit improvement in the Loss compared to the initial random assignment. The parameters followed the best practices shown in Extended Data Fig. 2, although minor variations were used between designs to reduce runtimes.

The algorithm can either be run directly within the scripting framework or through #-CAD. #-CAD comes bundled with a python server containing the C based evolutionary algorithm and allows use of the algorithm directly within the graphical interface. The algorithm can be run on a personal computer, but multiple CPU cores increase the computation speed. A typical run takes approximately 4 h for small designs of about 192 slats and up to 12 h for larger designs of about 368 slats when using eight parallel threads, such as for the bird in Fig. 4a.

For Extended Data Fig. 2 and Extended Data Fig. 3, the parameter sweep was conducted on Harvard Medical School’s O2 cluster running Red Hat Enterprise Linux version 9.7.

### Method 6: DNA Material Preparation

All DNA used for this work was purchased from IDT. Slat staple strands and crisscross handles were purchased at 100 nmol scale with standard desalting. Cargo staple strands were purchased at either 10 nmol or 100 nmol scale, also with standard desalting. DNA origami scaffold derived from the M13 bacteriophage P8064 ssDNA plasmid was produced through a fermentation process and purchased from uBriGene. The same scaffold was used for both the slat and seed origami structures (see Method 10). All strands with fluorophores were bought at the 100 nmol scale with HPLC purification from IDT (except for those described in Method 15).

To facilitate automation, all assembly handle, cargo and seed strands were purchased in Echo-compatible 384-well plates. The assembly handles were purchased dry (full yield) then resuspended to a final concentration of 1 mM in ultra-pure water using an Eppendorf EpMotion 5075. For certain plates, we instead resuspended the DNA using manual multi-channel pipettes, selecting a volume that would ensure the DNA concentration is 1 mM or more for all wells.

From these ‘master’ plates, we created various ‘working stock’ 384-well Echo-compatible plates at a concentration of either 100, 200, or 500 µM in 1×TEF (5 mM Tris base, 1 mM EDTA, pH 8.0), using a ThermoFisher Multidrop Combi or manual multichannel pipettes to expedite buffer transfer. Master stock plates were stored at -20°C while working stock plates were stored at 4°C. Working stock plates were replenished from the master stocks as necessary. A similar process was followed for cargo and seed strands, with minor variations to the final concentrations based on the production yield provided by IDT. The new handle library was used for all megastructure designs unless otherwise specified.

### Method 7: Slat Handle Pooling

Slats consist of two groups of DNA: core staple strands and variable handles. Each six-helix bundle slat contains 64 variable handles; 32 on the H2 helix and 32 on the H5 helix. The core staple strands were pooled by hand into a single mixture which can be used for all slats. The variable handles were pooled as required to form customized slats for each unique design. The workflow used to pool variable handles was as follows:

- Starting from a #-CAD design file, import excel sheets containing the sequences and positions of variable handles in a set of input plates. A standardized format for a new plate is available on our GitHub page. Plates can be imported both into the scripting library and into #-CAD.
- Assign handles to a design. Handles can automatically be matched to slat positions through their ID, which is either an assembly handle, a cargo handle or a seed-binding handle.
- For slat positions without custom handles, assign replacement ‘flat’ handles which effectively render that slat position non-binding.
- Once this is complete, utilize #-CAD or the scripting library to export .csv instructions for a liquid handler to transfer staple strands from input ‘source’ plates to ‘destination’ 96-well plates containing the variable handle pools. A single well is created for each slat in the design.
- Conduct the pooling operation using a liquid handler of choice. For all our designs we used a Beckman Coulter Echo 525 combined with an Access Laboratory Workstation to automate the transfers for source and destination plates in one go.
- The above is all automated within our scripting library, which can also provide guideline files and graphics to help organize and manage slat pool layouts in destination plates.

Core staple and assembly handle sequences have been provided in the Supplementary Information, along with a caDNAno file for the six-helix bundle slat.

### Method 8: Slat Folding

Slats were typically folded directly in 96-well plates containing the assembly handle staple strands distributed by our liquid handler. The mixture of structural staple strands, scaffold, buffer, and magnesium was added to each slat preparation to achieve a final concentration of 50 nM for the scaffold, at least 500 nM for the staple strands, 1×TEF, and 6 mM MgCl_2_. All slats were individually folded in one batch using the standard thermal annealing ramp from [19] (80 °C for 10 min; 60–45 °C in 160 steps of 6.75 min, decreasing by 0.1 °C per step; 16 °C thereafter until sample collection).

### Method 9: Slat Purification

After annealing, slat mixtures were combined into pools. We adhered to the general guideline of limiting the number of slats in a pool to at least 16 and fewer than 32. Each pool was then PEG purified individually using the following protocol:

- Add 1 M MgCl_2_ to the slat pool to raise the Mg concentration to 15 mM.
- Add 3× PEG solution (1×TEF, 22.5% PEG-8000, and 765 mM NaCl) to the pool to reach a final concentration of 1×. Mix well.
- Spin the pool at 16 kg for 30 min at room temperature.
- Carefully remove the supernatant without dislodging the pellet.
- Add 150 µl of 1×TEF with 20 mM MgCl_2_ (without dislodging the pellet) and spin again at 16 kg for 30 min.
- Remove the supernatant and add 1×TEF with 10 mM MgCl_2_ to yield a total slat concentration of 2 µM or lower.
- Allow the slats to disperse in solution by shaking the pool for 1 h or longer at 33 °C and 800 rpm.
- Use a Nanodrop to confirm the final concentration. Minimize the delay between removal from the thermoshaker and the Nanodrop measurement to reduce measurement error. Yields for slat pools typically range from 60–100%.
- Instructions and volumes for the above steps can be pre-computed using our scripting library.

### Method 10: Seed Folding & Purification

To reduce the number of unique parts needed for crisscross production, we re-designed our seed origami to use the p8064 scaffold rather than the p8634 scaffold used previously [19]. We also added eight poly-A extensions on selected seed origami staples to enable magnetic bead pulldown (Method 12). The staple set and the caDNAno schematic for this design are provided in the Supplementary Information. To fold and purify the seed origami, we used the following workflow:

- Combine all seed staple strands with the p8064 scaffold, magnesium, and buffer to achieve final concentrations of 40 nM for the scaffold, 400 nM for the staple strands, 1×TEF, and 10 mM MgCl_2_.
- Anneal the seed using the following thermal ramp: 80 °C for 15 min; 60–55 °C for 2 h 15 min (-0.1 °C per 2:42, 50 cycles); 55–25 °C for 13 h 45 min (-0.1 °C per 2:45, 300 cycles); then hold at 25 °C before use.
- Purify the seed by agarose gel electrophoresis. Prepare a 1% gel by mixing 175 ml of 0.5×TBE (45 mM Tris, 45 mM boric acid, 0.78 mM EDTA) with 11 mM MgCl_2_, and dissolve the agarose by heating to boiling in a microwave. Let the solution cool for 5 min, then add 17.5 µL of 10,000× SYBR Safe to obtain a 1× pre-stained gel for casting in a Thermo Scientific Owl EasyCast B2 setup. In our setup, we tape the comb to create a single long well that accommodates approximately 300 µl of sample.
- Mix the seed with 6× NEB (no SDS) loading dye to a final concentration of 1×. Load each well to its maximum capacity. In our setup, the single long well is filled with approximately 300 µl.
- Run the gel for 105 min at 85 V at room temperature.
- Identify the seed monomer band using a blue-light transilluminator and excise the corresponding gel fragment.
- Cut the gel into small pieces, transfer them to a Freeze’N Squeeze (Bio-Rad) column, and extract the seed following the manufacturer’s protocol.
- Pre-wet a 50 kDa Amicon (Merck) filter with 1×TEF + 10 mM MgCl_2_ and spin for 6 min at 5 kg.
- Remove the buffer from the Amicon collection tube and add the extracted seed solution, ensuring the total volume does not exceed 500 µl. Spin again for 6 min at 5 kg.
- Add additional seed solution up to a maximum of 500 µl, spinning after each addition.
- For the final spin, top up to 500 µl using 1×TEF + 10 mM MgCl_2_.
- Recover the concentrated sample by flipping the filter upside down and spinning for 3 min at 3 kg.
- Confirm the final concentration using a Nanodrop.
- Adjust the Amicon spin time or speed as needed to achieve the desired concentration.
- Expect a final yield of 10–20% relative to the starting scaffold concentration.
- An alternative method to estimate the sample’s concentration is to run an agarose gel with a sample of the concentrated origami and a sample of the non-concentrated purified origami prior to Amicon concentration. Band fluorescence intensity comparisons via image analysis-based quantification can then be used to determine the true monomer concentration.

### Method 11: Megastructure Assembly

With purified slat pools and seed origami available, megastructure assembly can be initiated by combining the required slats with the seed in assembly buffer. In brief:

- Predefine a final seed concentration for the assembly mixture. We used 0.5–1 nM.
- Disperse slat pools by shaking them at 800 rpm and 33 °C for at least 30 min.
- Add slat pools to achieve at least a 5× concentration ratio of any individual slat to seed. The total slat concentration should not exceed 2 µM because higher concentrations can slow the assembly kinetics. For smaller designs, a 10× slat-to-seed ratio is feasible and can further accelerate assembly.
- Add Tween-20, MgCl_2_, and 10×TEF to reach final concentrations of 0.01% Tween, 15 mM MgCl_2_, and 1×TEF (assembly buffer).
- Heat the mixture to 45 °C for 4 h to inhibit slat:slat binding and promote seed binding.
- Incubate at 37 °C for the desired duration. For squares we incubated for 24–48 h, whereas larger designs typically required 3–10 days. Longer incubation times yield more fully assembled structures.

### Method 12: Megastructure Magnetic Bead Pulldown

After assembly, a megastructure can be imaged directly (Method 14), but the mixture contains a large excess of unincorporated slats. To extract megastructures from this solution, we perform magnetic bead purification using 100- or 150-nm streptavidin-coated magnetic beads from Ocean Nanotech (SV0100/SV0150):

- Extract 100 µl of beads from the storage solution and wash twice with 200 µl of DNA binding buffer (20 mM Tris pH 7.5, 1 M NaCl, 1 mM EDTA, and 0.05% Triton X-100). We use magnetic racks from Sergi Lab Supplies to perform washes and magnetic pulldowns.
- Resuspend the beads in 100 µl of DNA binding buffer and add 1.26 µl of 1 mM biotinylated capture DNA (Supplementary Information), which provides a large excess of DNA relative to the number of beads.
- Incubate the mixture for 1 h at room temperature on a rotator spinning at a minimum of 16 rpm.
- Wash the beads three times with 200 µl of assembly buffer and resuspend them in 100 µl of assembly buffer.
- Add assembled megastructures to achieve a final bead-to-structure ratio of 3–5.
- Allow the megastructures to attach to the biotinylated capture DNA through their seed, which carries poly-A extensions. Incubate for at least 4 h at 37 °C on a rotator spinning at 16 rpm or higher.
- Wash the beads 3–5 times using at least twice the reaction volume of assembly buffer. Increasing the number and volume of washes removes more unincorporated slats at the cost of slightly reduced yield. Perform washes at 37 °C to prevent aggregated slats from obstructing bead movement.
- Resuspend the beads in assembly buffer supplemented with 50 µM invader DNA (Supplementary Information). Incubate for at least 4 h at 37 °C on a rotator spinning at 16 rpm or higher. The invader DNA cleaves megastructures from the beads by a strand-displacement reaction, releasing them into solution.
- Pull down the beads and collect the solution containing the released megastructures.
- Reaction volumes can be scaled as needed when processing larger sample quantities.

### Method 13: DNA Invader Cleanup

As an optional final step to remove invader DNA, reduce free slat concentrations without bead pulldown, or perform buffer exchange, megastructures were further purified by repeated centrifugation. The workflow was as follows:

- Top up the megastructure solution to a minimum of 50 µl with assembly buffer.
- Spin the sample for 10 min at 15 kg. The protocol also works at 1–14 kg, although lower speeds can reduce megastructure yield.
- Remove the supernatant, leaving approximately 10 µl at the bottom of the tube.
- Top up the sample to 50 µl with buffer and repeat the centrifugation step.
- Repeat the above steps at least three times to remove invader DNA. The procedure can be extended to deplete slats after solution assembly.
- Dilute the final sample as desired. If the sample was also purified by bead pulldown, a Nanodrop measurement provides a rough estimate of the final yield.

### Method 14: Electron Microscopy Imaging

Megastructure integrity and gallery images were verified by TEM. We followed one of two protocols to stain and image our structures:

We negatively stained samples when we wished to see the granular detail of individual slats on a megastructure:

- Treat a formvar/carbon-coated copper grid (400 mesh, FCF400-CU, Electron Microscopy Sciences) in a plasma chamber (PELCO easiGlow) for 15s/25mA.
- Deposit 3-4 µl of a sample on the grid clamped on tweezers and allow to incubate for 4 mins.
- Wick the sample using the edge of a filter paper, then deposit 10 µl of 0.5% uranyl formate or 2% uranyl acetate.
- Wick the stain immediately using the edge of a filter paper.
- Deposit an additional 10 µl of stain and allow to incubate for 40-60 s. Wick the stain using the edge of a filter paper and allow to dry.

We positively stained samples when conducting large-scale counting or doing quick sample integrity checks:

- Treat a formvar/carbon-coated copper grid (400 mesh, FCF400-CU, Electron Microscopy Sciences) in a plasma chamber (PELCO easiGlow) for 15 s/25 mA.
- Deposit 3-4 µl of your sample on the grid and allow to incubate for 2-4 mins.
- Wick the sample using the edge of a filter paper, then deposit 4 µl of 0.5% uranyl formate or 2% uranyl acetate.
- Blot the stain immediately using full contact with the flat side of a filter paper.

All imaging was conducted at 80kV on a JEOL JEM 1400 Plus microscope. Image brightness and contrast were adjusted linearly post-imaging to optimize data display. No other corrections were applied.

### Method 15: Fluorescent Slats

To create fluorescent slats, 31 core staple strands (Supplementary Information) were purchased with 3’ amino modifications from IDT (standard desalting). These were conjugated to NHS-ester fluorophores as a pool using the below workflow:

- Resuspend all staple strands to 500 µM in ultra-pure water.
- Combine 20 µl of each staple together into a pool.
- Combine 80µl of the pool with 25 µl of 1M NaHCO_3_ (pH 7.7) and 20 µl of ultra-pure water to a final oligo pool concentration of 320 µM and NaHCO_3_ concentration of 0.2 M.
- Combine 50 µl of the oligo pool with 30 µl of 5.33 M fluorophore dye dissolved in DMSO.
- Incubate at room temperature for 2 hours with 750 rpm shaking. Keep the sample covered from light from here on.
- Ethanol precipitate the pool to remove excess dye. In brief (protocol based on [33]):
  – Mix oligo pool with 3× (w.r.t. volume) ice cold 100% ethanol and 0.1× (w.r.t. volume) 3 M sodium acetate.
  – Cool solution in a -80 °C freezer for 1 hour, then spin at 16 kg for 30 mins at 4 °C.
  – Remove supernatant and add 500 µl of ice cold 75% ethanol. Re-spin at 16 kg for 5 mins at 4 °C.
  – Remove the supernatant, and repeat the addition of 75% ethanol, spin and supernatant removal.
  – Allow the final pellet to air dry for 2 hours.
  – Resuspend in 70 µl of ultra-pure water by shaking at 37 °C/650 rpm for 30 mins.
  – Nanodrop the final solution to confirm concentration.

The workflow was conducted separately for two fluorophores: Atto 488 NHS ester (AAT Bioquest, 2815) and Atto 565 NHS ester (ATTO-TEC, 72464), creating two individual staple pools.

To assemble a fluorescent slat, the same procedure as in Method 8 was carried out, but the core staple mix was replaced with an alternative mixture containing the fluorescent staple pool.

### Method 16: Nanocube Folding & Megastructure Attachment

The component strands were purchased from IDT at 10 nmol scale (standard desalting). We rehydrated the dehydrated nanocube strands in water at approximately 100 µM and pooled all strands at equal volumes. The 28 nanocube strands, originally published with 4T brushes [19], were modified to use 6T brushes. We additionally modified one strand to include a 21 nt cargo handle on its 3’ end and another to include an Atto488 fluorophore on its 5’ end.

The nanocube was folded by combining 1 µM of each strand (with the handle-tagged strand at 2 µM) in 1×TEF and 40 mM MgCl_2_. The mixture was subjected to the following temperature gradient: an initial step at 80 °C for 10 min, followed by a ramp from 65–36 °C consisting of 290 steps of 8.69 min with a 0.1 °C decrease per step, and a final hold at 16 °C until collection. After folding, we purified the nanocube by gel electrophoresis as described in Method 10.

To attach nanocubes, we first bound megastructures containing nanocube cargo dock handles to 150 nm beads (Method 12). After washing away free slats, we added nanocubes to the megastructures in a 5× excess relative to the number of available binding sites in the assembly buffer. Hybridization proceeded overnight at room temperature on a rotator spinning at 16 rpm or higher. Subsequently, we washed away excess nanocubes and displaced the megastructures from the beads before imaging.

### Method 17: Fluorescence Imaging

To record the fluorescence microscopy images in Fig. 4, we used the following workflow:

- Prepare coverslips for imaging by wiping both sides with a Kimwipe soaked in 5% Hellmanex solution. Place the coverslips in a hot 5% Hellmanex bath and sonicate for 30 min. Rinse in distilled water and sonicate for an additional 30 min in distilled water. Store clean coverslips in distilled water. Before use, dry the coverslips with a nitrogen stream and clean them in a plasma chamber (PELCO easiGlow) for 15 s at 25 mA.
- Pre-clean a glass slide using dish soap and dry it with a nitrogen stream.
- Construct a sample flow channel by placing two precut strips of double-sided silicone tape (GENNEL; 10 mm × 25 m, clear, double-sided) adjacent to each other on a glass slide, forming a channel approximately 1.5 mm wide. Remove the tape backing and adhere a clean coverslip on top. Multiple channels can be placed on a single slide.
- Introduce a megastructure sample (approximately 1–100 pM) into the channel using a pipette. Each channel holds approximately 10 µl, and the sample should enter by capillary action.
- Seal both ends of the channel using a two-component silicone glue (e.g., Dragon Skin^TM^ 10).
- Proceed to imaging.

Images were recorded using an Olympus IX70 microscope equipped with a 100×/1.4 oil-immersion objective and an Andor Zyla-4.2P-CL 10-W camera (2048 × 2048 pixels). For each field of view, we recorded 10 frames at 100 ms exposure. In post, these frames were averaged into a single image and the brightness/contrast was adjusted linearly for optimal display.

### Method 18: Measuring the Relation Between the Loss and Yield

In Fig. 5, we tested how the assembly yield of four distinct handle assignments of a Square varies with their effective parasitic interaction bond count (Loss). We designed, assembled, and evaluated the structures as described below:

- A standard Square was designed in #-CAD.
- The handle assignments with Loss 7.3 were randomly generated. Only 32 unique handles were allowed.
- The handle assignments with Loss 4.7 were generated by running the evolutionary algorithm (Method 4) for 10 iterations using default #-CAD parameters. Only 32 unique handles were allowed.
- The handle assignments with Loss 2.6 & 2.1 were obtained by running the evolutionary algorithm using our standard criteria (Method 4) and 64 unique handles.
- The slats for all four squares were pooled and purified separately. Each megastructure was assembled with the seed concentration at 1 nM and all slat concentrations at 10 nM. All Squares were allowed to incubate at 37 °C for 41 hours.
- The standard magnetic bead pull-down procedure was used to purify all four Squares (Method 12). We incubated the megastructures with the beads for 15.5 hours at 37 °C before excess slats were washed away.
- 50 TEM images with standardized dimensions of 23.6 by 18.4 µm were recorded per handle assignment (one per grid square). All samples were positively stained (Method 14).
- Megastructure yield was determined by counting individual particles in each image using QuPath v0.6 [34].

### Method 19: Comparing Megastructure Assembly Kinetics at Different Effective Parasitic Interaction Bond Counts

In Extended Data Fig. 1, we compare the assembly kinetics of two Square megastructures with different Loss values. The experiment was conducted as follows:

- Slats for the Squares with Loss values of 7.3 and 2.6 were reused from Method 18.
- As before, megastructure assembly was initiated using 1 nM seed and 10 nM of each slat. A total volume of 70 µl was prepared for each design.
- After an initial 4 h incubation at 45 °C, samples were transferred to 37 °C for the remainder of the experiment.
- At 3, 6, 12, 24, 48, and 96 h, 10 µl aliquots were withdrawn and individually purified using the standard bead-pulldown protocol (Method 12). Each aliquot was incubated with beads for 15.5 h at 37 °C before washing away excess slats.
- All purified samples were positively stained, and 30 images (23.6 µm *×* 18.4 µm) were acquired under standardized imaging conditions.
- Megastructure yield was quantified by counting individual particles in each image using QuPath v0.6 [34].

### Method 20: Effect of Loss on Structural Completeness and Yield of Multilayer Structures

In Fig. 6, we used the Sunflower to compare how the Loss affects the structural completeness of multilayer assemblies. We used the same design to evaluate the new and old handle libraries, as shown in Fig. 7e. We conducted the experiment as follows:

- We designed the Sunflower in #-CAD.
- We created several unique handle assignments with different Losses by enforcing the restrictions listed below:
  a. Use all 64 unique handles with no placement restrictions. Loss of 2.76.
  b. Use all 64 unique handles. For each handle interface layer, only half (32) of the available handles were allowed. The allowed set alternated for each consecutive layer i.e. handles between layers 1 and 2 used IDs 1–32, whereas handles between layers 2 and 3 used IDs 33–64. Loss of 3.6.
  c. Use only 32 unique handles with no placement restrictions. Loss of 3.77.
  d. Use all 64 unique handles. For each layer interface, only one quarter (16) of the available handles were allowed. The allowed set alternated for each consecutive layer i.e. handles between layers 1 and 2 used IDs 1–16, between layers 2 and 3 used IDs 17–32, and between layers 3 and 4 used IDs 33–48. Loss of 4.69.
  e. Use only 32 unique handles. For each layer interface, only half (16) of the available handles were allowed. The allowed set alternated for each consecutive layer; for example, handles between layers 1 and 2 used IDs 1–16 and those between layers 2 and 3 used IDs 17–32. Loss of 5.11.
  f. Use only 32 unique handles from the old handle library [19] with no placement restrictions. Loss of 3.77.
- For each of the above, the evolutionary algorithm was run to optimize handle assignments with our standard criteria (Method 4). The same parameters were used for all designs: *G* = 2000, *s* = 3, *P* = 50, *m* = 2, *p_d_* = 0.85.
- All slats were pooled and purified using the standard methodology.
- Megastructures were assembled with a seed concentration of 0.7 nM and slat concentrations of 7 nM. They were allowed to incubate for 90 hours at 37 °C after the initial 4 hour 45 °C incubation.
- All megastructures were pulled down and purified using the standard bead protocol (with a capture time of 14 hours at 37 °C).
- All samples were positively stained twice on two different grids. 15 images with standardized dimensions of 23.6 by 18.4 µm were recorded for each grid.
- The number of complete ‘petals’ (stages) for each structure were counted manually using QuPath v0.6. Given the difficulty of distinguishing structures at stage 5 or 6, we counted these structures in one group. Examples of structures at all the different stages are shown in Figure 6.
- The final results were tallied by combining all images from both grids, resulting in a total of 30 images per sample.

### Method 21: Implementation of the Orthogonal Sequence Search Algorithm

We selected our 64-sequence handle library using the algorithm described in Figure 7, with the technical details of our approach provided below.

Thermodynamic hybridization free energies were computed using the NUPACK 4.0 [27] Python module at 37 °C with ionic conditions of 0.05 M Na^+^ and 0.025 M Mg^2+^.

To prepare for the sequence search, we generated the complete set of all possible 7-nt sequences, appended a 5*^′^*-terminal ‘TT’ flank, and removed any sequences containing four identical nucleotides in a row. For each remaining sequence pair, we evaluated the ‘on-target’ hybridization free energy Δ*G* and selected only those pairs within the window *−*10.25 to *−*9.2 kcal mol*^−^*^1^, yielding a final set of *n* = 1315 on-target pairs. Each sequence within a pair was assigned an arbitrary label (plus-handle or minus-handle) for bookkeeping.

For the selected subset, we computed all off-target interaction energies. This includes all plus:plus handle interactions given by 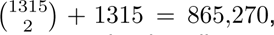 the same number of minus:minus handle interactions, and all cross-interactions between plus-handles and minus-handles given by 1315^2^ *−* 1315 = 1,727,910. In total, this required the evaluation of 865,270 + 865,270 + 1,727,910 = 3,458,450 off-target hybridization energies, yielding a complete interaction matrix that is saved and used for the subsequent optimization steps. Because the candidate sequence space scales by 4*^L^* (where *L* is the sequence length), exhaustive off-target energy evaluation grows exponentially with length and becomes impractical even for 8-mers.

Next, the precomputed interaction matrices were loaded, and an off-target energy threshold of *−*7.4 kcal mol*^−^*^1^ was applied. Any combination of sequence pairs that formed at least one undesired interaction (plus:plus, plus:minus, or minus:minus) below this threshold was classified as incompatible. This yields a conflict graph in which each vertex corresponds to a sequence pair and each edge marks an incompatibility. The goal is thus to identify a large independent set of this graph, i.e., a collection of sequences that share no conflicts [28]. Finding maximum independent sets is NP-hard and cannot be solved exactly at the required scale, so we employed a heuristic approach.

The core of the algorithm is an iterative elimination rule that removes vertices in descending order of their conflict degree, following maximum-degree pruning strategies [29, 30]. At each step, the vertex involved in the largest number of remaining conflicts is eliminated. If several vertices share the same maximum degree, a tie-breaking rule is applied: we restrict attention to the subgraph induced by the tied vertices, count how many neighbors each tied vertex has within this subgraph, and randomly remove one of those with the fewest internal connections. Vertices with many neighbors inside the tied group tend to be resolved naturally in later pruning steps.

Because a single greedy pass can settle into a particular local solution, we maintain a small population of equally good independent sets, initialized with a single greedy solution. The algorithm then performs a sequence of refinement iterations. In each iteration, every independent set in the population is perturbed by temporarily reintroducing a fixed number of previously excluded sequences. This perturbation re-creates a small, localized conflict set. To restore validity, we reapply the same degree-based elimination rule, but only to the newly created conflicts, producing a repaired independent set derived from the perturbed one.

If a repaired independent set is larger than the best set found so far, it becomes the new best solution and the population is reset to contain only this improved set. If a repaired set has the same size as the current best but represents a distinct combination of sequences, it is added to the population to preserve multiple high-quality alternatives. After each iteration, the population is pruned to a fixed maximum size by random sampling to prevent uncontrolled growth while preserving diversity.

This iterative perturb-and-repair strategy is repeated for a predefined number of iterations. The largest independent set encountered during the entire process is returned as the final library of orthogonal sequences.

Notably, the number of sequences identified by the algorithm cannot be predicted in advance for given energy thresholds. Consequently, constructing a 64-pair library required iterative adjustment of the thresholds and empirical trial-and-error.

The above algorithm can also be applied for other scenarios requiring orthogonal sequences, and is available as a standalone module, *orthoseq generator*, within our Python scripting library (Method 1).

### Method 22: Handle library verification by measuring crisscross polymerization speed

For Fig. 7d, we created a zig-zag repeating unit design which could grow indefinitely on both ends. An individual structure’s length can be measured by counting the number of corners in the ribbon. We conducted the experiment as below:

- We designed the zig-zag crisscross polymer by optimizing a standard square megastructure with a reduced set of 32 unique handle-pairs allowed. We then rearranged a subset of these handles to create a repeating unit which resulted in a final Loss of 4.73. We created two versions of this structure in the lab: one mapping the IDs to handle sequences from the old handle library and one to handle sequences the new handle library.
- All slats were pooled and purified using the standard methodology.
- Megastructures were assembled with a seed concentration of 0.5 nM and slat concentrations of 20 nM. They were allowed to incubate for 20 hours at 37 °C after the initial 4 hour 45 °C incubation. Three different magnesium concentrations were used for each design: 12.5 mM, 15 mM and 17.5 mM, along with 1×TEF and 0.01% Tween-20.
- At specific time points, 1 µl of the megastructure mixture was extracted, diluted by 10 in the corresponding magnesium buffer and shaken for 10 minutes at 37 °C/800 rpm before positive staining and TEM imaging (Method 14).
- Timepoints at 2, 8 and 12 hours were taken, resulting in a total of 18 samples.
- For each sample, we identified up to 63 individual ribbons by systematically scanning the grid from the center outward to reduce selection bias. We excluded ribbons that were crumpled or aggregated. We recorded an image of each ribbon and measured its length by counting the corners of the structure. If the counted length of a ribbon was inconsistent between measurements, we excluded that structure.
- We used the final dataset to generate Fig. 7D.

## Acknowledgments

We thank Anastasia Ershova, Christopher Wintersinger, Hawa Dembele, Sera Mathew, and Jaylan Shaw for their advice and wet lab support when we were designing our handle library and megastructure experiments. Furthermore, we thank Thomas Ferrante for his help in setting up our fluorescent microscope, and John Alberta & Talya Levitz for their support in the lab and for electron microscopy imaging. Finally, we thank ‘knightro63’ for their crucial assistance in configuring the three js library for #-CAD’s graphical release.

The O2 High Performance Compute Cluster, supported by the Research Computing Group at Harvard Medical School, was used to accelerate development of the evolutionary algorithm and the final large-scale parameter sweeps.

This project was supported by multiple funding sources: the UK Medical Research Council Precision Medicine Transition Fellowship [grant number MR/N013166/1] (M.A.), the Dana-Farber Cancer Institute Claudia Adams Barr Program for Cancer Research (M.A., W.M.S.), a Wyss Institute Northpond Alliance Director’s Fund Award (M.A.), the German Research Foundation (Deutsche Forschungsgemeinschaft, DFG) through the Walter Benjamin Programme [project number 553862611] (F.K.), the U.S. Department of Energy, Office of Science, Basic Energy Sciences, Biomolecular Materials Program [Award No. DE-SC0024136] (F.K., W.M.S.), the Carlsberg Foundation [grant CF23-1125] (M.A.D.N.), a Sloan foundation grant [grant ID G-2021-16495] (S.S.W., W.M.S.), the Harvard College Research Program (C.B.), the Korea-US Collaborative Research Fund (KUCRF) funded by the Ministry of Science and ICT and Ministry of Health & Welfare, Republic of Korea [grant RS-2024-00468463] (M.A., F.K., S.S.W., J.L., S.L., W.M.S.), Novo Nordisk Foundation [grant NNF23OC0084494] (Y.Z., W.M.S.), and the Wyss Institute Molecular Robotics Initiative (W.M.S.).

## Declarations

William M. Shih is an inventor on a patent (PCT/US2017/045013) entitled ‘Crisscross Cooperative Self-assembly’, which is related to the basic principle of crisscross assembly.

## Availability of data and materials

The entire #-CAD codebase and scripting library has been open-sourced under the MIT license, and is available on GitHub at: https://github.com/mattaq31/Hash-CAD. The code used to generate all quantitative figures in this paper has also been provided in the same repository. The electron microscopy datasets used for all analyses, along with all #-CAD design files, have been deposited to Zenodo, for which access is available at: https://doi.org/10.5281/zenodo.17914052. All DNA sequences and all caDNAno/Scadnano design schematics used in this study have been provided as Supplementary Data files.

## Authors’ Contributions

M.A., F.K., M.A.D.N., S.S.W. & W.M.S. conceptualized the idea of an end-to-end crisscross pipeline. F.K. & M.A. developed the evolutionary algorithm to minimize the Loss. M.A. built #-CAD, which used an early javascript prototype developed by C.B. as a foundation. M.A., F.K. & S.S.W. built the python scripting library. F.K. developed the orthogonal sequence generation algorithm. S.S.W., M.A., F.K.& H.C. developed the megastructure pulldown protocol. S.S.W., J.L., M.A., F.K., M.A.D.N., Y.Z. & S.H.S. optimized the overall megastructure assembly protocols. F.K., M.A.D.N., M.A., S.S.W., C.B., Y.Z., S.H.S. & J.F. designed and tested the various megastructures in the showcase gallery. F.K. & S.H.S. ran the experiments using square megastructures to assess the impact of parasitic interactions on yield and rate of assembly. M.A.D.N. designed and conducted the sunflower growth experiments with contributions from M.A., F.K & S.H.S. The extended handle library polymer experiments were conducted by S.S.W, M.A., F.K., & Y.Z. The manuscript was written by M.A. & F.K. Final data processing and analysis was conducted by F.K. and M.A. The figures were generated by M.A., F.K. & M.A.D.N. All authors reviewed, adjusted and approved the final manuscript. M.A., F.K., S.S.W., J.L., S.L. & W.M.S. secured funding for the project. M.A., F.K. & W.M.S. coordinated the overall project.

