## Supplementary Information for "A computational framework for designing micron-scale crisscross DNA megastructures"

### 1 Supplementary Note 1: Interpretation and Derivation of the Loss Function

#### 1.1 Interpretation

Here we provide a thermodynamic interpretation of the loss function introduced in Figure 5 in the main text.

$$\mathrm{Loss} = \frac{1}{-\Delta G} \ln \left( \frac{1}{N_{\text{Pairs}}} \sum_{\text{Pairs}} \sum_{\text{Interactions}} e^{-n_{ijk}\Delta G} \right). \tag{S1}$$

Here, $\Delta G$ denotes the change in Gibbs free energy for a single plus-handle binding to a minus-handle, and $n_{ijk}$ parameterizes all bond counts of the parasitic interactions between slats. Notably, we changed the index $i$ of the bound count from the main text to its multi-index form $ijk$. The outer sum runs over all possible pairs $ij$ of plus-handle bearing slats and minus-handle bearing slats, while the inner sum runs over all possible interactions $k$ between a given pair $ij$. $N_{\text{Pairs}}$ denotes the total number of possible pairs.

As mentioned in the main text, the primary purpose of the Loss function is to classify handle assignments of a megastructure based on its slats' parasitic interactions. Contributions from parasitic interactions with higher bond counts are exponentially penalized and are therefore eliminated by the evolutionary algorithm at earlier stages.

We show that, within the framework of a toy model, the Loss function can be interpreted as an effective bond count of the parasitic interactions. In this framework, we approximate all possible interactions between slats by a single effective reaction between all plus-handle bearing slats $X$ and all minus-handle bearing slats $Y$, forming a single representative complex $Z$:

$$X + Y \rightleftharpoons Z.$$

Within this approximation, complex formation is characterized by a single effective change in Gibbs free energy $\Delta G_{\text{eff}}$. From $\Delta G_{\text{eff}}$, we compute an effective bond count of the parasitic interactions by normalizing with the change in Gibbs free energy $\Delta G$ for a single plus-handle binding to a minus-handle. The effective parasitic interaction bond count is thus given by

$$n_{\text{eff}} = \mathrm{Loss} = \frac{\Delta G_{\text{eff}}}{\Delta G}.$$

Specifically, a Loss of $n_{\text{eff}} = 2$ means that plus-handle bearing and minus-handle bearing slats interact as if they could form a single complex with the binding energy corresponding to two handles.

#### 1.2 Toy Model Formulation

We first formulate the model for a simple square-shaped crisscross megastructure, as shown in Figure 1 of the main text. In our designs, plus-handle sequences are restricted to the top helix (h5) of each slat, whereas minus-handle sequences are restricted to the bottom helix (h2). Each plus-handle is designed to be fully complementary to one corresponding minus-handle. Our square design therefore consists of a top layer with 32 slats, each carrying 32 minus-handles on the bottom helix, and a bottom layer with another 32 slats, each carrying 32 plus-handles on the top helix (h5). Thus, for the square structure, we have two groups of slats: one bearing plus-handles and one bearing minus-handles. In this toy model we consider only perfectly complementary plus-handle:minus-handle binding interactions (Type 2 in Figure 1B) and neglect all partially complementary interactions (Type 1 in Figure 1B). Consequently, only plus-handle bearing slats can hybridize with minus-handle–bearing slats, whereas plus-handle:plus-handle and minus-handle:minus-handle interactions are excluded.

We can therefore divide the slats into two groups: $X_i$ with $i = 1 \ldots 32$ are the plus-handle bearing slats, and $Y_j$ with $j = 1 \ldots 32$ are the minus-handle bearing slats. A specific slat $X_i$ and $Y_j$ can form different binding configurations, which we denote as $Z_{ijk}$, where the index $k$ labels the different possible alignments of these two slats. These configurations are identified by our algorithm by sliding slat $Y_j$ along slat $X_i$ and detecting positions where complementary plus-handles and minus-handles align (see Figure 5A).

For each configuration $Z_{ijk}$, we assign $n_{ijk}$, which corresponds to the number of plus-handle:minus-handle bonds formed between the two slats in that specific configuration. We define the change in Gibbs free energy of binding for the configuration $Z_{ijk}$ as

$$\Delta G_{ijk} = \Delta G^0 + n_{ijk}\, \Delta G, \tag{S2}$$

where $\Delta G^0$ is an initiation penalty for complex formation and $\Delta G$ is the Gibbs free energy change associated with a single plus-handle binding to its complementary minus-handle.

In our model we work with dimensionless energies and therefore normalize all free energy values by $RT \approx 0.62$ kcal/mol, using $T = 273 + 37 = 310$ K and $R = 1.987 \times 10^{-3}$ kcal mol$^{-1}$K$^{-1}$. We assume for simplicity that all handle sequences have the same binding free energy $\Delta G$. The actual Gibbs free energies of hybridization computed by NUPACK 4.0 (`celsius=37`, `[Na`$^+$`]=0.05 M`, `[Mg`$^{2+}$`]=0.025 M`) lie in the range $[-10$ kcal/mol, $-8.45$ kcal/mol$]$, which corresponds to $[-16.1, -13.6]$ in dimensionless units. However, because the handles are tethered to the slats, their effective binding strength is likely reduced compared to the ideal solution case. To keep the model simple while staying in the right physical range, we use a single heuristic value of $\Delta G = -10$, which captures the correct order of magnitude.

With these definitions at hand, we define the equilibrium constant for dimer formation in a specific binding configuration $k$ of a slat pair $(X_i, Y_j)$ as

$$K_{ijk} = \frac{[Z_{ijk}]}{[X_i][Y_j]} = e^{-\Delta G_{ijk}} = e^{-(\Delta G^0 + n_{ijk}\Delta G)}. \tag{S3}$$

Here, $[Z_{ijk}]$ denotes the concentration of the dimeric complex formed in configuration $k$, and $[X_i]$ and $[Y_j]$ denote the concentrations of the free, unbound slats. The total concentrations of each slat species are conserved. Thus, the total concentrations $[X_i^0]$ and $[Y_j^0]$ of every plus-handle bearing slat $i$ and minus-handle bearing slat $j$ are given by

$$[X_i^0] = [X_i] + \sum_j \sum_k [Z_{ijk}], \tag{S4}$$

$$[Y_j^0] = [Y_j] + \sum_i \sum_k [Z_{ijk}]. \tag{S5}$$

In practice, slats can form multimeric parasitic complexes in which multiple slats bind together in a disordered manner, potentially leading to macroscopic aggregates, as shown in Figure 1 of the main text. Importantly, the formation of any higher-order aggregate is initiated by dimer formation. Thus, dimerization can be regarded as a sentinel event marking the onset of parasitic multimerization.

#### 1.3 Derivation

First we compute the total concentration of dimeric complexes $Z$ and insert Equation S3:

$$[Z] = \sum_{i=1}^{n_h} \sum_{j=1}^{n_{ah}} \sum_k [Z_{ijk}] = \sum_{i=1}^{n_h} \sum_{j=1}^{n_{ah}} \sum_k K_{ijk}\, [X_i][Y_j] = \sum_{i=1}^{n_h} \sum_{j=1}^{n_{ah}} [X_i][Y_j] \sum_k e^{-\Delta G^0 - n_{ijk}\Delta G}. \tag{S6}$$

Here $n_h$ is the total number of plus-handle bearing slats and $n_{ah}$ is the total number of minus-handle bearing slats.

The total concentrations of plus-handle-bearing ($[X^0]$) and minus-handle–bearing slats ($[Y^0]$) are obtained by summing over all species of the respective type, whether free or bound:

$$[X^0] = \sum_{i=1}^{n_h} [X_i^0] = \sum_{i=1}^{n_h} \left( [X_i] + \sum_{j=1}^{n_{ah}} \sum_k [Z_{ijk}] \right) = [X] + [Z], \tag{S7}$$

$$[Y^0] = \sum_{j=1}^{n_{ah}} [Y_j^0] = \sum_{j=1}^{n_{ah}} \left( [Y_j] + \sum_{i=1}^{n_h} \sum_k [Z_{ijk}] \right) = [Y] + [Z] \tag{S8}$$

Here we used the definition of $[Z]$ and defined the total free slat concentrations $[X] = \sum_{i=1}^{n_h} [X_i]$ and $[Y] = \sum_{j=1}^{n_{ah}} [Y_j]$.

To find an effective description of the model we now apply a mean-field approximation. We assume that all plus-handle bearing slats $X_i$ interact on average in the same way with all minus-handle–bearing slats $Y_j$.

This implies that the concentrations of free slats within each group are equal. We therefore write

$$[X_i] = \frac{[X]}{n_h} \quad \text{and} \quad [Y_j] = \frac{[Y]}{n_{ah}},$$

where $[X]$ and $[Y]$ denote the total concentrations of plus-handle bearing and minus-handle bearing slats, respectively. Here we replace the individual slat concentrations by their average across each group. Note that we implicitly assume that the initial slat concentrations $[X_i^0]$ and $[Y_j^0]$ are identical within each group, which is typically the case in our experiments. Inserting this into Equation S6 yields

$$[Z] = \sum_{i=1}^{n_h} \sum_{j=1}^{n_{ah}} \frac{[X]}{n_h} \frac{[Y]}{n_{ah}} \sum_k e^{-\Delta G^0 - n_{ijk}\Delta G} = \frac{[X][Y]}{n_h n_{ah}} e^{-\Delta G^0} \sum_{i=1}^{n_h} \sum_{j=1}^{n_{ah}} \sum_k e^{-n_{ijk}\Delta G}. \tag{S9}$$

Dividing this equation by $[X][Y]$ yields a definition of the effective equilibrium constant $K_{\text{eff}}$:

$$K_{\text{eff}} = \frac{[Z]}{[X][Y]} = \frac{1}{n_h n_{ah}} e^{-\Delta G^0} \sum_{i=1}^{n_h} \sum_{j=1}^{n_{ah}} \sum_k e^{-n_{ijk}\Delta G}. \tag{S10}$$

We can now define an effective change in Gibbs free energy $\Delta G_{\text{eff}}$ from the effective equilibrium constant using the standard thermodynamic relation

$$\Delta G_{\text{eff}} = -\ln\left( \frac{1}{n_h n_{ah}} e^{-\Delta G^0} \sum_{i=1}^{n_h} \sum_{j=1}^{n_{ah}} \sum_k e^{-n_{ijk}\Delta G} \right) \tag{S11}$$

$$\Delta G_{\text{eff}} = \Delta G^0 - \ln\left( \frac{1}{n_h n_{ah}} \sum_{i=1}^{n_h} \sum_{j=1}^{n_{ah}} \sum_k e^{-n_{ijk}\Delta G} \right). \tag{S12}$$

Akin to our definition of the change in Gibbs free energy of binding for the configurations $Z_{ijk}$ in Equation S2, we now define

$$\Delta G_{\text{eff}} = \Delta G^0 + n_{\text{eff}}\, \Delta G, \tag{S13}$$

where $n_{\text{eff}}$ is the effective interaction bond count of the parasitic interactions. Comparing this to Equation S12 yields the definition of our loss function:

$$n_{\text{eff}} = \frac{1}{-\Delta G} \ln\left( \frac{1}{n_h n_{ah}} \sum_{i=1}^{n_h} \sum_{j=1}^{n_{ah}} \sum_k e^{-\, n_{ijk}\Delta G} \right) = \text{Loss} \tag{S14}$$

where the product $n_h n_{ah}$ is just the number of pairs $N_{pair}$ from Equation S1.

Lastly, we can restate the effective equilibrium constant using the new definitions:

$$K_{\text{eff}} = \frac{[Z]}{[X][Y]} = e^{-(\Delta G^0 + n_{\text{eff}} \Delta G)}. \tag{S15}$$

#### 1.4 Generalization to Multi-layer Structures

The toy model described above applies only to two-layer structures, where one group of slats carries only plus-handles and the other carries only minus-handles. In multilayer crisscross structures, however, a single slat can have plus-handles on one side (for example, on the top helix) and minus-handles on the other (for example, on the bottom helix). Such slats can therefore take part in parasitic interactions with both plus-handle bearing and minus-handle–bearing slats at the same time.

To extend the model, we keep the idea that the first step of parasitic multimerization is the formation of a plus:minus bond. However, instead of describing binding between whole slats, we now describe binding between individual binding sites. In this version, we interpret $X_i$ and $Y_j$ as the concentrations of free plus-handle sites and free minus-handle sites, respectively. The indices $i$ and $j$ label specific sites on specific slats — for example, $X_i$ could be the handle site on slat $p$.

With this reinterpretation, the derivation above remains unchanged. The only difference is in how the quantities are understood: $X$ and $Y$ are now the total concentrations of free plus-handle and minus-handle sites in the pool, and $Z$ represents the total number of bonds formed between plus-handle and minus-handle sites. Importantly, the model is no longer restricted to physical dimers of whole slats. It now captures individual bonding events between sites.

#### 1.5 Histogram form

We note an alternative and computationally convenient representation of Equation S1, which is the form implemented in our software package. We define $n_\nu$ as the number of interactions in the pool with bond count $\nu$. In this representation, the loss, or effective parasitic interaction bond count, is given by

$$\text{Loss} = \frac{1}{-\Delta G} \ln\left( \frac{1}{N_{\text{Pairs}}} \sum_{\nu=1}^{\infty} n_\nu \, e^{-\,\nu \Delta G} \right). \tag{S16}$$

### 2 Supplementary Figures

The below figures contain example microscope images of the megastructures used for figures 4-7 in the main text. A much larger image dataset has been uploaded to Zenodo with DOI: https://doi.org/10.5281/zenodo.17914052. This dataset contains:

- Various images of each megastructure in Figure 4 at different magnifications.
- The entire set of images used for the quantitative results in figures 5-7.
- The data used to generate all three extended data figures.

All images in the Zenodo repository have been provided at full resolution in their raw format.

#### 2.1 Wide Field-of-View Images of Figure 4 Megastructure Gallery

The Loss, incubation time and imaging concentration for each sample has been provided in the caption for each figure. While the concentration can give some idea of the structure yield, deposition speed on a TEM micrograph is also dependent on a particle's physical dimensions and so it is likely that each structure will deposit at a different rate.

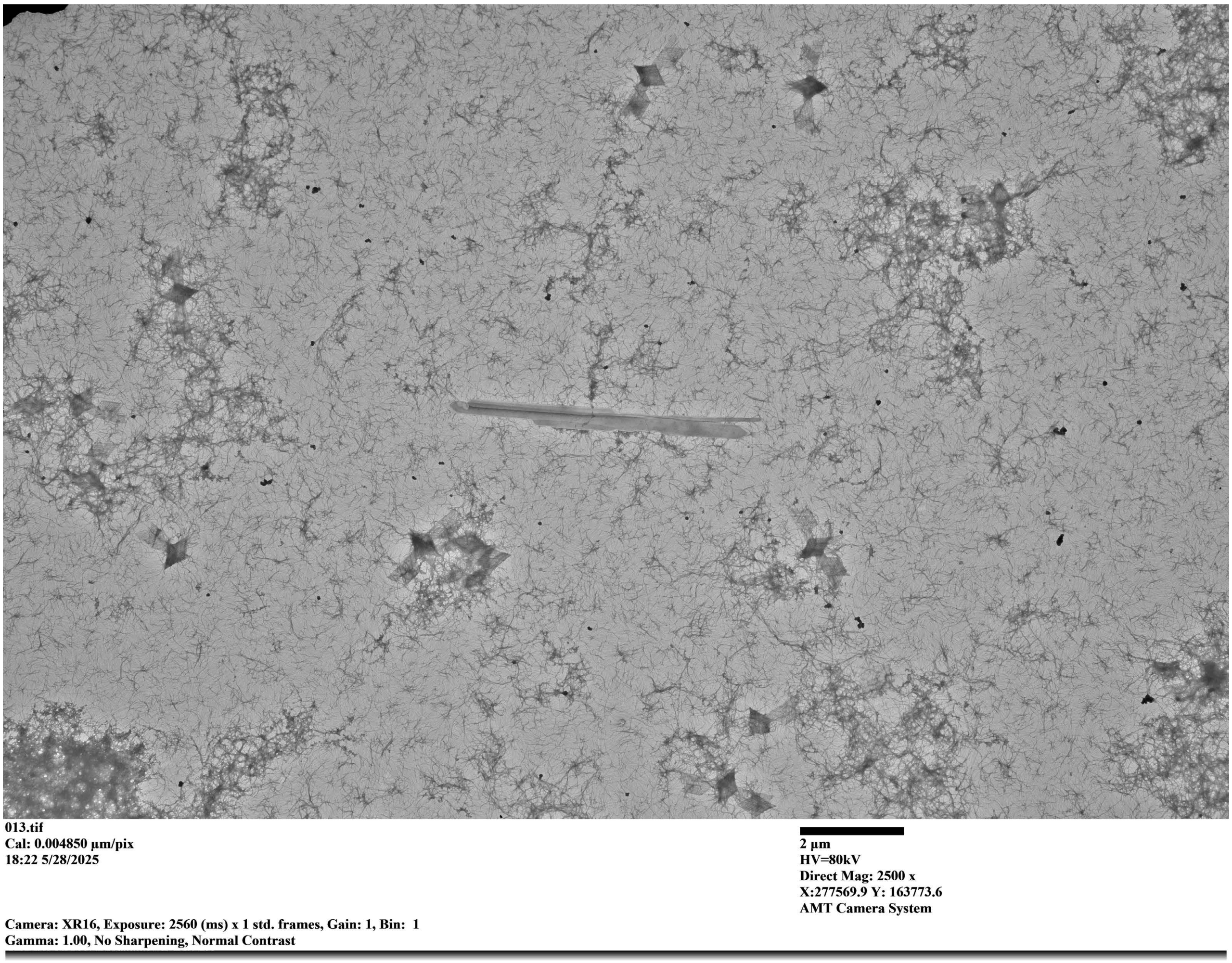

**Supp. Fig. 1**: Field-of-View TEM image of the ‘Bird’ design with a Loss of 4.46. Sample imaged at 60 pM seed concentration after 38 days of incubation. This long incubation time illustrates how megastructures can continue to assemble over long time periods while remaining stable at 37 ℃.

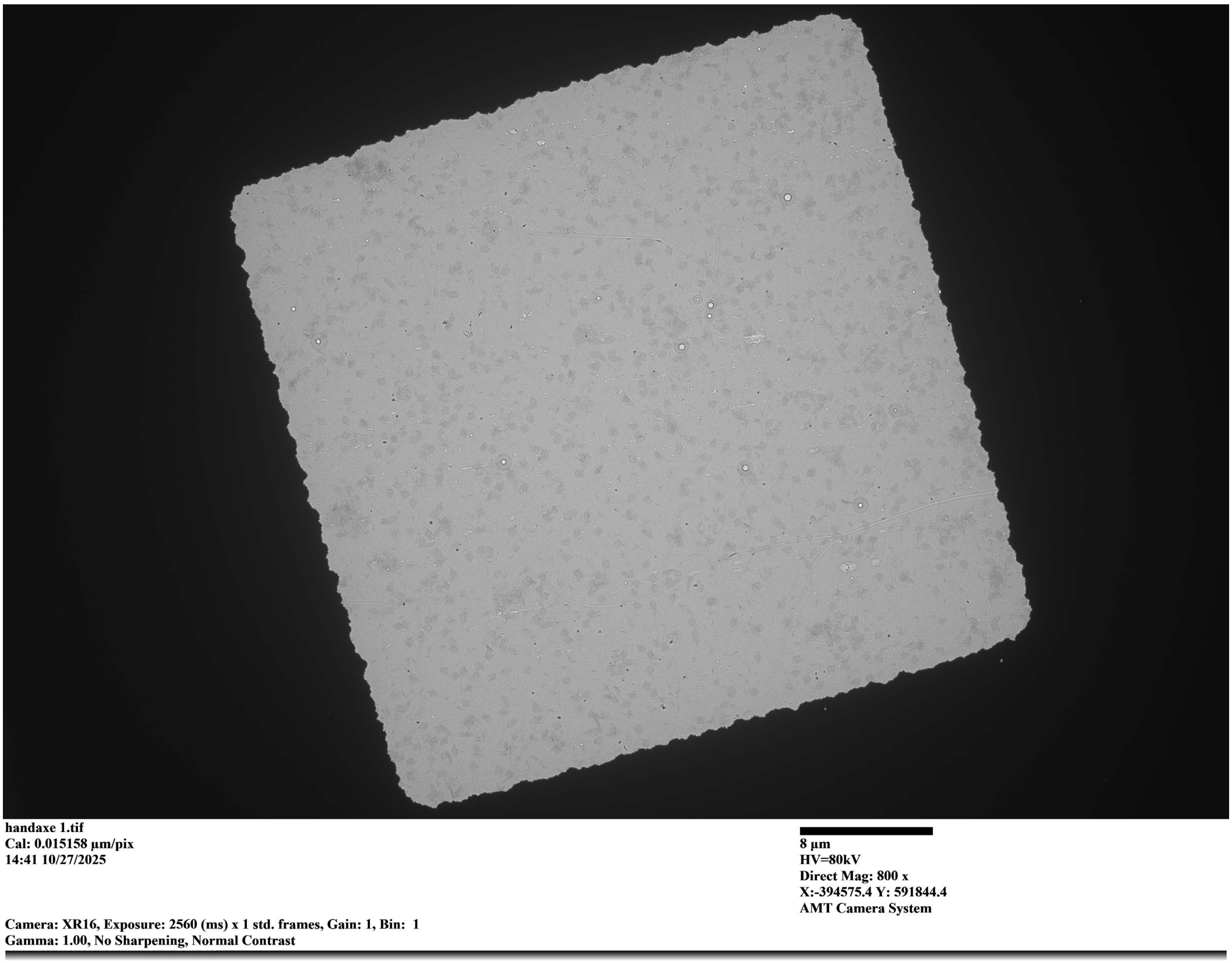

**Supp. Fig. 2**: Field-of-View TEM image of the ‘Handaxe’ design with a Loss of 2.06. Sample imaged at 500 pM seed concentration after 106 hours of incubation.

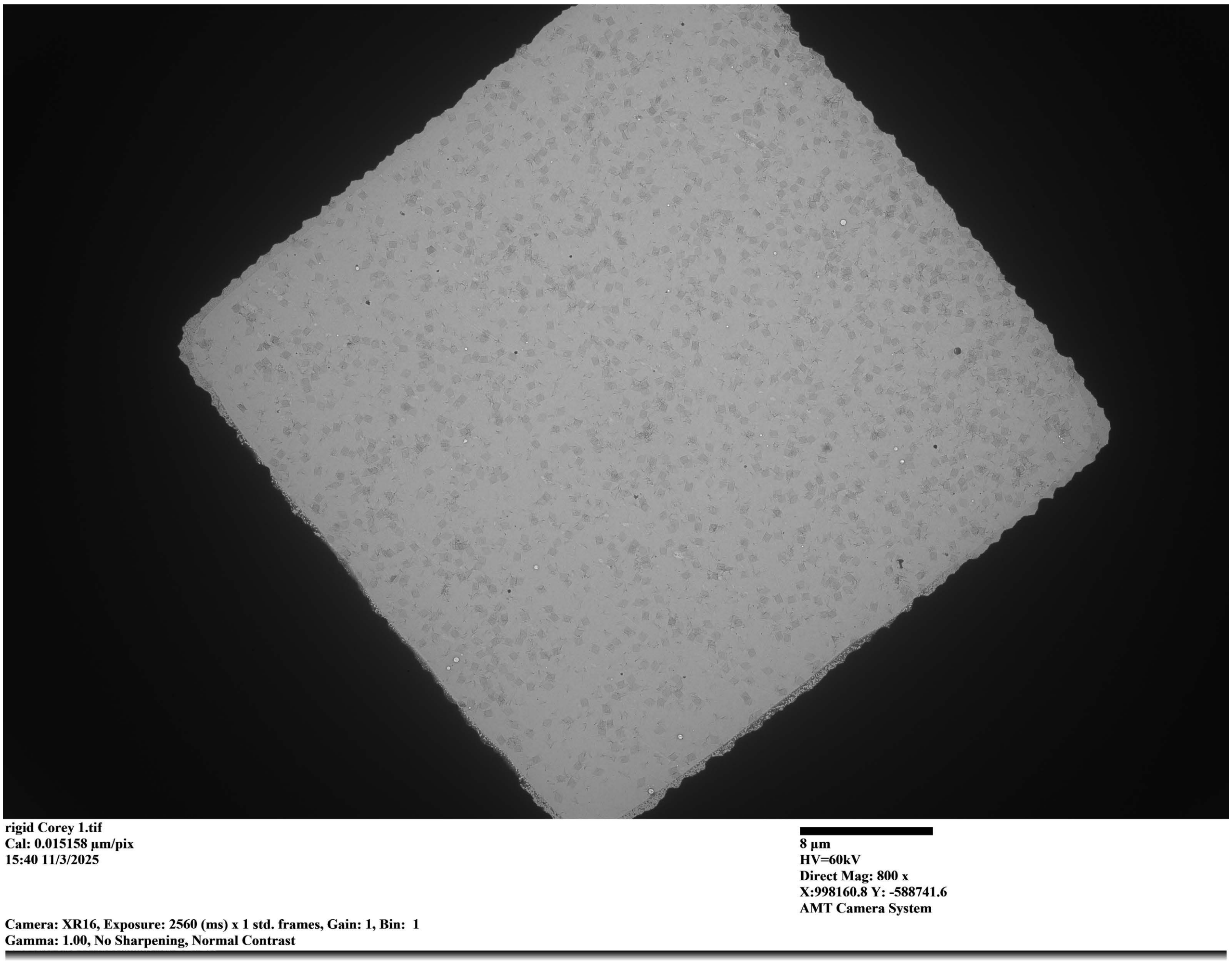

**Supp. Fig. 3**: Field-of-View TEM image of the ‘Rigid square’ design with a Loss of 2.84. Sample imaged at 123 pM seed concentration after 168 hours of incubation.

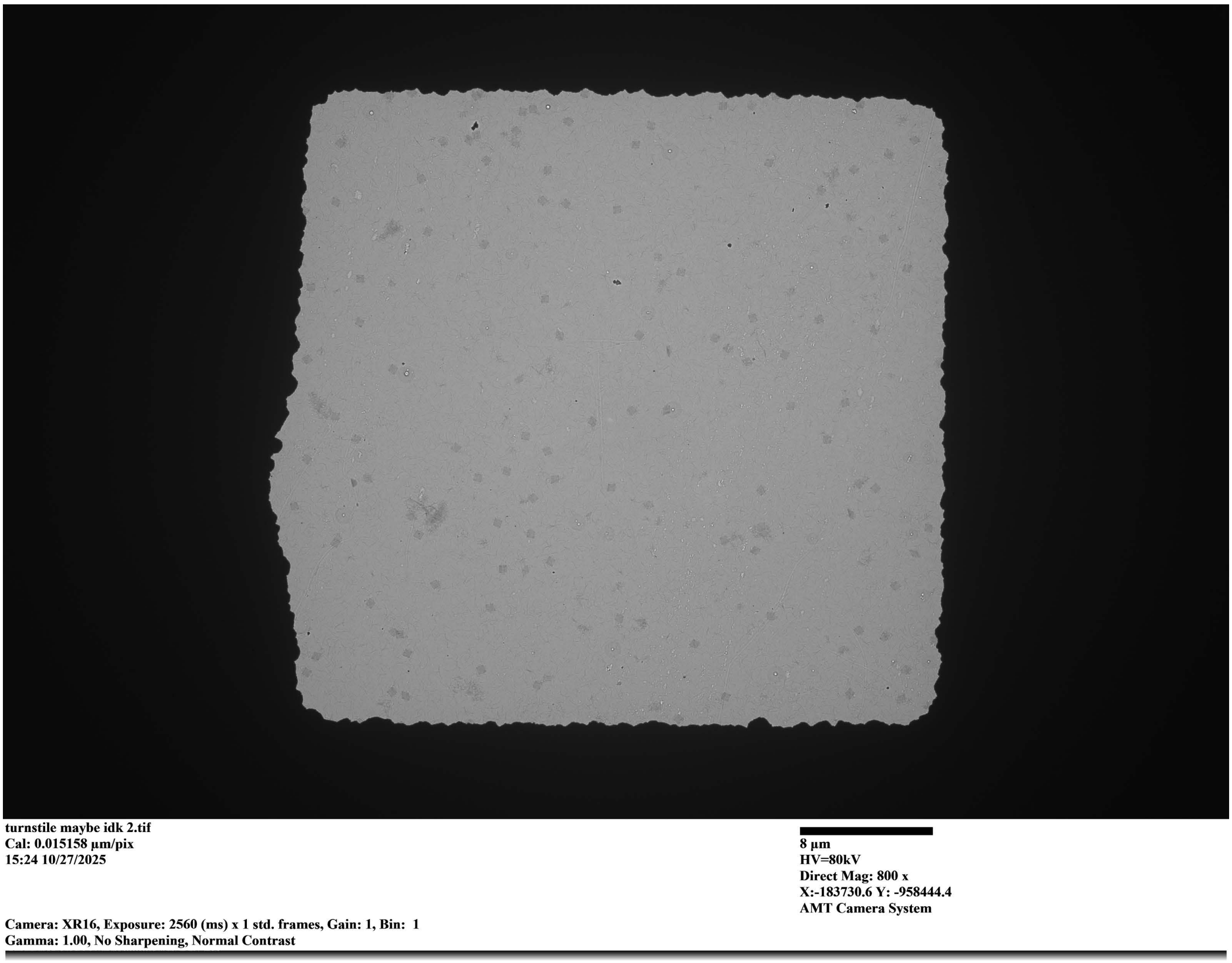

**Supp. Fig. 4**: Field-of-View TEM image of the ‘Turnstile’ design with a Loss of 2.76. Sample imaged at 250 pM seed concentration after 84 hours of incubation.

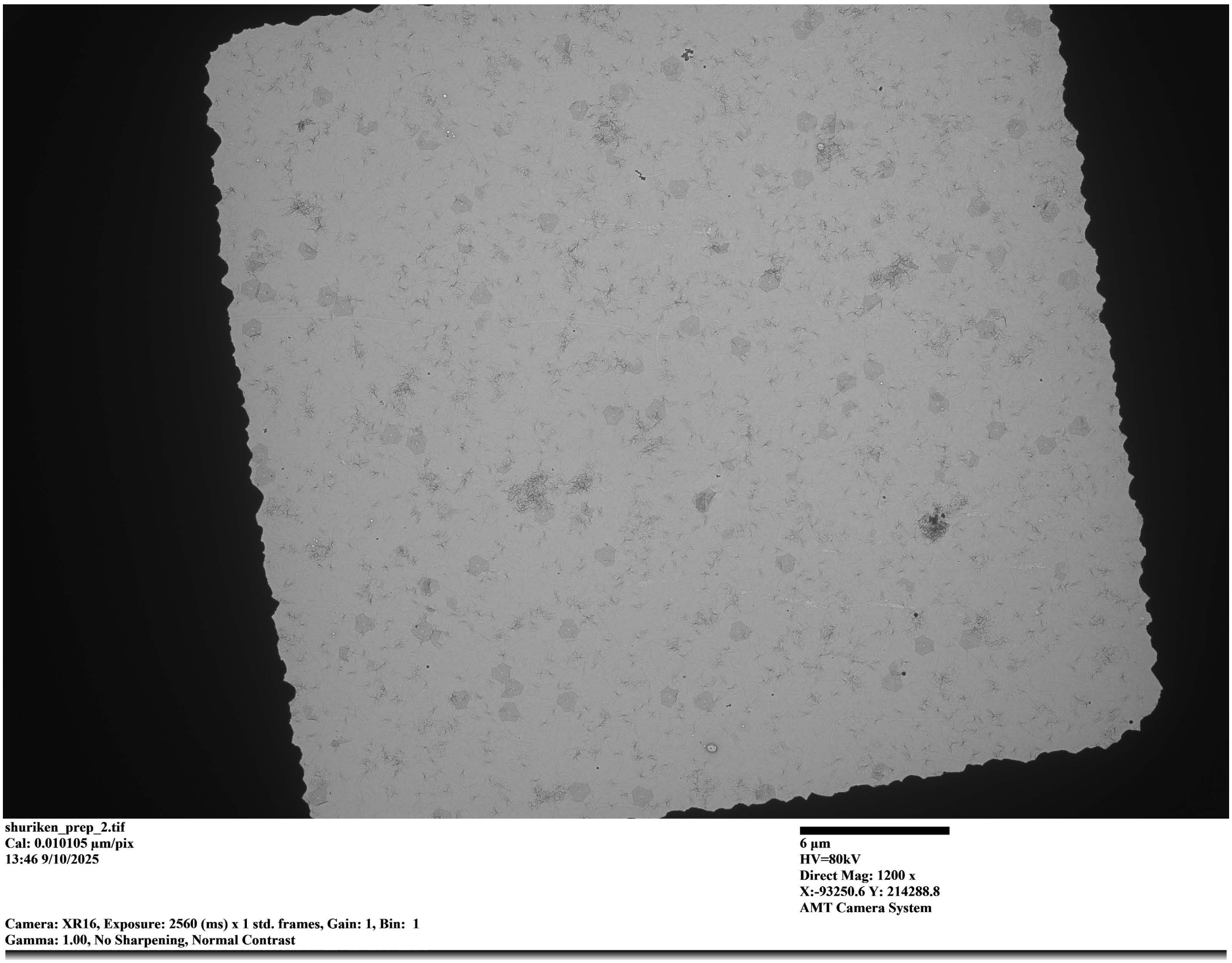

**Supp. Fig. 5**: Field-of-View TEM image of the 'Shuriken' design with a Loss of 2.7. Sample imaged at 93 pM seed concentration after 72 hours of incubation.

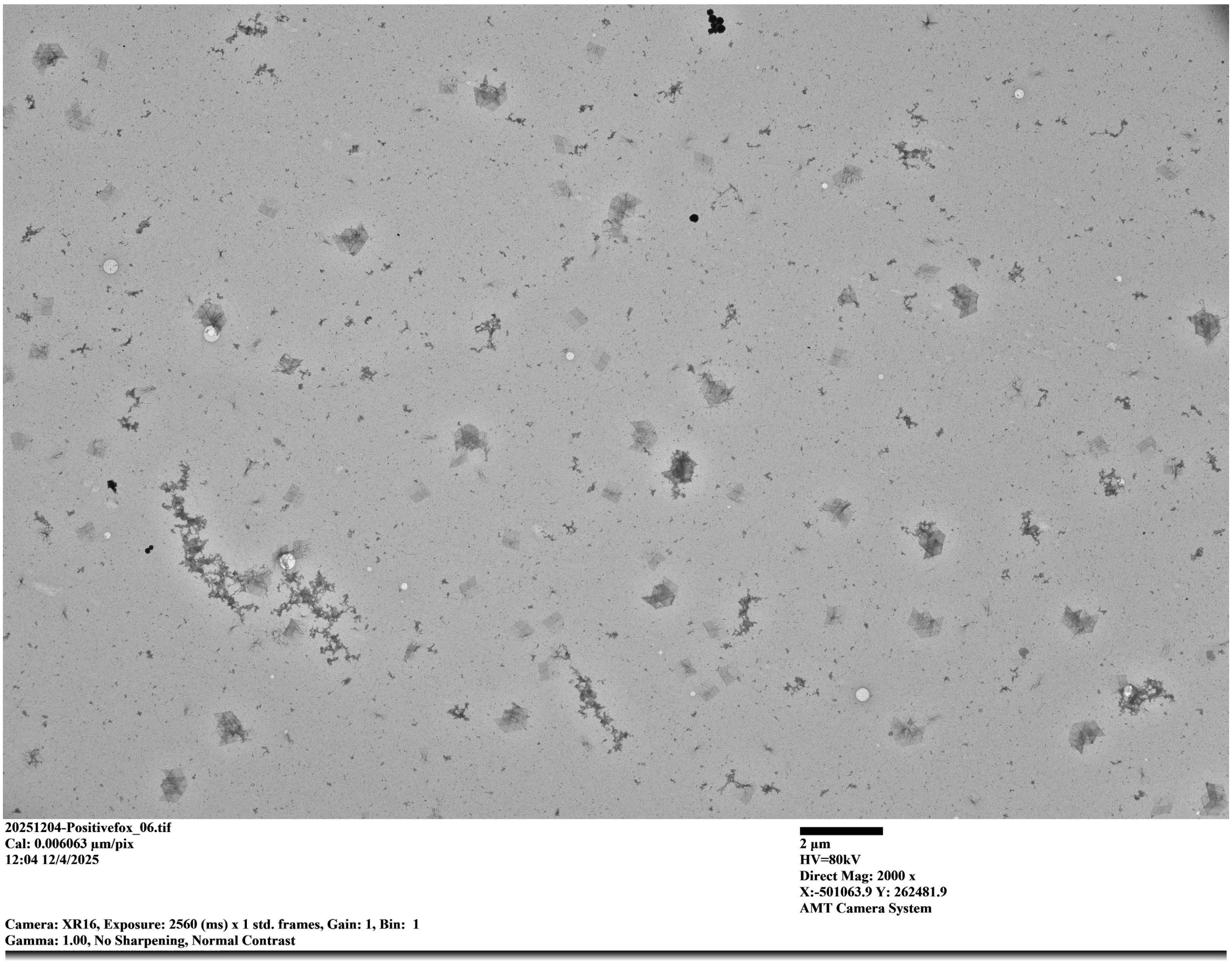

**Supp. Fig. 6**: Field-of-View TEM image of the ‘Fox’ design with a Loss of 2.73. Sample imaged at 70 pM seed concentration after 276 hours of incubation.

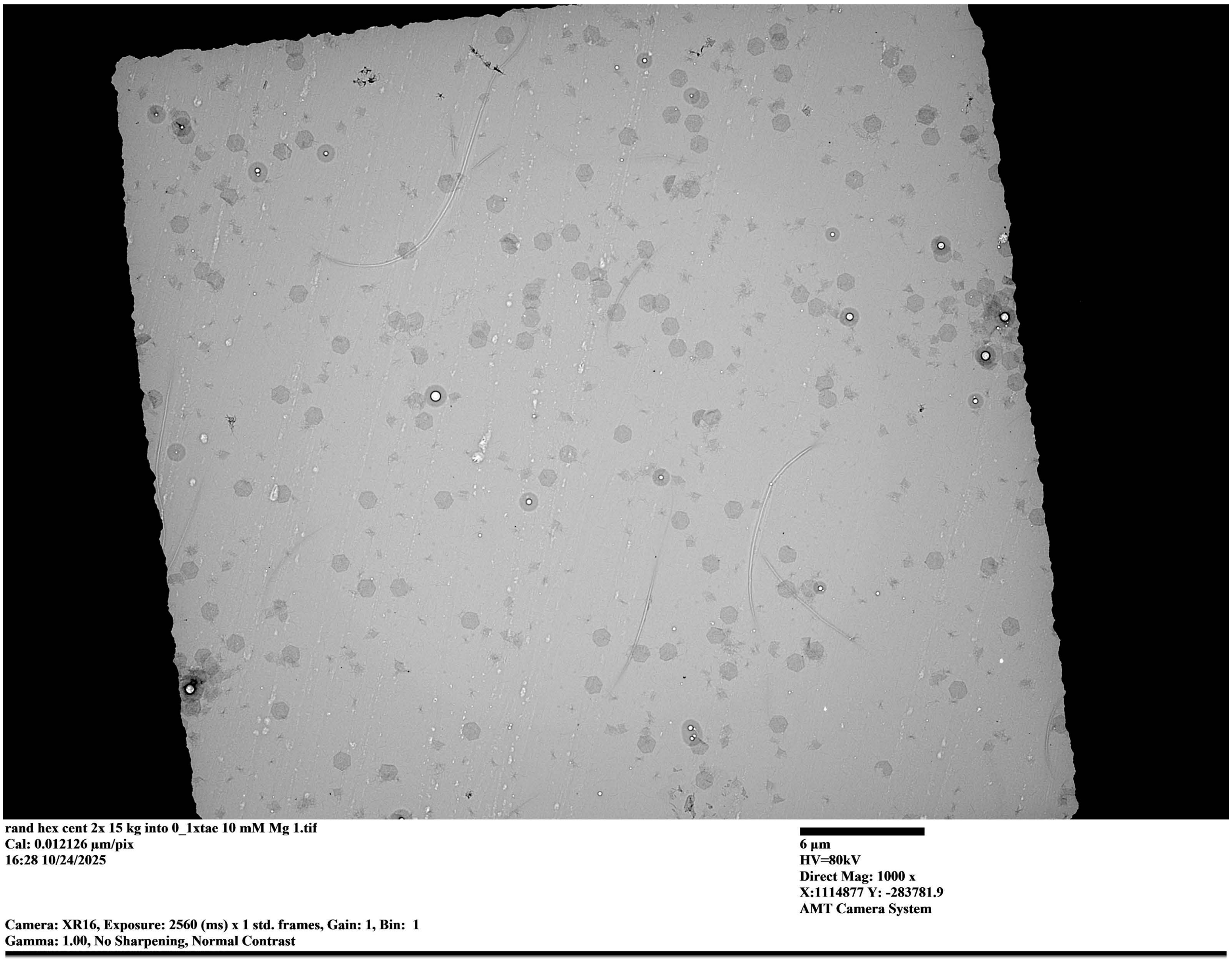

**Supp. Fig. 7**: Field-of-View TEM image of the 'Hexagon' design with a Loss of 2.7. Sample imaged at 350 pM seed concentration after 144 hours of incubation.

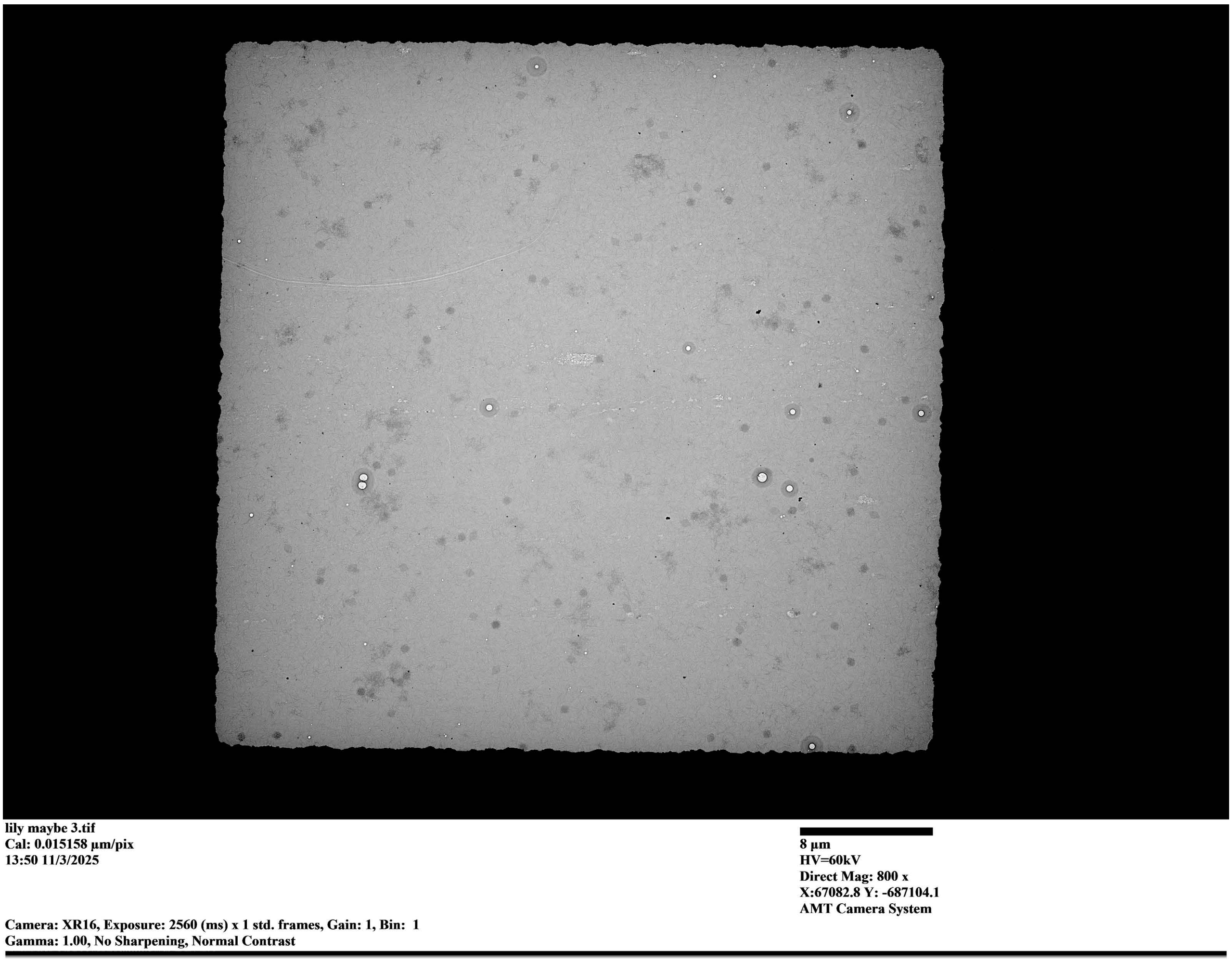

**Supp. Fig. 8**: Field-of-View TEM image of the ‘Lily’ design with a Loss of 3.8. Sample imaged at 250 pM seed concentration after 84 hours of incubation.

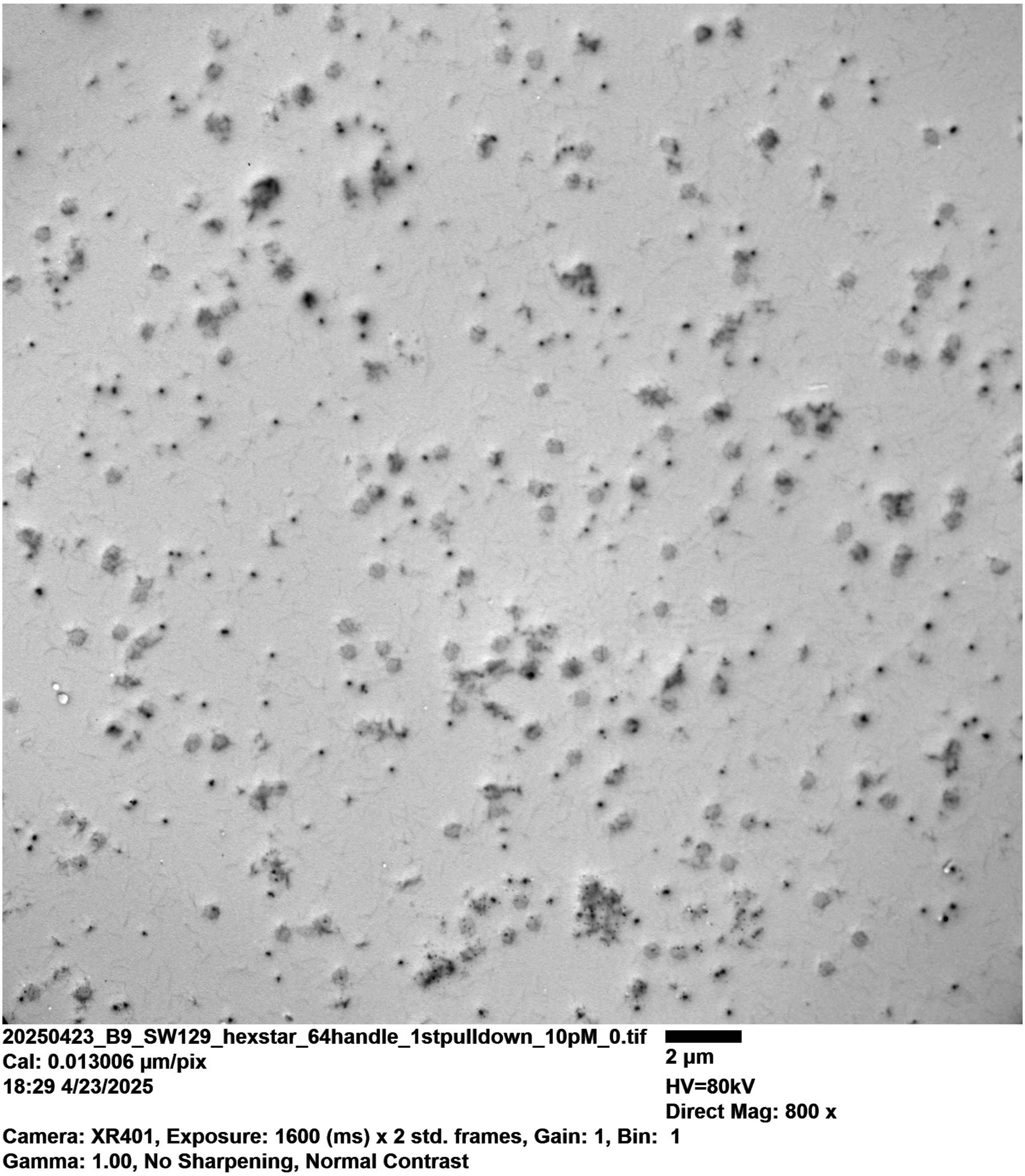

**Supp. Fig. 9**: Field-of-View TEM image of the ‘Daffodil’ design with a Loss of 2.62. Sample imaged at 10 pM seed concentration after 68 hours of incubation.

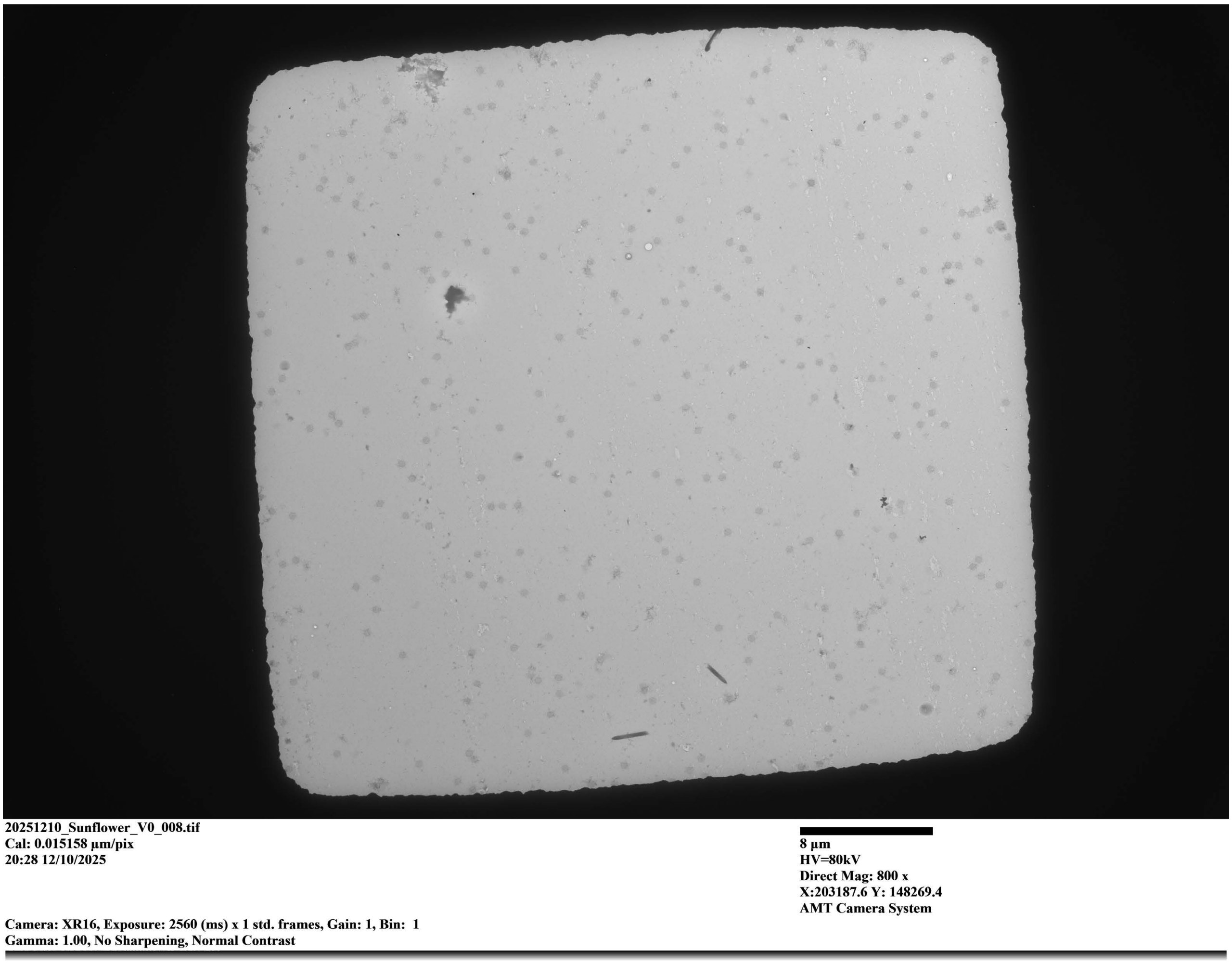

**Supp. Fig. 10**: Field-of-View TEM image of the 'Sunflower' design with a Loss of 3.6. Sample imaged at 130 pM seed concentration after 90 hours of incubation.

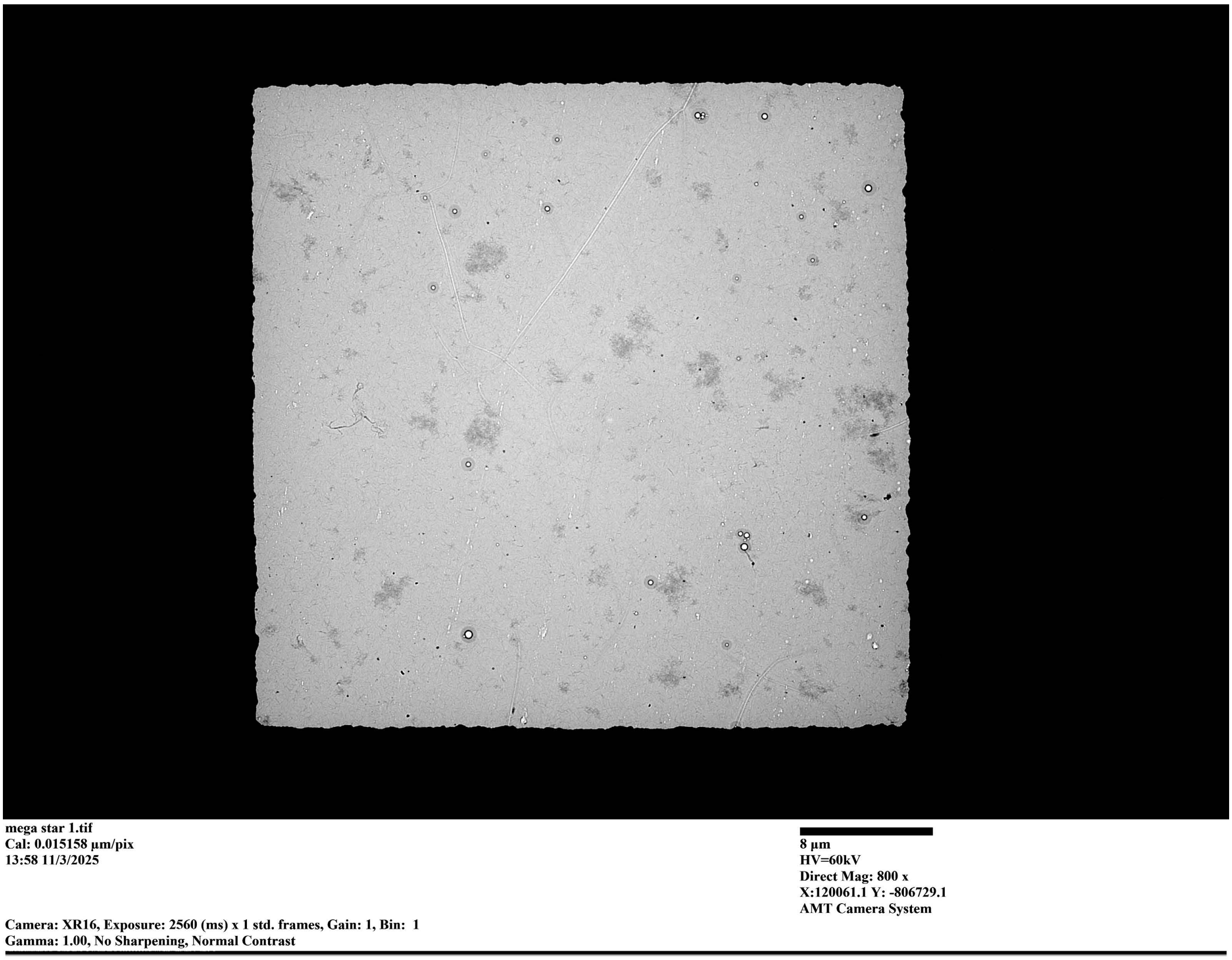

**Supp. Fig. 11**: Field-of-View TEM image of the ‘Megastar’ design with a Loss of 3.38. Sample imaged at 50 pM seed concentration after 106 hours of incubation.

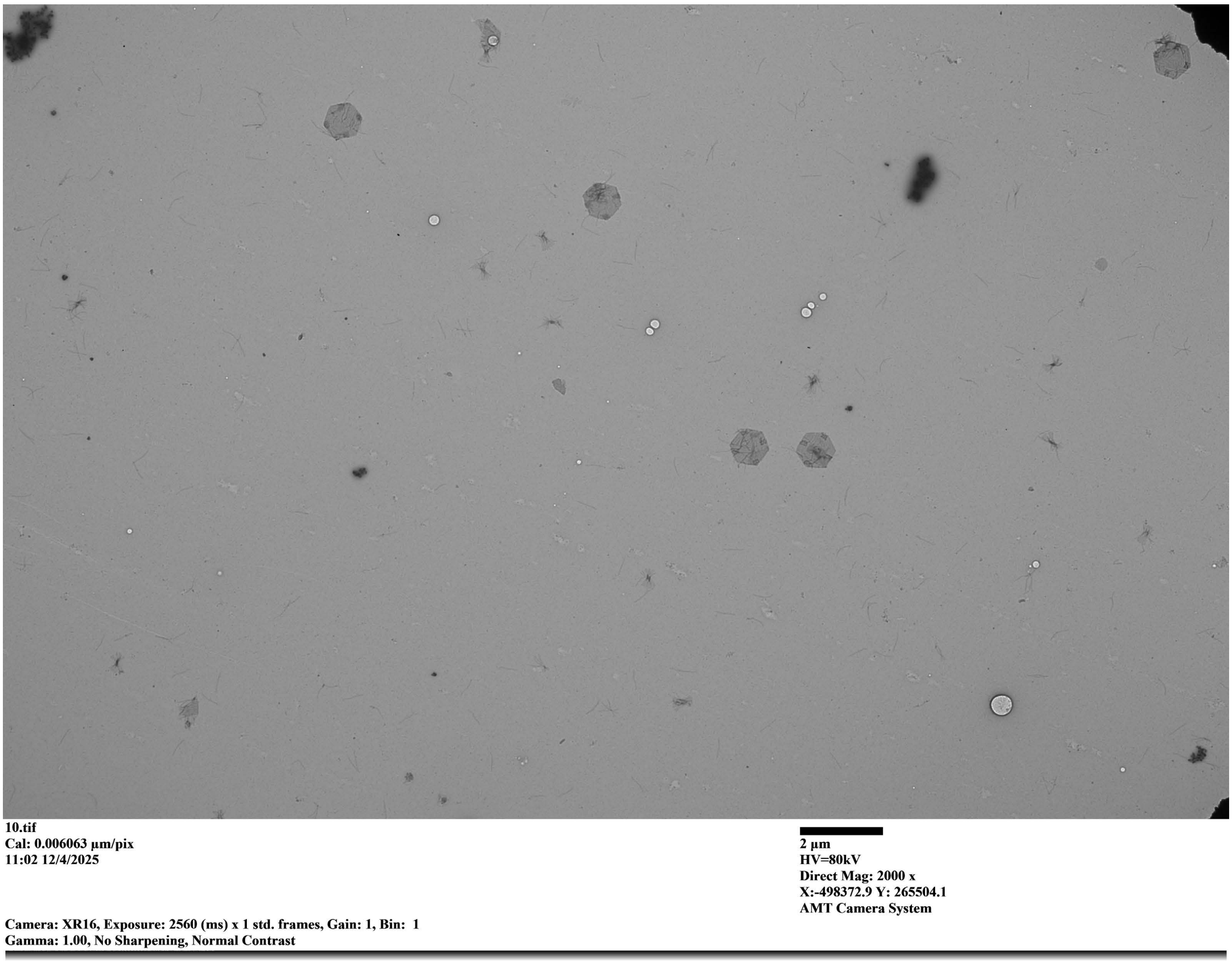

**Supp. Fig. 12**: Field-of-View TEM image of the ‘Hexagon’ with nanocubes attached. Sample imaged at 70 pM seed concentration after 140 hours of incubation.

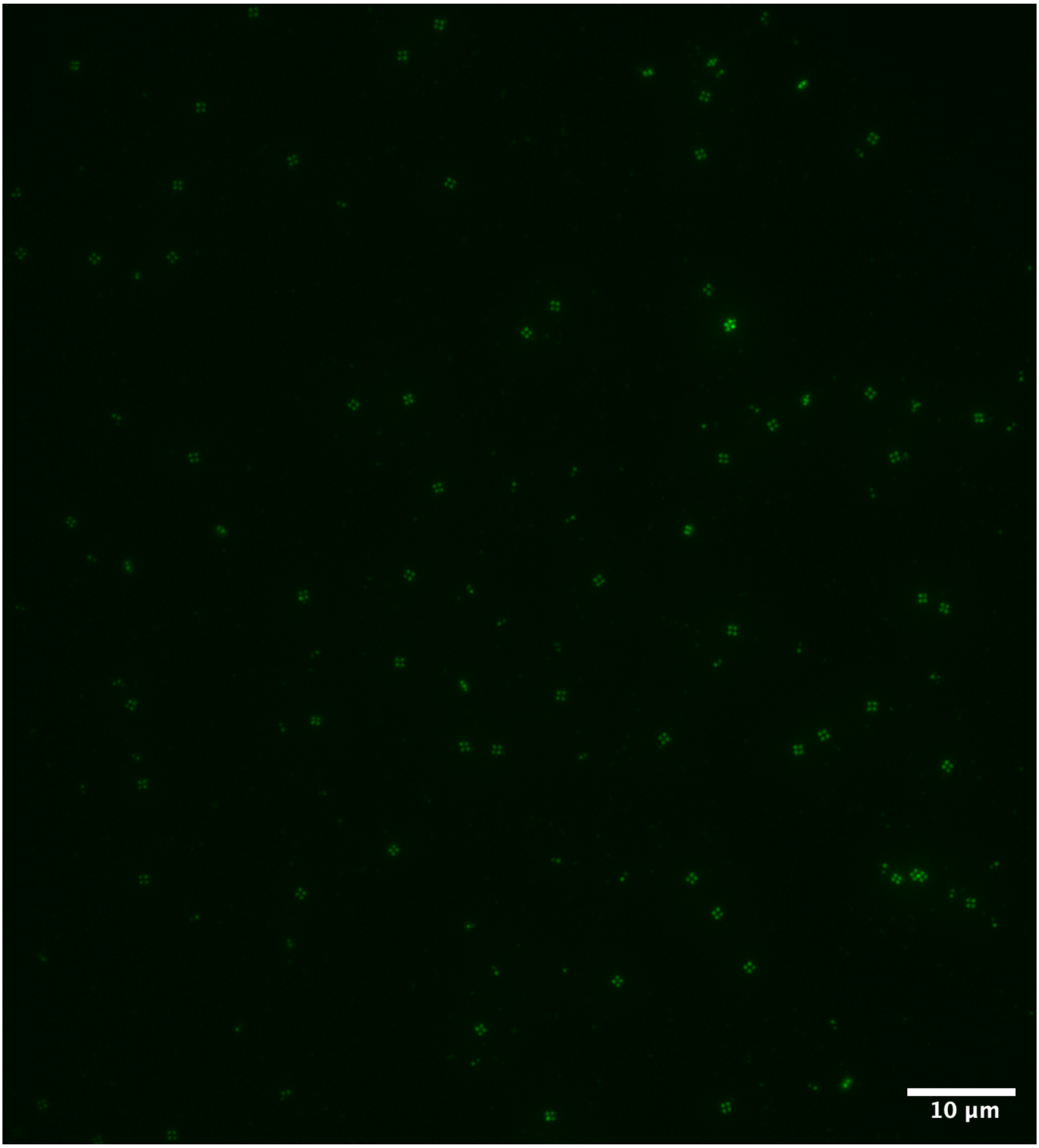

**Supp. Fig. 13**: Field-of-View fluorescent image of the ‘Hexagon’ with nanocubes attached (incubation time identical to that of Supp. Fig. 12).

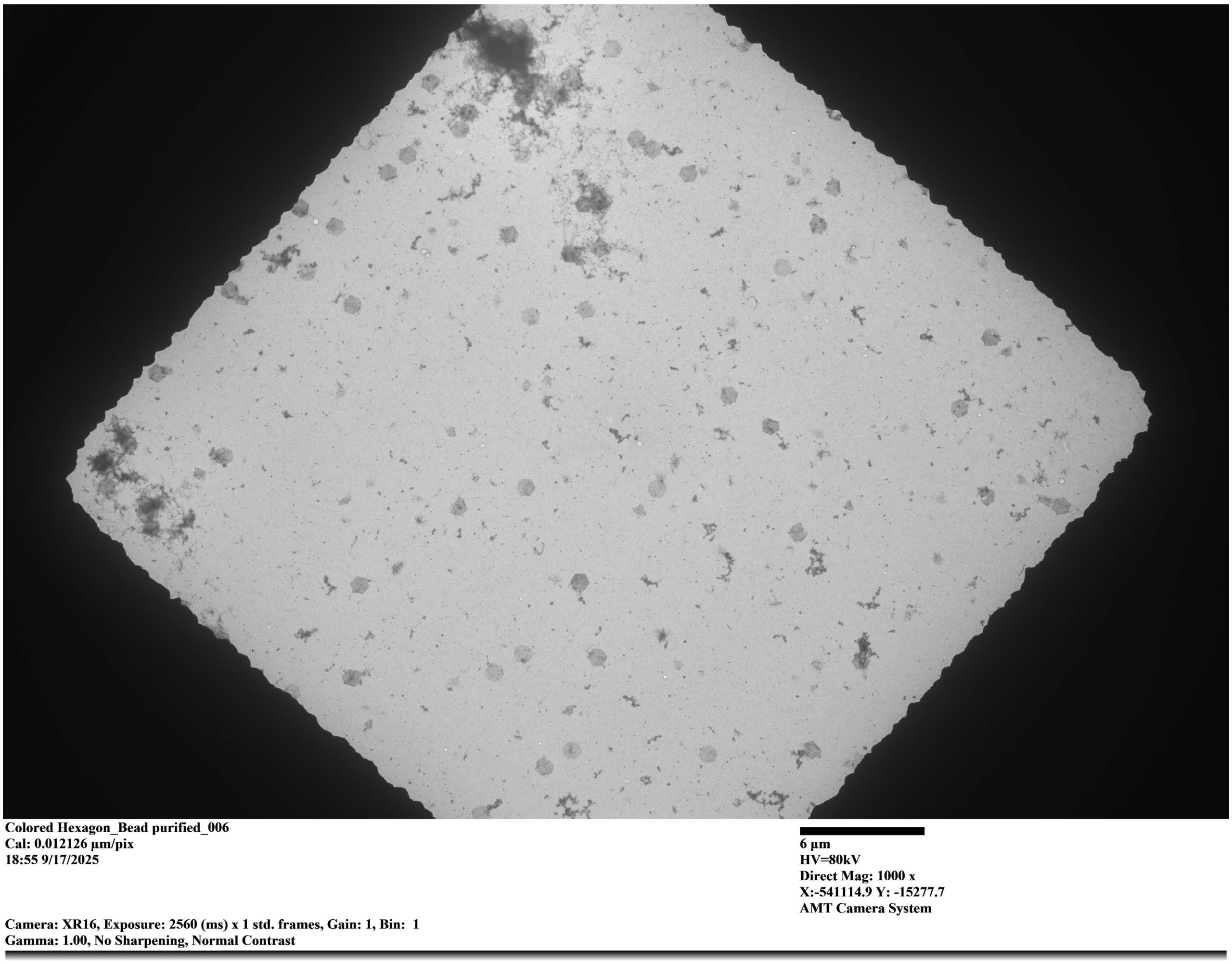

**Supp. Fig. 14**: Field-of-View TEM image of the ‘Hexagon’ with color-coded slats. Sample imaged at 70 pM seed concentration after 84 hours of incubation.

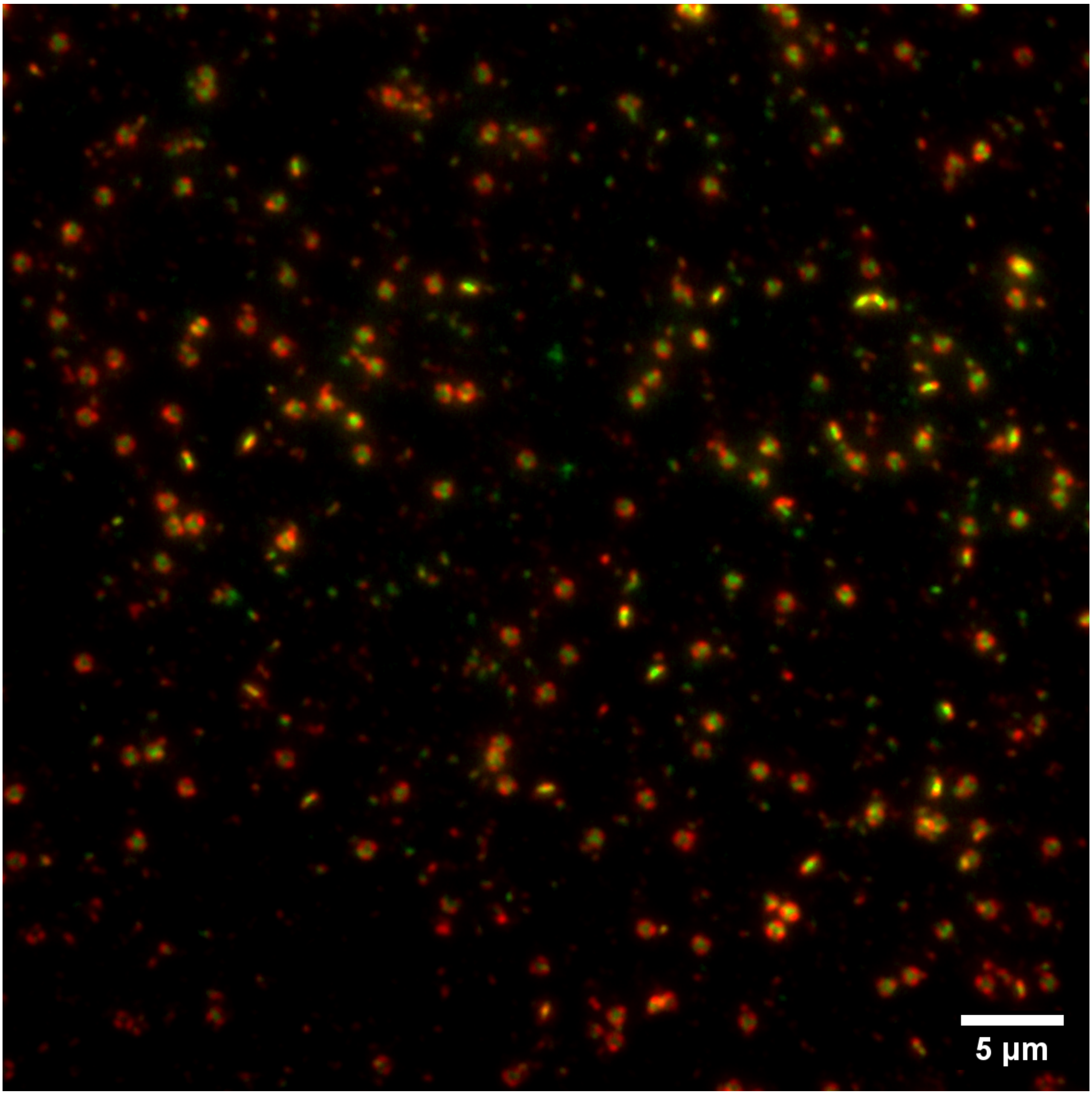

**Supp. Fig. 15**: Field-of-View fluorescent image of the 'Hexagon' with color-coded slats (incubation time identical to that of Supp. Fig. 14).

#### 2.2 Example TEM Images from Figure 5

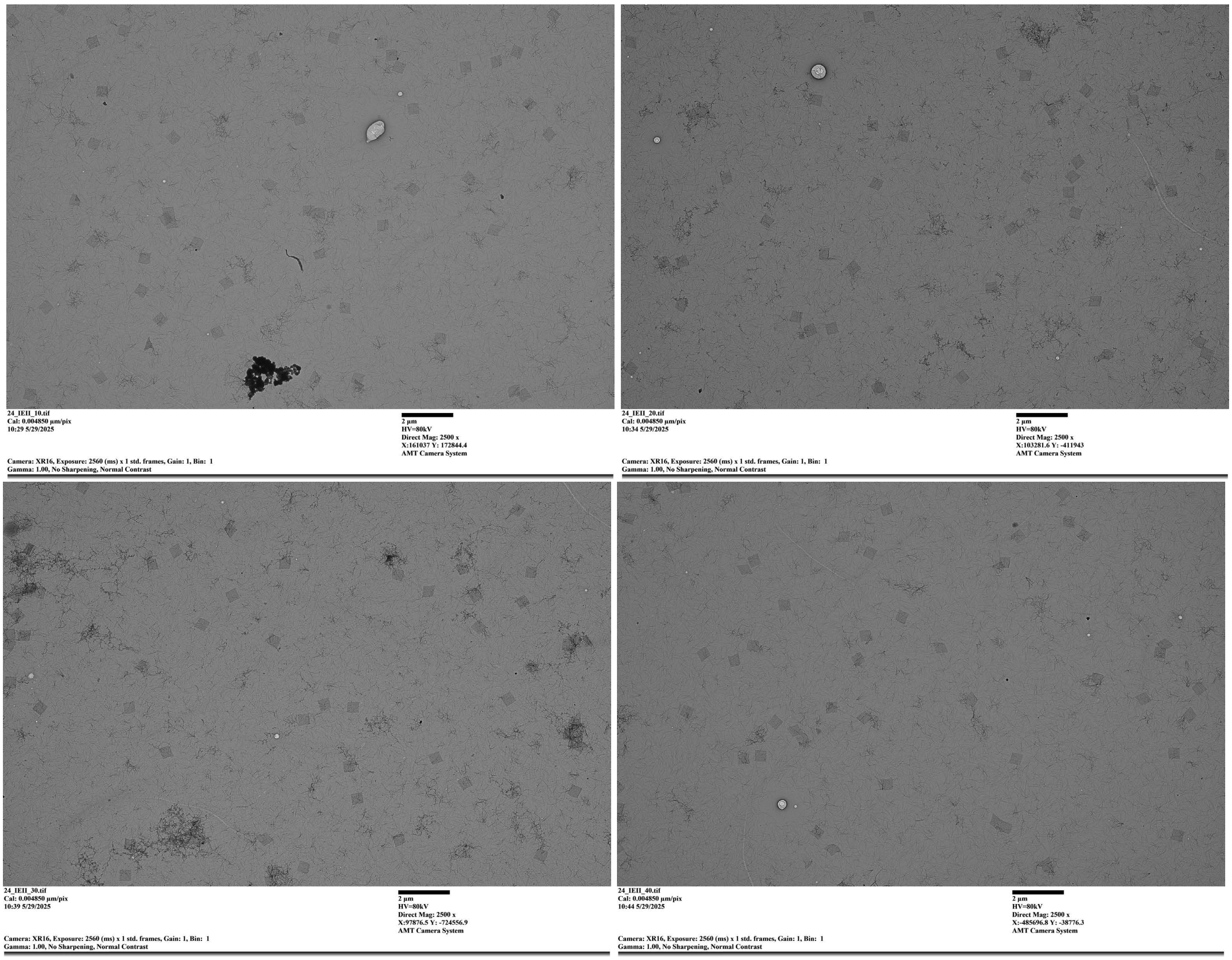

**Supp. Fig. 16**: Representative TEM images used to quantify the yield of the square megastructure design with **Loss = 7.3**. The corresponding yield data are shown in Figure 5.

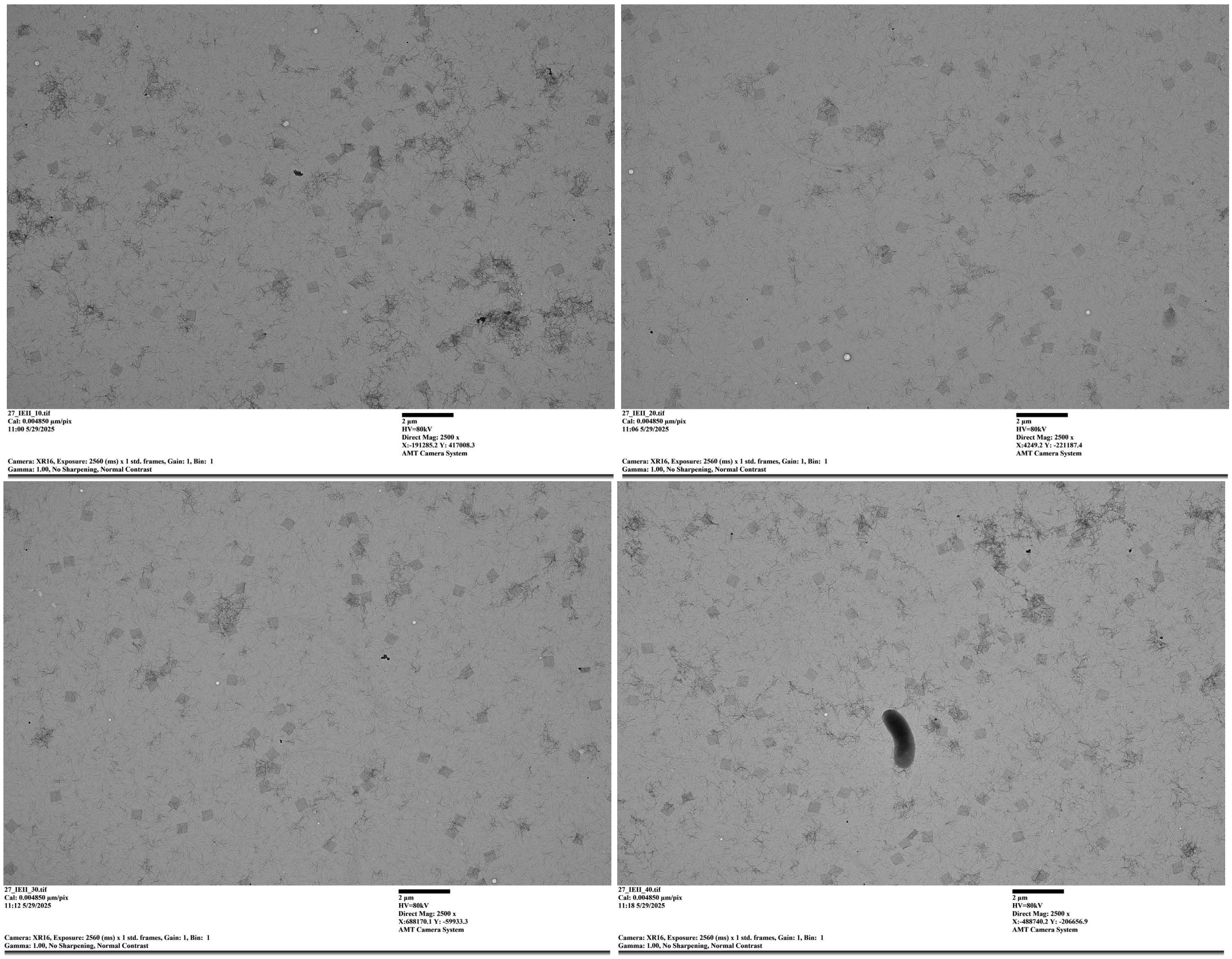

**Supp. Fig. 17**: Representative TEM images used to quantify the yield of the square megastructure design with **Loss = 4.7**. The corresponding yield data are shown in Figure 5.

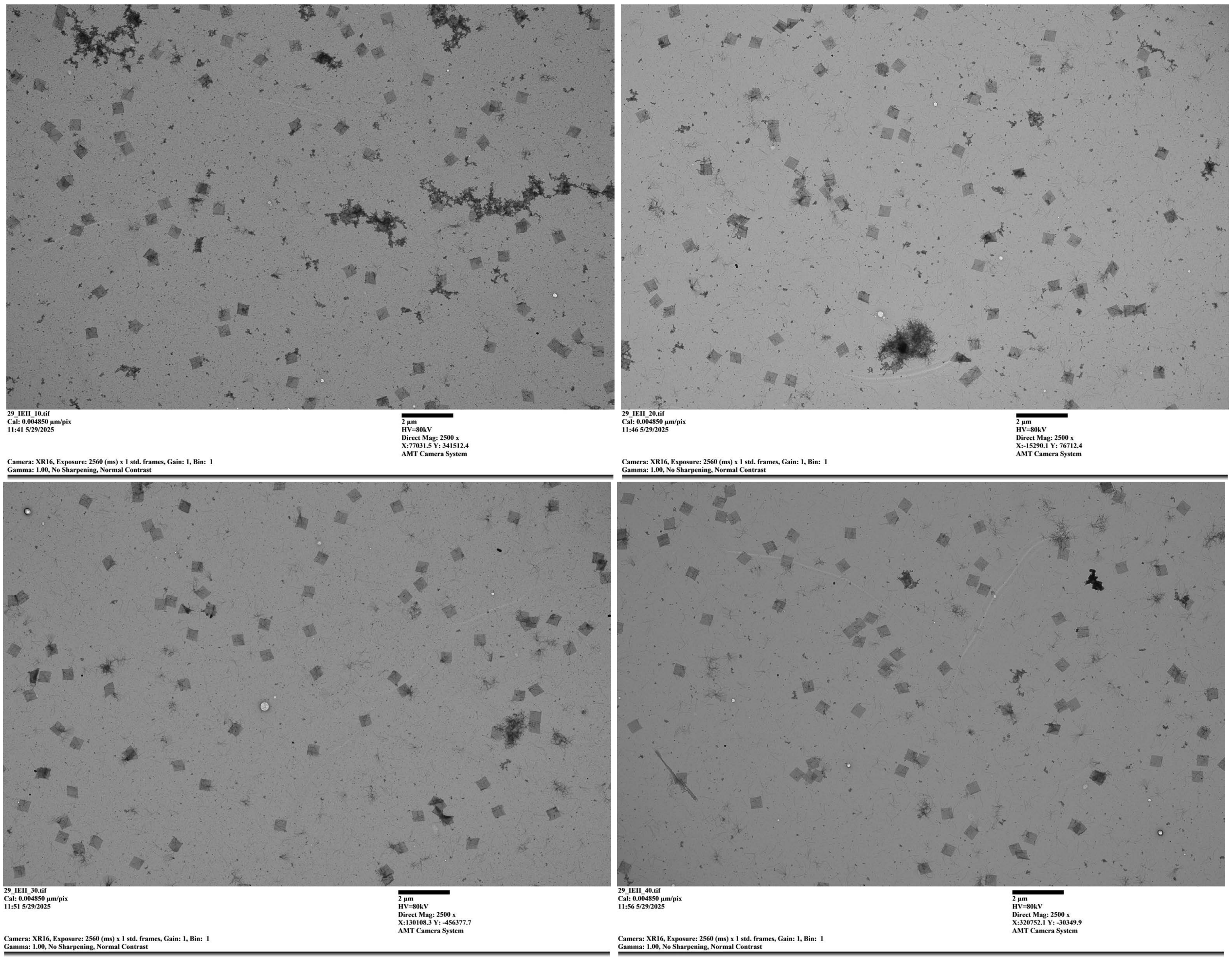

**Supp. Fig. 18**: Representative TEM images used to quantify the yield of the square megastructure design with **Loss = 2.6**. The corresponding yield data are shown in Figure 5.

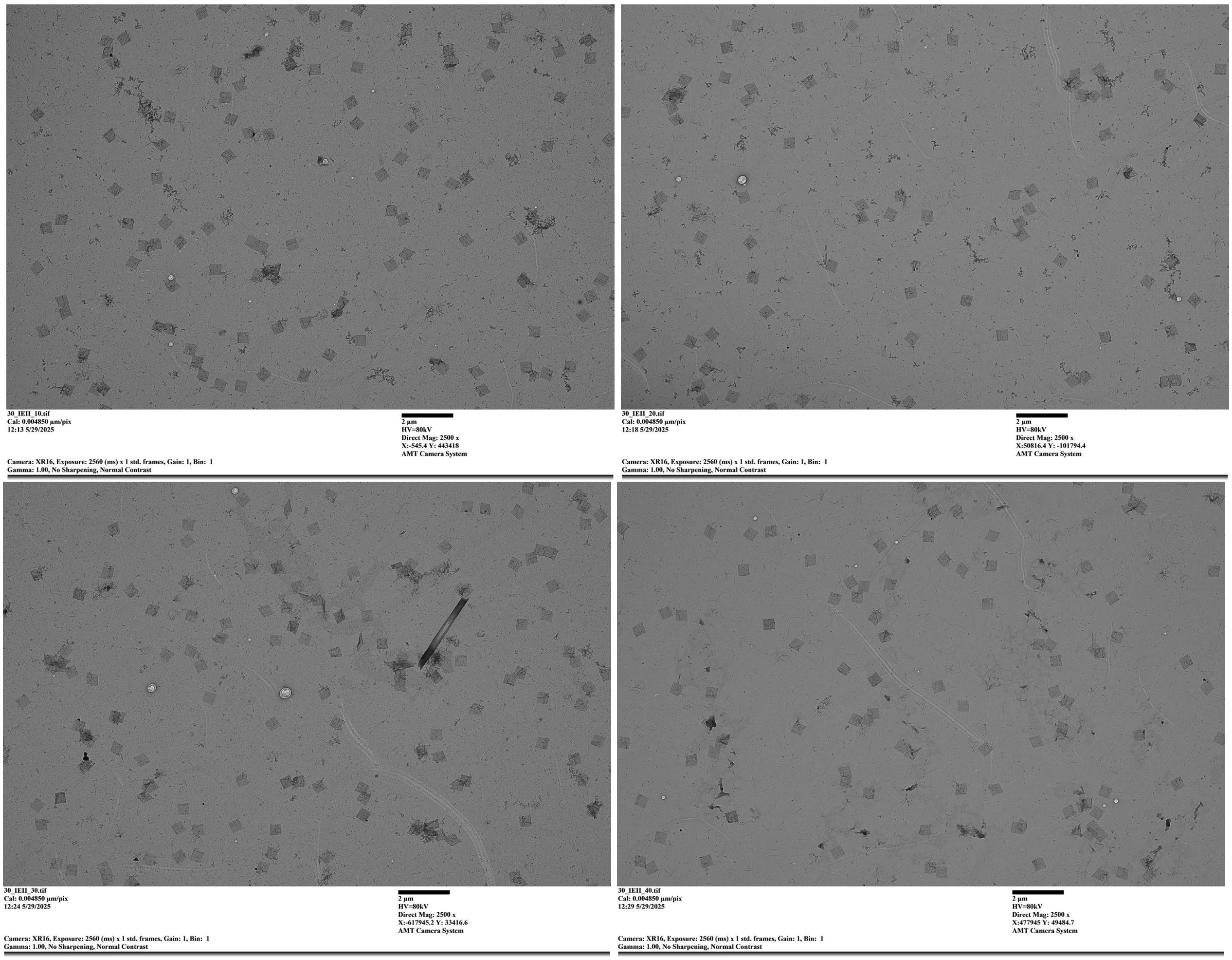

**Supp. Fig. 19**: Representative TEM images used to quantify the yield of the square megastructure design with **Loss = 2.1**. The corresponding yield data are shown in Figure 5 of the main text.

#### 2.3 Example TEM Images from Figure 6/7 (sunflowers)

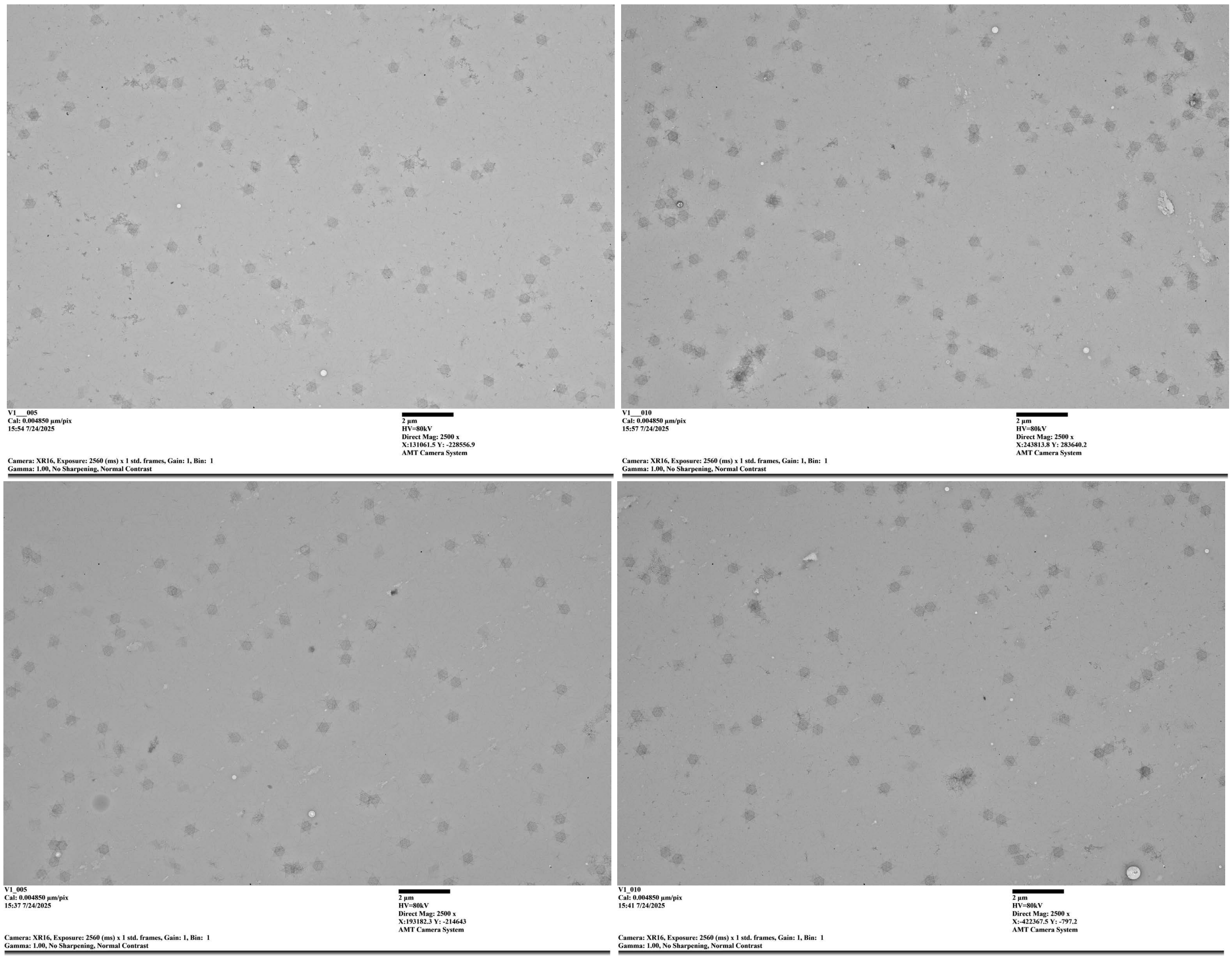

**Supp. Fig. 20**: Representative TEM images used to quantify the yield and structural completeness of the Sunflower megastructure design with **Loss = 2.8** using the **new** handle library. The corresponding yield data are shown in Figure 6 of the main text.

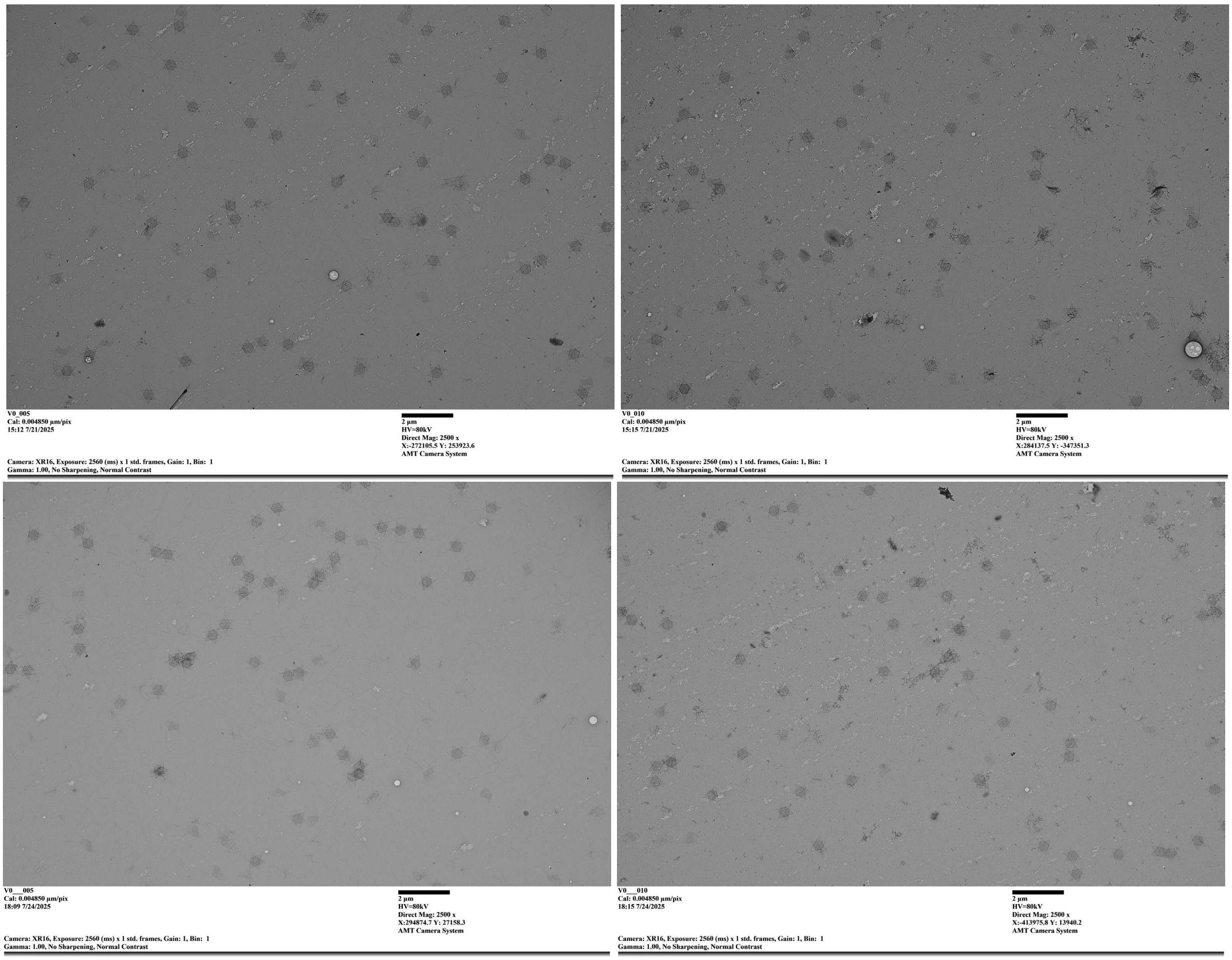

**Supp. Fig. 21**: Representative TEM images used to quantify the yield and structural completeness of the Sunflower megastructure design with **Loss = 3.6** using the **new** handle library. The corresponding yield data are shown in Figure 6 of the main text.

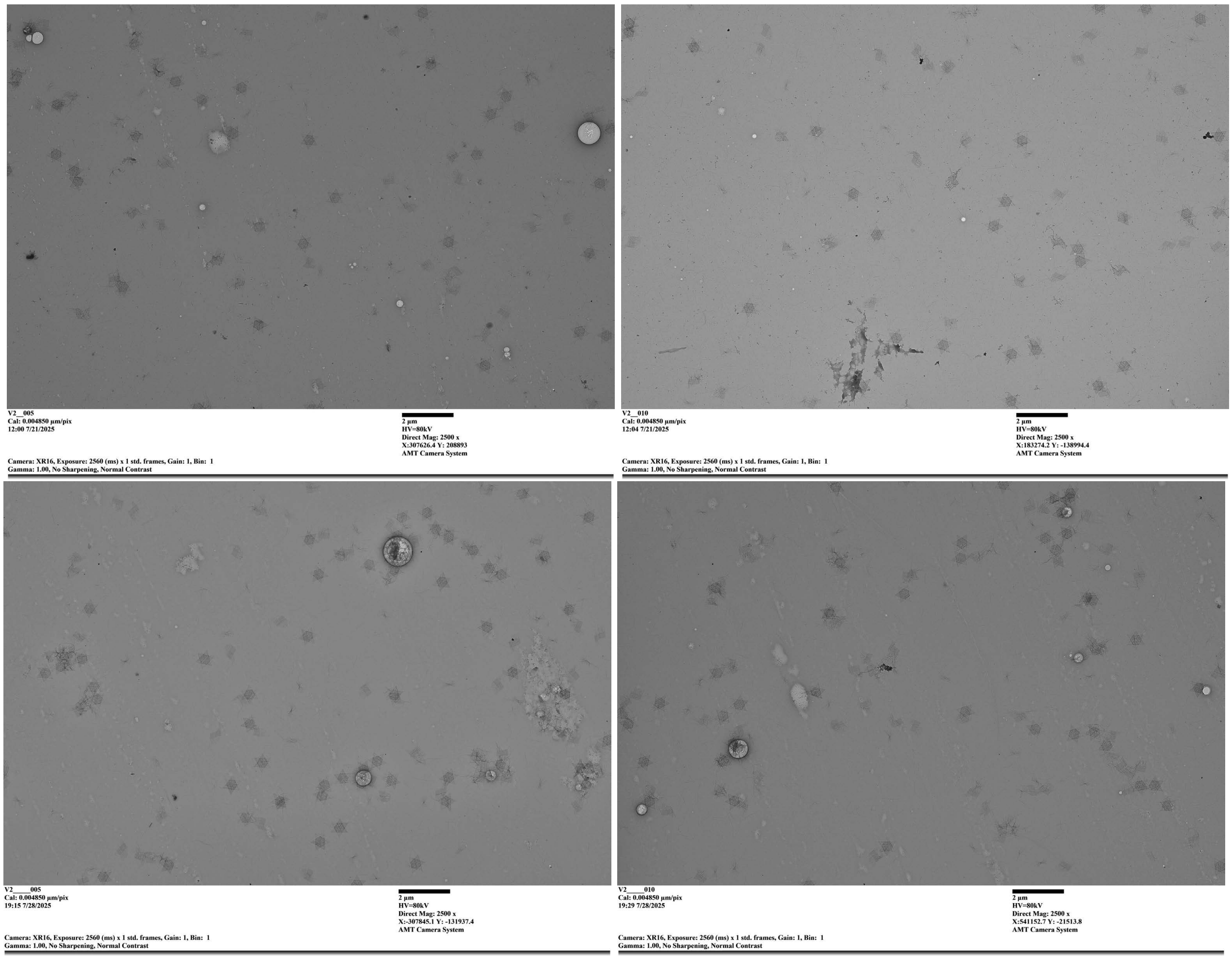

**Supp. Fig. 22**: Representative TEM images used to quantify the yield and structural completeness of the Sunflower megastructure design with **Loss = 3.8** using the **new** handle library. The corresponding yield data are shown in Figure 6 of the main text.

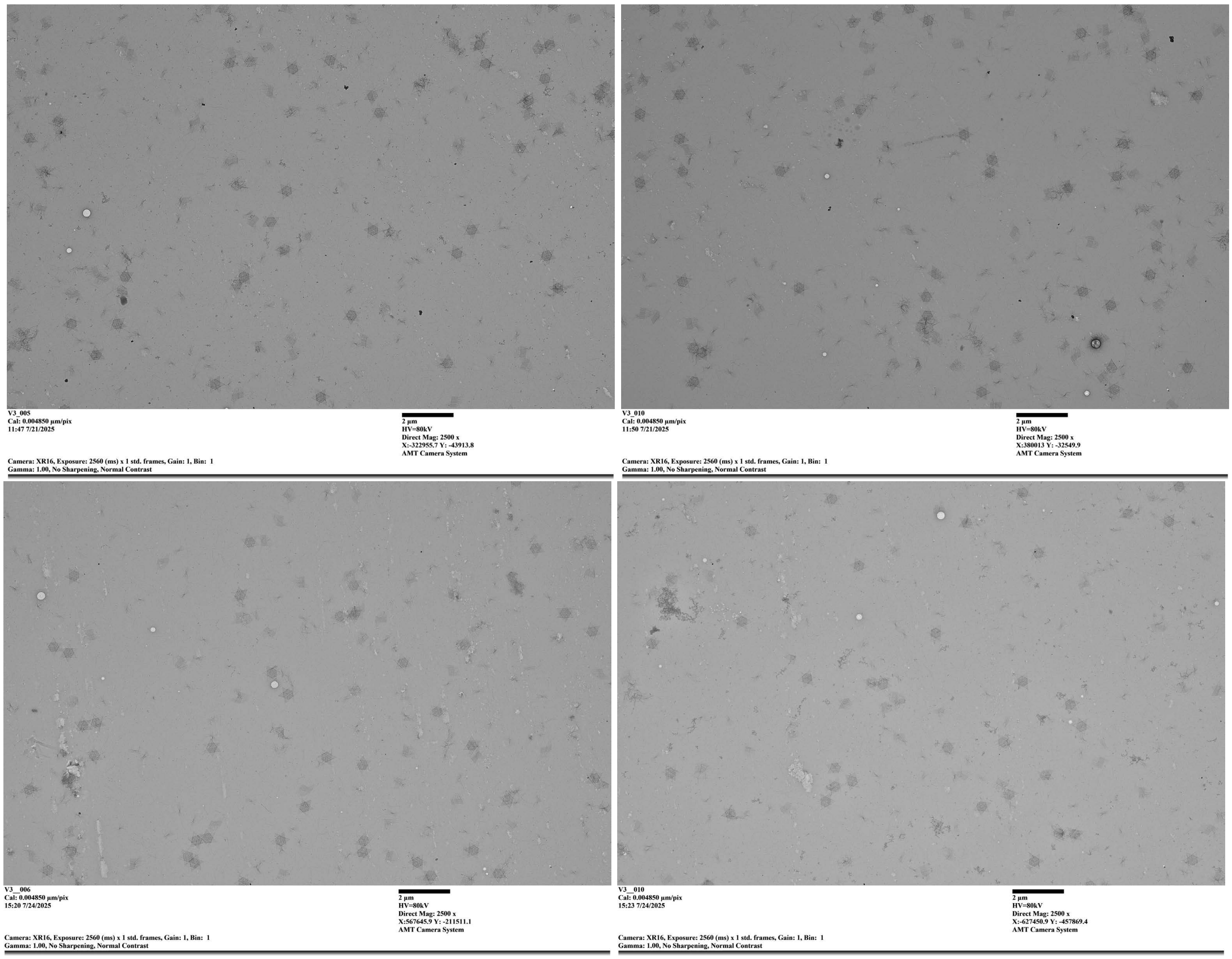

**Supp. Fig. 23**: Representative TEM images used to quantify the yield and structural completeness of the Sunflower megastructure design with **Loss = 3.8** using the **old** handle library. The corresponding yield data are shown in Figure 6 of the main text.

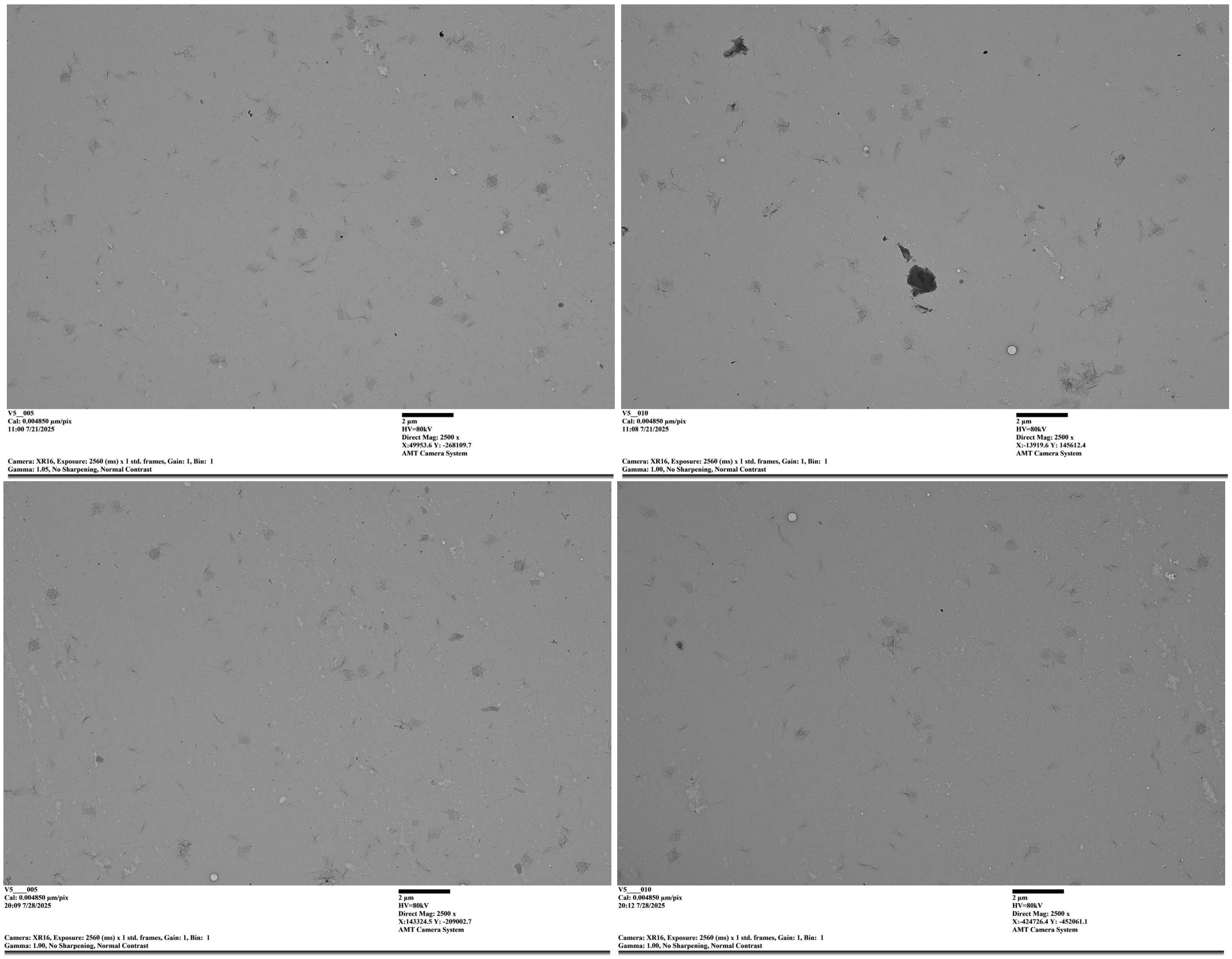

**Supp. Fig. 24**: Representative TEM images used to quantify the yield and structural completeness of the Sunflower megastructure design with **Loss = 4.7** using the **new** handle library. The corresponding yield data are shown in Figure 6 of the main text.

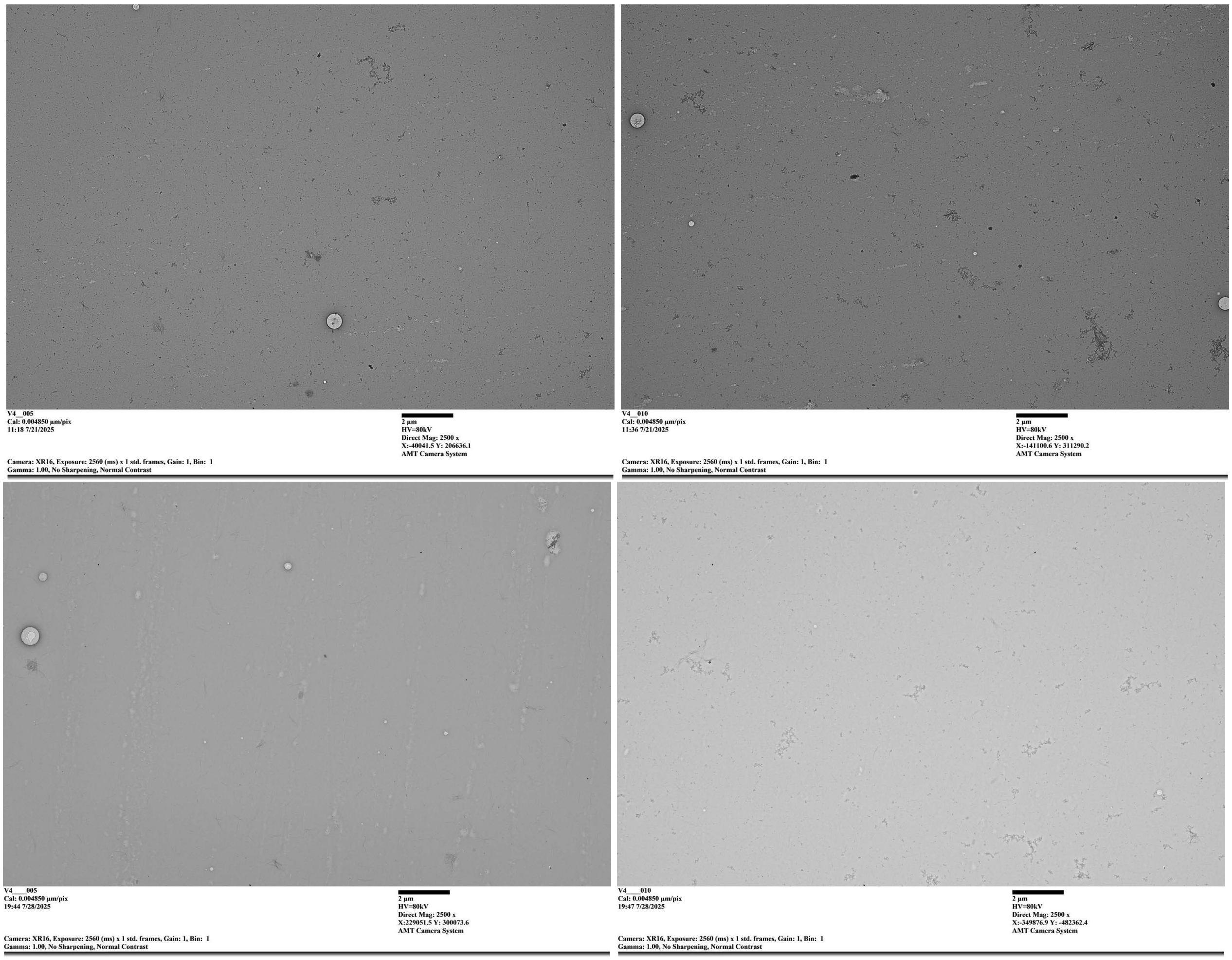

**Supp. Fig. 25**: Representative TEM images used to quantify the yield and structural completeness of the Sunflower megastructure design with **Loss = 5.1** using the **new** handle library. The corresponding yield data are shown in Figure 6 of the main text.

#### 2.4 Example TEM Images from Figure 7 (ribbon)

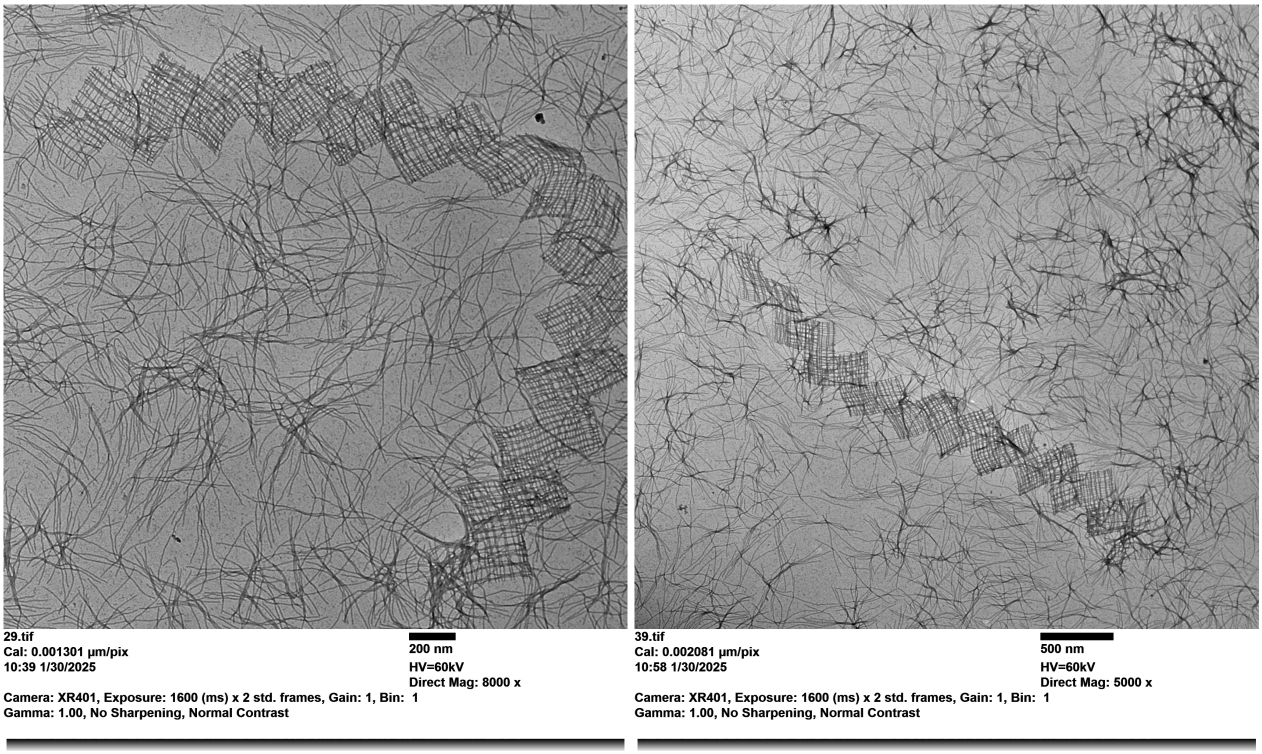

**Supp. Fig. 26**: Representative TEM images of the infinite ribbon megastructures used to compare the old and new handle libraries. The corresponding count data are shown in Figure 7 of the main text. Both images were taken from the ribbon sample using the new handle library, 15mM $MgCl_2$ and an 8 hour incubation time.
