## Supplementary material for "A computational framework for designing micron-scale crisscross DNA megastructures": seed_staple_generation_code.ipynb

```
# %% [code]
import pickle
import copy
import inspect
import random
import sys

# %% [code]
def comp_seq_FN(raw_sequence):
    uppercase = {'a':'a', 'A':'A', 'c':'c', 'C':'C', 'g':'g', 'G':'G', 't':'t', 'T':'T', '.':''}
    complement = {'a':'t', 'A':'T', 'c':'g', 'C':'G', 'g':'c', 'G':'C', 't':'a', 'T':'A', '.':''}
    antisense_seq = ''
    for letter in raw_sequence:
        if letter in uppercase:
            antisense_seq = complement[letter] + antisense_seq
    return antisense_seq

# %% [code]
# update filepaths to your system here
scaffold_file = '/Users/matt/Downloads/seed_design_package/p8064.txt'
cadnano_file = '/Users/matt/Downloads/seed_design_package/seed_design.json'

# %% [code]
# scaffold
input_file = open(scaffold_file, 'r')
maxiscaf_seq = input_file.read()
input_file.close()

# cadnano
input_file = open(cadnano_file, 'r')
all_file = eval(input_file.read())  # json unpacking
input_file.close()

second_hairpin_seq = 'GGGTGATGGTTCACGTAGTGGGCCATCGCCC'
maxiscaf_seq = comp_seq_FN(comp_seq_FN(maxiscaf_seq))  # reverse complement twice to remove any incorrect characters or carriage returns in the scaffold sequence
end_hairpins_position = maxiscaf_seq.find(second_hairpin_seq) + len(second_hairpin_seq)
maxiscaf_seq = maxiscaf_seq[end_hairpins_position:] + maxiscaf_seq[:end_hairpins_position]

pre_note = '241130_p8064_gridiron_seed_v0, ' 

vstrands = all_file['vstrands']
num_vstrands = len(vstrands)
num_bases = len(vstrands[0]['scaf'])
vsn = {}
for vstrand_num in range(num_vstrands):
    vsn[vstrands[vstrand_num]['num']] = vstrand_num
vsn[-1] = -1
null_bp = [-1, -1]

stap_color_dc  = {}

stap_color_dc[0]        = 'black'

stap_color_dc[4848714]  = 'dark gray'
stap_color_dc[4211199]  = 'most blue'
stap_color_dc[8421631]  = 'mid blue'
stap_color_dc[12631295] = 'less blue'

stap_color_dc[10197915] = 'light gray'

stap_color_dc[13632027] = 'pink red'
stap_color_dc[16098851] = 'vivid orange'
stap_color_dc[16312092] = 'bright yellow'
stap_color_dc[9131818] = 'earthy brown'
stap_color_dc[8311585] = 'lime green'
stap_color_dc[4289797] = 'dark spring green'
stap_color_dc[12390624] = 'magenta'
stap_color_dc[9442302] = 'purple'
stap_color_dc[4886754] = 'cyan-blue'
stap_color_dc[5301186] = 'green-cyan'
stap_color_dc[12118406] = 'soft green'
stap_color_dc[4868682] = 'muted teal'
stap_color_dc[4210943] = 'alt light blue'
stap_color_dc[12632319] = 'light lavender'

# %% [code]
# Generate arrays for stap paths and scaf paths

count_dc = {}

start_base_ra = [49, 266, 770]  # these are precisely selected to match the seed design staple positioning
for i in range(17):
    start_base_ra.append(53 + i*42)
    start_base_ra.append(94 + i*42)

for start_base in start_base_ra:
    count_dc[start_base] = 0

# this is basically collecting the scaffold and staple positions in the whole design.  The scaf is placed in its own variable, while all staples are placed in a list of lists (one list per staple)
scaf_path_ra = []
stap_path_ra = []
for vstrand_num in range(num_vstrands):
    for base_num in range(num_bases):
        for [parity, path_ra] in [['scaf', scaf_path_ra], ['stap', stap_path_ra]]:
            [prev_helix_num, prev_base_num, next_helix_num, next_base_num] = vstrands[vstrand_num][parity][base_num]
            if ([prev_helix_num, prev_base_num] == null_bp) and ([next_helix_num, next_base_num] != null_bp):
                sub_ra = []
                curr_helix_num, curr_base_num = vstrands[vstrand_num]['num'], base_num
                end_of_strand = False
                while not end_of_strand:
                    sub_ra.append([curr_helix_num, curr_base_num])
                    [curr_helix_num, curr_base_num] = vstrands[vsn[curr_helix_num]][parity][curr_base_num][2:]
                    end_of_strand = [curr_helix_num, curr_base_num] == null_bp
                if (parity == 'scaf') or (sub_ra[0][1] in start_base_ra):
                    path_ra.append(sub_ra)
                    if sub_ra[0][1] in start_base_ra:
                        count_dc[sub_ra[0][1]] += 1

print('number of scaf paths is', len(scaf_path_ra))
print('number of stap paths is', len(stap_path_ra))
    
for sub_ra in scaf_path_ra:
    print(len(sub_ra), sub_ra[0], sub_ra[1])


#Determine length of maxiscafs
maxiscaf_length = 0
for sub_ra in scaf_path_ra[:1]:
    for [helix_num, helix_base_num] in sub_ra:
        maxiscaf_length += 1 + vstrands[vsn[helix_num]]['skip'][helix_base_num] + vstrands[vsn[helix_num]]['loop'][helix_base_num]

print('maxiscaf length', maxiscaf_length)
seq_counter_dc = {maxiscaf_length:0, 1176:0}
seq_dc = {maxiscaf_length:[maxiscaf_seq]}

# Assign scaf base sequences
# This basically assigns the scaffold sequence where present in the design and an 'a' otherwise (so in the positions before and after the occupies helices in the design)
sub_ra = ['a' for i in range(num_bases)]
scaf_base_seq_dc = {}
for vstrand in vstrands:
    helix_num = vstrand['num']
    scaf_base_seq_dc[helix_num] = sub_ra[:]
for sub_ra in scaf_path_ra:
    print('scaffold path length is ', len(sub_ra))
    seq_length = 0
    for [helix_num, helix_base_num] in sub_ra:
        seq_length += 1 + vstrands[vsn[helix_num]]['skip'][helix_base_num] + vstrands[vsn[helix_num]]['loop'][helix_base_num]
    seq = seq_dc[seq_length][seq_counter_dc[seq_length]]
    seq_counter_dc[seq_length] += 1
    seq_pointer = 0
    for [helix_num, helix_base_num] in sub_ra:
        base_length = 1 + vstrands[vsn[helix_num]]['skip'][helix_base_num] + vstrands[vsn[helix_num]]['loop'][helix_base_num]
        scaf_base_seq_dc[helix_num][helix_base_num] = seq[seq_pointer:seq_pointer + base_length]
        seq_pointer += base_length

# this assigns the color for each staple in the entire design 
# colors are assigned only to the first base of the staple in a specific helix (this includes the 'start point' coming in from another helix)
meta_stap_color_dc = {}
for vstrand in vstrands:
    helix_num = vstrand['num']
    meta_stap_color_dc[helix_num] = [-1 for base_num in range(num_bases)]
    for [base_num, stap_color_int] in vstrand['stap_colors']:
        meta_stap_color_dc[helix_num][base_num] = stap_color_dc[stap_color_int]

# %% [code]
#Generate staple strand output

# this converts the staples to a simplified dictionary rep with the color, length and start/end positions and the sequence itself
stap_output_ra = []
for sub_ra in stap_path_ra:
    seq = ''
    for [helix_num, base_num] in sub_ra:
        seq += comp_seq_FN(scaf_base_seq_dc[helix_num][base_num])
    start_pointer = sub_ra[0]
    end_pointer = sub_ra[-1]
    [first_helix_num, first_base_num] = start_pointer
    stap_color = meta_stap_color_dc[first_helix_num][first_base_num]
    stap_output_ra.append([stap_color, len(seq), start_pointer, end_pointer, seq])

print('number of stap_output entries is', len(stap_output_ra))

#Sort staple strands according to caDNAno color
sorted_stap_output_ra = sorted(stap_output_ra, key = lambda stap_output:stap_output[0])

print()
print()

# these loops are manually assigning labels to different staples - edge, core and socket.  The socket ones are the ones that will be truncated to allow slats to attach.
edge_stap_output_ra = []
for sub_ra in stap_output_ra:
    if sub_ra[2][1] in [49, 266, 770]:
        edge_stap_output_ra.append(sub_ra)

core_stap_output_ra = []
for sub_ra in stap_output_ra:
    if (sub_ra[2][0] in range(17, 22)) and (sub_ra[2][1] in [53, 766]):
        core_stap_output_ra.append(sub_ra)    
    if (sub_ra[2][0] in range(17)) and (sub_ra[2][1] in range(53, 263)):
        core_stap_output_ra.append(sub_ra)
        
socket_stap_output_ra = []
for i in range(16):
    for sub_ra in stap_output_ra:
        if (sub_ra[2][0] in range(17, 22)) and (sub_ra[2][1] in [94 + i*42, 95 + i*42]):
            socket_stap_output_ra.append(sub_ra)

        
#Print edge staple strand sequences with annotations
tabber = '\t\t'
print('edge staple strands', len(edge_stap_output_ra))
for sub_ra in edge_stap_output_ra:
    seq = sub_ra[-1]
    note = pre_note + ' edge stap strand, ' + str(sub_ra[0]) + ', ' + str(sub_ra[1]) + 'mer, ' + str(sub_ra[2]) + ' start, ' + str(sub_ra[3]) + ' end'
    print(seq + tabber + note)
print()
print()


#Print non-toehold core staple strand sequences with annotations
print('core staple strands', len(core_stap_output_ra))
for sub_ra in core_stap_output_ra:
    seq = sub_ra[-1]
    note = pre_note + ' core stap strand, ' + str(sub_ra[0]) + ', ' + str(sub_ra[1]) + 'mer, ' + str(sub_ra[2]) + ' start, ' + str(sub_ra[3]) + ' end'
    print(seq + tabber + note)
print()
print()


#Print socket staple strand sequences with annotations
print('socket staple strands', len(socket_stap_output_ra))
for i in range(16):
    for sub_ra in socket_stap_output_ra[5*i:5*i + 5]:
        seq = sub_ra[-1][10:] # IMP: the first 10 bases of each socket staple are removed here (to be replaced by slat handles)
        # basically, 8 of the socket staples are extended with the seed toehold/poly-A sequence here.  They alternate throughout the entire length of the seed.
        if (sub_ra[2][0] == 17) and ((sub_ra[2][1] - 95)%84 == 0):
            seq += 'ttCACAACTACCAAAAAAAAAAAAAAAAAAAAAAAAA'
        note = pre_note + ' socket stap strand, ' + str(sub_ra[0]) + ', ' + str(len(seq)) + 'mer, ' + str(sub_ra[2]) + ' start, ' + str(sub_ra[3]) + ' end'
        print(seq + tabber + note)
    print()
print()
print()


#Print toeholds from neg-toehold core staple strand sequences with annotations
print('plugs from core staple strands', len(socket_stap_output_ra))
for i in range(16):
    for sub_ra in socket_stap_output_ra[5*i:5*i + 5]:
        seq = sub_ra[-1][:10] # IMP: the first 10 bases of each socket staple are removed here (to be replaced by slat handles)
        note = pre_note + ' plug from socket stap strand, ' + str(sub_ra[0]) + ', ' + str(len(seq)) + 'mer, ' + str(sub_ra[2]) + ' start, ' + str(sub_ra[3]) + ' end'
        print(seq + tabber + note)
    print()
print()
print()

# %% [code]
```
